# Achieving near single-molecule sensitivity in separation-free protein detection using engineered kinetic proofreading

**DOI:** 10.64898/2026.10.05.756571

**Authors:** Feng Xuan, Tsz Wing Fan, Xi Chen, Yu Wang, WonHee Kim, Wenjing Jiang, Jidong Tan, Xue Jiang, Yu Zhang, Pia Kivisakk, Bruno L. Hammerschlag, Oscar L. Lopez, Ann D. Cohen, Xuemei Zeng, Taraneh E. Atri, Alpana Singh, Nicholas J. Ashton, Steven E. Arnold, Thomas K. Karikari, Peng Yin

## Abstract

Unlike nucleic acids, proteins cannot be directly amplified. Ultrasensitive detection must therefore couple efficient analyte-to-signal conversion with strong suppression of target-independent background. Consistently detecting ~100 molecules (below which Poisson sampling noise alone causes CV to exceed 10%) across protein targets in a format as scalable and accessible as PCR remains unsolved. Separation-based ultrasensitive immunoassays require target binding stable enough to survive stringent washing and are therefore constrained by wash-induced signal loss and, more fundamentally, the relative scarcity of exceptionally tight binders. Performance consequently varies across targets; detection limits below 1,000 molecules remain rare. Conventional separation-free proximity assays avoid washing but incur background from random probe collisions, with reported detection limits frequently exceeding 100,000 molecules. Here we introduce Successive Proximity Extension Amplification Reaction (SPEAR), a separation-free architecture in which autonomous engineered DNA nanodevices emulate biological kinetic proofreading. Two successive irreversible proximity-extension steps compound discrimination between analyte-dependent and background reactions, suppressing spurious signals while preserving efficient analyte-to-signal conversion. This design reduces reliance on high-affinity binders and enables sub-fM detection from 1 µL samples through simple 3-step mixing followed by standard qPCR readout. Across 11 biomarkers tested, SPEAR consistently achieved sub-100-molecule detection limits. Where compared, it outperformed ultrasensitive separation-based platforms, analytically and clinically, including for low-abundance Alzheimer’s disease biomarkers. SPEAR also precisely quantified SARS-CoV-2 neutralizing antibody titers from finger-prick dried blood spots, concordant with the cell-based reference assay on venous blood. It detected a distinct mixed phosphorylation state of Tau, and targeting this state resolved previously observed capillary–venous discordance and improved amyloid-status discrimination. These results establish engineered autonomous successive molecular discrimination as a practical strategy for near-single-molecule protein detection, extending the sensitivity, scalability and accessibility of PCR-based testing to non-amplifiable analytes.

## INTRODUCTION

An ideal platform of biomolecule detection should feature high sensitivity – ideally at single molecule level, simplicity – ideally performed in homogeneous solutions, and accessibility – ideally without the need for complex and sophisticated instruments. One class of assay that embodies these attributes is real-time nucleic acid amplification tests (NAAT) such as qPCR. However, the success of such assays is largely predicated on the amplifiable nature of the nucleic acid analytes, an advantage that is not readily transferrable to other types of analytes such as proteins^(*1,2*)^. For non-amplifiable analytes, high sensitivity requires both efficient analyte-to-signal conversion and strong suppression of target-independent background. Conventional protein detection assays such as enzyme-linked immunosorbent assay (ELISA) typically exhibit limit of detection (LOD) in the pM to nM range. Considering the typical sample volumes of 10 to 100 µL in these assays, the LOD is equivalent to 10^7^ to 10^11^ molecules. In pursuit of higher sensitivity, one common approach is to employ stronger signal-amplification methods and/or innovative instrumentation, exemplified by immuno-PCR^(*3*)^ and digital ELISA (dELISA)^(*4*)^. However, these methods typically require separation of detection antibody molecules bound to the immobilized analyte from their unbound counterparts, often through lengthy and stringent wash steps to minimize the non-specific binding of the detection antibody to the capture surface. Unfortunately, the wash steps also cause true signal to diminish due to dissociation of the capture or the detection antibody from the analyte, reducing assay sensitivity^(*5*)^. For example, in many dELISA assays where high sensitivity is prioritized, 60 to 90 minutes of operation is needed (including multiple washes and sequential addition of detection antibodies and signal-generation enzymes) after the completion of the capture of the analyte by the surface-attached capture antibodies. Simple kinetic modeling shows that during these long operations, even if both antibodies have *K*_D_ as low as 200 pM, and dissociation rate constant (*k*_off_) as low as 1×10^−4^ s^−1^, only 14.4% of the analyte molecules initially bound by the capture antibodies are converted to signal-generating molecules. If the *K*_D_ is 1 nM and *k*_off_ is 5×10^−4^ s^−1^, which are in the common range of the affinities for research-use antibodies, this conversion rate drops to only 0.78% (**Supplementary Fig. S1**). The need to retain both capture and detection binders through prolonged and stringent washing therefore demands exceptionally tight binder pairs with slow dissociation rates. Such binder pairs remain relatively scarce, posing a fundamental constraint on achieving consistently ultrasensitive detection across targets with separation-based assays. In practice, LODs vary widely: among 62 commercially available dELISA assays on the Simoa platform^(*6*)^, the 25^th^, 50^th^ and 75^th^ percentile LODs are ~16,000, ~60,000 and ~250,000 molecules, respectively (compiled from manufacturer data sheets; **Supplementary Table S1**).

Separation-free immunoassays such as competition-^(*7–9*)^ and proximity^(*10,11*)^-based immunoassays where unbound antibodies do not need to be removed may be less impacted by these washing-related limitations and enjoy simpler workflows, but often suffer from even lower sensitivity, because even in the absence of the target protein, the condition for signal generation still exists. For example, in proximity ligation and proximity extension assays, the two antibody-conjugated probes may still encounter in the solution in an analyte-independent manner and react to form background amplicons. In these methods, the underlying mechanism of signal generation (extension or ligation) is the same for the true and background amplicons, and the ability to discern the two originates from the fact that the effective local concentration of the two probes in the tripartite immunocomplex (formed by the two antibodies and the analyte) is higher than their effective global concentration in the solution. Thus, the ratio between these two concentrations is the key determinant of sensitivity. Unfortunately, this ratio (which we refer to as the ‘characteristic ratio’, or *R*_C_) cannot be arbitrarily high, as the local concentration cannot exceed what is afforded by the molecular geometry of the tripartite immunocomplex, and the global concentration cannot be lower than what is necessary to achieve efficient binding (which is usually around the *K*_D_ of the antibody, see **Supplementary Text 2.1.1–2.1.4**). These boundaries ultimately limit the sensitivity of the proximity-based approach. In practice, a report of 90 separation-free PEA assays showed that the LODs at the 25^th^, 50^th^ and 75^th^ percentiles are ~160,000, ~1,000,000 and ~7,700,000 molecules, respectively (**Supplementary Table S2**)^(*12*)^.

Recently, proximity-based signal generation has been combined with dual capture-and-wash steps using orthogonal affinity handles, leading to lower background than separation-free PEA^(*13*)^ but reintroducing the surface-and separation-related vulnerabilities discussed above. As a prominent example of this strategy, NUcleic acid Linked Immuno-Sandwich Assay (NULISA) improves sensitivity to LODs at the 25^th^, 50^th^ and 75^th^ percentiles of ~1,600, ~8,000 and ~59,000 molecules, respectively, excluding assays intentionally desensitized via hot/cold antibody tuning (105 of 183 assays retained) (**Supplementary Table S3**)^(*13*)^.

To fundamentally improve sensitivity of proximity-based immunoassays in separation-free format, one must ‘break’ the limitation posed by the aforementioned characteristic ratio. This challenge resembles those in many biological processes where exquisitely high substrate specificity is necessary. For instance, in DNA replication, the ratios of the affinities of the correct versus incorrect nucleotide for the active site of the extending DNA polymerase are usually 10-to 1,000-fold. Yet, the ratio of correct versus incorrect nucleotide incorporation can be greater than 10^6^-fold even in the absence of post-synthesis DNA repair^(*14*)^. Protein translation also happens with much higher accuracy (error rate of ~1 in 3,000) than what the difference in affinities between correct and incorrect codon-anticodon pair (~100-fold) affords^(*15*)^. Such high substrate specificity was famously posited to be achievable through kinetic proofreading^(*16,17*)^, where the accuracy can be greatly enhanced through multiple energy-consuming (e.g., ATP-or GTP-hydrolyzing), irreversible steps, each of which serves as a checkpoint for the correct substrate and reads the same affinity difference independently, so that the discriminations compound. Although the exact mechanisms were later discovered to involve additional concepts such as induced fit^(*18,19*)^, kinetic proofreading stands as a key framework to understand substrate specificity in biology. One form of kinetic proofreading begins with breaking a reaction into two successive irreversible steps, i.e., by first forming an intermediate and then forming the final product. For example, in protein translation, the EF-Tu·GTP·aminoacyl-tRNA ternary complex first engages the ribosomal A site, hydrolyzes the GTP, releases EF-Tu·GDP and leaves the aminoacyl-tRNA held in the A site by its codon-anticodon interaction (the intermediate). Next, this intermediate is accommodated into the peptidyl transferase center to receive the growing peptide, forming the final product. Because the affinity of the codon-anticodon pair governs the rate of reaction or success rate of both steps^(*15,19*)^, the difference in affinity is compounded to achieve much higher substrate specificity.

Here, we present Successive Proximity Extension Amplification Reaction (SPEAR, **Fig. 1**), a separation-free assay that uses autonomous engineered DNA nanodevices to emulate biological kinetic proofreading and distinguish analyte-dependent from background signals. Like its biological counterparts, in SPEAR, a primer is first extended on a template to form an intermediate (i.e., the first energy-consuming, irreversible step), spontaneously dissociates from its template, and is then extended on a different template for a second time (i.e., the second energy-consuming, irreversible step) to form the final amplicon. Because the primer must dissociate and rebind between the two irreversible extensions, each step discriminates independently. To apply SPEAR in protein detection, the primer is attached to one antibody for the analyte, and both templates are attached to the other antibody for the analyte. Thus, both extension reactions are sped up in a proximity-dependent fashion when the two antibodies bind to the same analyte molecule. Because of the two-step kinetic-proofreading process, the rate ratio of proximity-dependent to background amplicon formation can approach the square of the characteristic ratio, and as a result, greatly reduce background and enhance sensitivity. The final amplicon can be easily and accurately quantified using standard qPCR. Kinetic modeling predicts that this design improves sensitivity by nearly 10,000-fold (**Supplementary Text 2.1.5–2.1.7**) over conventional homogeneous proximity-dependent amplicon-generation assays (PAG, e.g., PLA and PEA) and can approach near single-molecule sensitivity even when the affinities of the antibodies are suboptimal.

**Figure 1.**
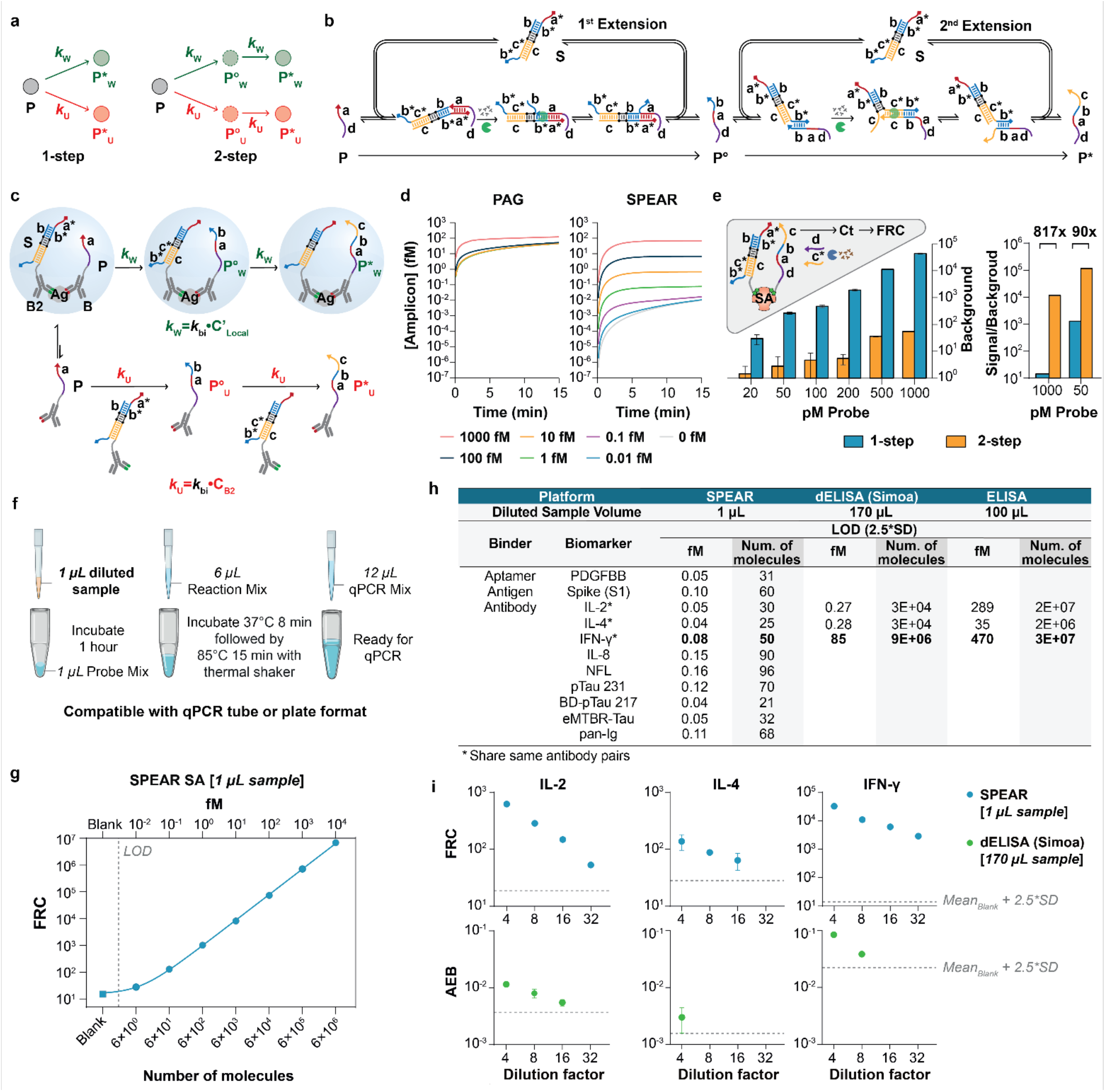
SPEAR: 2-step kinetic-proofreading assay for near single-molecule protein detection. **a**, Model of 1-step and 2-step kinetic-proofreading process. The precursor (**P**) is converted to the wanted product (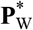) and the unwanted product (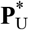) directly (1-step) or through intermediates (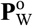, 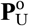 2-step) using rate constants ***k***_W_ and ***k***_U_. **b**, Design of a 2-step extension cascade: Primer (P) undergoes two successive extensions to form P^0^ and P^∗^, using probe S and polymerase with strand-displacement activity. **c**, Integration of a 2-step extension cascade into a proximity-based protein assay: When binders (B and B2), for example, antibodies, bind to the target protein, colocalization accelerates the formation of target-dependent products 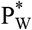, while target-independent interactions create background 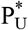. **d**, Simulation of proximity-based amplicon-generation assay (PAG, using **Eqs. 2.21** and **2.10.6** in **Supplementary Information**) and 2-step kinetic-proofreading DNA extension reactions (using **Eqs. 2.18** and **2.22.1**) showing substantial background suppression, improving signal differentiation at ultra-low target concentrations (0.01 fM, see **Supplementary Text 2.1.7** for models, parameters: *k*_uni_ = 0.01 s−^1^, *k*_bi_ = 5,000 M−^1^s−^1^, *k*_on_ = *k*_on2_ = 5×10^5^ M−^1^s−^1^, *K*_D_ = *K*_D2_ = 1 nM, *C*_B_ = *C*_B2_ = 100 pM in the 8 µL extension step, i.e., 400 pM in the 2 µL binding step, *t* = 0 to 15 min; target and amplicon concentrations are those in the 8 µL extension step; total amplicon = *S*_True_ + *S*_BG_). **e**, Comparison of background (left) and signal-to-background ratio (right, at 1 pM Streptavidin (SA)) for 1-step and 2-step kinetic-proofreading assay across varying probe concentrations (20-1,000 pM) based on qPCR readout. Signal and background are presented as Full Record Count (FRC) calculated from qPCR reaction. **f**, Workflow of a separation-free SPEAR assay for SA detection. **g**, Performance of SPEAR SA assay, showing detection limit down to single-digit count of molecules. Grey dashed line: LOD. **h**, Versatility of SPEAR in binder selection (aptamer, antigen and antibody) with consistent attomolar sensitivity across 11 different biomarkers, including comparative sensitivity analysis of SPEAR with dELISA (digital ELISA, Simoa HD-X analyzer), and conventional ELISA (BD OptEIA™ kits) for low-abundance human cytokines (IL-2, IL-4, and IFN-γ) using diluted sample volumes of 1 µL, 170 µL, and 100 µL, respectively. Limits of Detection (LOD) are calculated as mean of blank + 2.5×(SD of blank) and reported in both concentration and number of molecules. **i**, Quantitative comparison of SPEAR and dELISA measurements on the same serially diluted serum samples for IL-2, IL-4, and IFN-γ from 4-to 32-fold dilution. Blue: SPEAR; Green: dELISA; Grey dashed line: LOD; AEB: Average Enzymes per Bead.

Consistent with the model, the SPEAR assays we developed for 11 protein biomarkers, many of which have low abundance, all achieved LODs below 100 molecules. In matched analytical and clinical comparisons, SPEAR also improved detection or quantification over dELISA and NULISA for the assayed targets.

Notably, SPEAR effectively detected plasma eMTBR-Tau^(*20*)^, an emerging low-abundance biomarker that is being explored as a Tau-PET alternative and as a tool for patient staging and selection for amyloid-or Tau-targeting therapies^(*21*)^. With a simple workflow comprising three mixing steps followed by standard qPCR readout, SPEAR is well suited for applications requiring both high sensitivity and broad accessibility. We thus applied SPEAR in two such scenarios: immune surveillance for SARS-CoV-2 infection or vaccination, and blood-based assessment for aging-related neurodegenerative diseases. Toward the former, we developed dried blood spot (DBS) eluate-based assays that not only detect Spike proteins and anti-Spike antibodies but also quantify neutralization ability of serum/plasma antibodies. Toward the latter, we leveraged the sensitivity of SPEAR to detect a distinct mixed phosphorylation state of Tau using an antibody pair targeting phosphorylated T217 and an upstream epitope containing unphosphorylated S191, S195 and S199. A SPEAR assay targeting this state resolved the capillary-venous measurement discordance seen with conventional pTau 217 assays in a small cohort, addressing a key analytical barrier to decentralized testing. Additionally, this assay (which we call the ‘epTau 217 assay’) showed greater discrimination between amyloid PET-positive and PET-negative subjects than conventional pTau 217 assays. It also distinguished amyloid-positive from amyloid-negative subjects in a separate cohort classified by CSF testing.

## RESULTS

### SPEAR as a 2-step kinetic proofreading DNA circuit

A simple form of 2-step kinetic proofreading is shown in **Fig. 1a**. Consider a precursor molecule (**P**) that may be converted to a wanted product (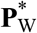) and an unwanted product (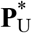) via reactions having rate constants ***k***_W_ and ***k***_U_, respectively (***k***_W_ > ***k***_U_, **Fig. 1a**, left). We refer to the ratio ***k***_W_/***k***_U_ as the characteristic ratio in this system, written as *R*_C_. It follows that at any time point, the ratio of the concentrations of 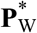 and 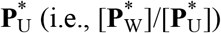 cannot exceed *R*_C_ (**Supplementary Text 2.1.5**). However, if **P** is first converted to wanted and unwanted intermediates 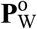 and 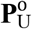 with rate constants ***k*** and ***k*** respectively, and then 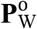 and 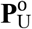 are in turn converted to products 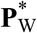 and 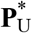 with rate constants ***k*** and ***k***_U_, respectively (**Fig. 1a**, right), the ratio 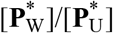 can exceed *R*_C_ and approach 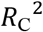 (**Supplementary Text 2.1.5**).

To recapitulate this process for proximity-based protein detection, we designed SPEAR (**Fig. 1b, Supplementary Movie 1**), where the primer P (having domains **d** and **a**) can be extended to form P^0^ (having domains **d, a** and **b**) in a toehold-mediated isothermal and autonomous reaction known as PER (Primer Exchange Reaction)^(*22*)^, using an oligonucleotide-based construct S as the template and DNA polymerase with strand-displacement activity (e.g., Bst). Specifically, the ~8-nt **a** domain of the P molecule can reversibly bind to the **a\*** domain of S. Upon binding, the **a** domain of P can be extended by the strand-displacement DNA polymerase while displacing the **b** domain of S. Importantly, the **b** domain consists of only A, T and C bases, and the reaction buffer only contains dATP, dTTP and dCTP. The nucleotide adjacent to the **b** domain on S (highlighted by a black dot) is a 2’-O-methylguanosine and serves as an extension stopper. After extension, the polymerase may dissociate from the DNA complex allowing the **b** domain on S and the newly synthesized **b** domain on P to exchange their position on the opposite **b\*** domain via random 3-way branch migration. When the branch migrates to (or near) the 5’ of the **b** domain on S, P may spontaneously dissociate from S. P^0^ can be extended to form P^∗^ (having domains **d, a, b** and **c**) in a similar reaction, using another part of S as the template. P^0^ and P^∗^ are also referred to as half record and full record, respectively. To detect an analyte, P and S are attached to binders (e.g., antibodies) B and B2, respectively. B and B2 can bind two different epitopes of the same target protein (**Fig. 1c**), forming a tripartite immunocomplex. After mixing the probes with the analyte for a reasonable period (e.g., 1 hr or overnight), Bst is added to the reaction. Within the immunocomplex, P and S are co-localized and both the P→P^0^ and P^0^→P^∗^ extensions are accelerated due to the proximity, with the higher first-order rate constant *k*_W_ (**Fig. 1c**, top), compared to the random diffusion-and-collision based process for P and S molecules that are not bound to the same analyte molecule, with the lower pseudo-first-order rate constant *k*_U_ (with respect to B-P and treating the concentration of B2-S as a constant, **Fig. 1c**, bottom). We refer to the fraction of P that is present in the tripartite immunocomplex as P_W_, and its extension products as 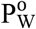 and 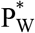 since these extension products represent the true, analyte-dependent signal and are ‘wanted’. Similarly, the extension products of P attached to free (not target-bound) binder B are referred to as 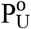 and 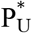since these extension products represent the background signal and are ‘unwanted’.

Mathematical analysis in **Supplementary Text 2.1.5–2.1.6** shows how this system recapitulates the kinetic proofreading process shown in **Fig. 1a** and is predicted to improve sensitivity by nearly 10,000-fold over simple proximity-based amplicon-generation assays. For a quick preview, consider the simple situation where the concentration of B-P conjugate (i.e., the probe) far exceeds its *K*_D_ to the target protein, in which case the concentration of P_W_ is approximately that of the analyte. We further consider a realistic situation where the analyte concentration is 1 fM and the probe concentration is 1 nM. In other words, [P] is a million-fold higher than [P_W_]. Even if the analyte-dependent extension 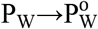 happens 1,000 times faster than the free extension 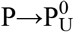 (i.e., *k*_W_ = 1,000*k*, or *R*_C_ = 1,000), the concentration of analyte-independent intermediate 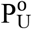 (leading to background signal) would still be 1,000 times higher than that of analyte-dependent 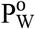. However, by introducing the second extension, we can accelerate the formation of target-dependent full record 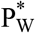 over that of 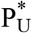 by a factor of *R*_C_, making the true signal more readily discernible from the background. Simulations using a hypothetical model and a set of realistic parameters (using the same binder pair each having 1 nM *K*_D_, see **Supplementary Text 2.1.7** for details) show that while the total signal (true plus background signal) can only be discerned from background signal once the analyte concentration exceeds 100 fM for a conventional proximity-based amplicon-generation assay, SPEAR can suppress the background by >4 orders of magnitude and allow signal generated by as little as 0.01 fM analyte to be readily detected above background (**Fig. 1d**).

To experimentally confirm the feasibility of SPEAR, we modified P and S with biotin and used streptavidin (SA) to induce their colocalization. As shown in **Supplementary Fig. S2a** (top), even in the absence of streptavidin, P^0^ began to appear 1 min after incubation. However, the twice-extended P^∗^ was undetectable until the 30 min time point, demonstrating the suppression of background. In the presence of streptavidin, the concentration of P^∗^ nearly plateaued at the earliest time point (1 min), showing that the 2-step process did not markedly slow down the generation of signal. When yield (defined as the fraction of P that is extended) is quantified, it can be seen that the ratio of yields with and without SA is substantially improved with the 2-step kinetic proofreading (**Supplementary Fig. S2a**, bottom right compared to bottom left). We optimized the toehold lengths and sequence of P and S and suppressed the production of P^∗^ to an undetectable level (using fluorescent labeling) without streptavidin, while its production in the presence of streptavidin was largely unaffected. We next used the more sensitive and quantitative qPCR as the readout (triangular inset of **Fig. 1e, Supplementary Fig. S2b**-bottom) and designed a control set of P and S probes (**Supplementary Fig. S2b**-top), where the **b** domain was replaced by the **c** domain, and the portion of the S supporting the second extension was omitted. These control probes allow us to detect the background of a simple proximity-dependent amplicon-generation assay. We observed 90-and 817-fold enhancements of signal-to-background ratio when the probe concentrations were 50 pM and 1 nM, respectively (**Fig. 1e**, right). The near 3-orders-of-magnitude enhancement of signal-to-background ratio at high probe concentration is critical to its application in immunoassays as it allows the probe to be used at a higher concentration, converting a higher fraction of analyte to amplicon, and ensuring timely completion of binding. Consequently, we were able to develop a simple workflow (**Fig. 1f**) requiring only 1 µL of 4× diluted sample, 1 µL of Probe Mix, 1 hour of antibody binding, 2 reagent addition steps (6 µL of Reaction Mix and 12 µL of qPCR Mix) and thermocycling to conduct SPEAR-based immunoassays. The wash-free nature of this SPEAR workflow also minimizes signal loss, since the amplicons are generated within minutes after the addition of the Reaction Mix (containing the DNA polymerase), during which a substantial fraction of the analyte molecules captured in the sandwich structure is converted to the amplicon. Kinetic modeling shows that, if both sandwiching antibodies have *K*_D_ of 200 pM and *k*_off_ of 1×10^−4^ s^−1^, this conversion rate is as high as 62.1%, compared to 14.4% for dELISA. More importantly, if the *K*_D_ is 1 nM and *k*_off_ is 5×10^−4^ s^−1^, typical for research-use antibodies, this conversion rate can still reach 20.2%, compared to 0.78% for dELISA (**Supplementary Fig. S1b**).

To experimentally validate this workflow, we first used the mixture of biotinylated P and S as a model biosensor, SA as a model analyte, and the copy number of full record (or full record count, FRC, as determined by qPCR) as the readout. As shown in **Fig. 1g**, in the absence of streptavidin, merely 15±1.6 molecules of full record were generated (see ‘Blank’). This low background enables LOD near single molecule level. Indeed, as little as 6 molecules of streptavidin generated signal (FRC) higher than the background with statistical significance. High linearity between analyte concentration and FRC across 6 orders of magnitude was observed.

### Near single-molecule sensitivity across 11 biomarkers

We next built SPEAR assays for 11 protein biomarkers (**Fig. 1h**) using antibodies, aptamers and antigens as the binders. These analytes include inflammatory cytokines (IL-2, IL-4, IL-8 and IFN-γ), a growth factor (PDGFBB), infectious disease markers (SARS-CoV-2 Spike protein S1, SARS-CoV-2 Pan-Ig), and neurology markers (neurofilament light chain (NFL), pTau 231, brain-derived pTau 217 (BD-pTau 217), and eMTBR-Tau), many of which require LOD at sub-to low-fM to be clinically useful when peripheral blood or eluted DBS are used as samples. All these SPEAR assays demonstrate LOD of 20 to 100 molecules (or 0.03 to 0.16 fM in 1 µL, **Supplementary Fig. S3**). We carried out head-to-head comparisons, using the same antibody pairs, with dELISA (on the commercially available Simoa platform) on 3 low-abundance inflammatory biomarkers (**Supplementary Fig. S4**). We found that although dELISA assays for IL-2 and IL-4 showed sensitivity in the typical range for commercial dELISA assays (**Supplementary Table S1**), with LOD around 3×10^4^ molecules, the sensitivity for IFN-γ was rather poor, with LOD about 9 ×10^6^ molecules, possibly due to lower antibody affinity. In contrast, SPEAR assays for all three analytes showed LOD at 25 to 50 molecules. These results are consistent with the notion that for existing immunoassays, optimal sensitivity requires antibodies with extremely low *K*_D_ (ideally in the pM range)^(*5*)^, whereas the quantitative model of SPEAR predicts that, thanks to its kinetic-proofreading nature, LOD below 100 molecules may be achieved even when the *K*_D_ is in the nM range (**Supplementary Text 2.1.6** and **Supplementary Fig. S5)**.

The superior sensitivity was maintained when clinical samples were used. When we challenged SPEAR and dELISA to detect low-abundance cytokines such as IL-2, IL-4 and IFN-γ in serially diluted serum samples and determined the highest level of dilution that generated signals above background, SPEAR consistently withstood at least 2-to 4-fold greater dilutions than dELISA while using 170-fold less sample volume (**Fig. 1i**).

To evaluate the suitability of SPEAR assays for neurological disorders, in which many clinically informative biomarkers exist at very low abundance, we first built an assay to detect NFL. This assay showed LOD and LLOQ of 0.16 fM and 1.84 fM, respectively, using 1 µL of 4-fold diluted sample (effectively 0.25 µL of plasma sample) per replicate, and minimal cross reactivity to neurofilament medium (NFM) and heavy (NFH) chains (differentiation by 5 to 6 orders of magnitude, **Supplementary Fig. S3d**). In healthy donors, 100% of EDTA plasma, heparin plasma, serum and CSF samples were quantifiable with readings well above functional LLOQ (fLLOQ, **Fig. 2a**, left). Importantly, the SPEAR assay exhibited high intra-plate precision (3.2% mean CV, **Fig. 2a**, middle), likely benefiting from its homogeneous and separation-free nature. The results of the SPEAR assay also showed high correlation with a well-established, commercial dELISA assay (Spearman ρ = 0.996, **Fig. 2a**, right).

**Figure 2.**
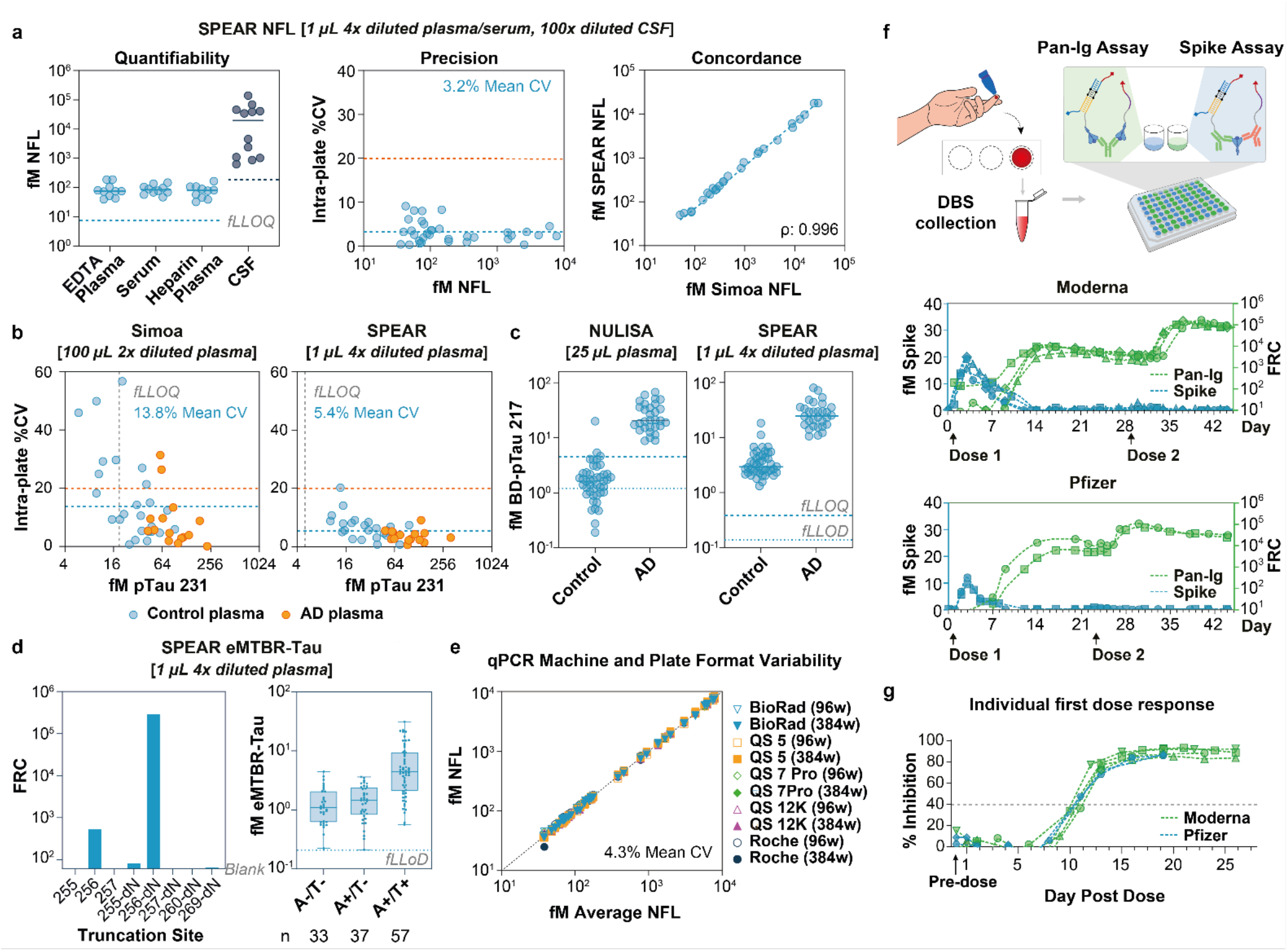
Detection of ultralow-abundance biomarkers using SPEAR assays. **a**, Neurofilament Light (NFL) assay performance. Left: Quantifiability in healthy EDTA plasma, serum, lithium heparin plasma, and CSF using 1 µL 4× diluted blood sample or 100× diluted CSF sample (*n* = 10, 10, 10, 12). Grey dashed lines indicate fLLOQ (NFL: 7.4 fM for plasma/serum, 184 fM for CSF). Bars represent median. Middle: Intra-plate precision for EDTA plasma and serum (*n* = 10, 24). Blue dashed line: mean %CV; orange dashed line: 20% CV threshold. Right: Concordance between SPEAR and Simoa assays for NFL (EDTA plasma: *n* = 10; serum: *n* = 24). Blue dashed line: linear regression fit. **b**, Comparison of intra-plate precision and quantifiability between Simoa and SPEAR pTau 231 assays with 100 µL of 2× diluted plasma and 1 µL of 4× diluted plasma from healthy controls (*n* = 25, blue) and Alzheimer’s disease (AD) patients (*n* =15, orange), respectively. Grey dashed line: fLLOQ (Simoa: 18.6 fM, SPEAR: 5 fM); Blue dashed line: mean % CV; orange dashed line: 20% CV threshold. **c**, Comparison of quantifiability between NULISA (sample: 25 µL plasma) and SPEAR (sample: 1 µL 4× diluted plasma) BD-pTau 217 assays. Thin and thick dashed lines indicate fLLOD (NULISA: 1.21 fM, SPEAR: 0.14 fM) and fLLOQ (NULISA: 4.54 fM, SPEAR: 0.38 fM), respectively. N.D.: Not detected. **d**, Performance of SPEAR eMTBR-Tau assays. Left: SPEAR readout of 10 pM of different peptides. Right: eMTBR-Tau values measured by the SPEAR assay in 33 A−/T−, 37 A+/T− and 57 A+/T+ subjects. **e**, Robustness of SPEAR NFL sample measurement (*n* = 39) across qPCR machines and plate format (96-well or 384-well). Mean %CV of the measurements is indicated. Dotted line: line of identity. **f**, Top: schematic of parallel measurement of anti-SARS-CoV-2 Pan-Ig and Spike assay from finger-stick DBS samples. Middle: longitudinal monitoring of anti-SARS-CoV-2 Pan-Ig (green) and Spike protein (blue) in COVID-19 vaccine recipients (Moderna: top, *n* = 4; Pfizer: bottom, *n* = 2) following first and second doses. Each assay used 1 µL of DBS eluate. The *x*-axis indicates days post-vaccination; arrows mark dose administration. **g**, Longitudinal monitoring of neutralizing antibody inhibition (%) in Moderna (*n* = 4, green) and Pfizer (*n* = 2, blue) recipients after the first dose using DBS samples and SPEAR neutralization assay.

Encouraged by these results, we challenged SPEAR to detect pTau 231 and BD-pTau 217, two AD-related biomarkers with even lower abundance, again using only 1 µL of 4× diluted plasma sample per replicate. A commercial electrochemiluminescence immunoassay can only detect pTau 231 in 7 to 14% of plasma samples from non-AD donors^(*23*)^. The more sensitive dELISA assay also failed to detect and quantify pTau 231 in 22% and 60% of non-AD donors^(*24*)^, respectively. The SPEAR pTau 231 assay showed LOD and LLOQ of 0.12 fM and 1.25 fM, respectively, as well as minimal cross reactivity to non-phosphorylated Tau (np-Tau) and other pTau analytes such as pTau 181 and pTau 217 (**Supplementary Fig. S3e**). This assay successfully quantified plasma pTau 231 in 100% (40 out of 40) of donors, 25 non-AD and 15 AD (**Fig. 2b**). As was the case for NFL, the SPEAR pTau 231 assay also showed high quantifiability and precision. Compared to commercial dELISA-based pTau 231 assay (using 100 µL of 2× diluted sample), which showed mean CV of 13.8% with 10 out of 40 samples exhibiting CV greater than 20%, the SPEAR-based assay showed mean CV of 5.4% with no sample exhibiting CV greater than 20% (**Fig. 2b**). Likewise, the SPEAR BD-pTau 217 assay, which targets phosphorylated T217 and the exon 4-5 junction with its two antibodies, showed LOD and LLOQ of 0.035 fM and 0.095 fM (**Supplementary Fig. S3f**), respectively, and was able to detect and quantify plasma BD-pTau 217 in 100% of controls (44 out of 44) and AD samples (30 out of 30). In comparison, the NULISA assay (using 25 µL sample) was only able to detect 68% (30 out of 44) and quantify 7% (3 out of 44) of controls, although both assays showed substantially elevated levels of BD-pTau 217 in AD samples (**Fig. 2c**).

To further challenge the sensitivity of SPEAR, we asked whether it could detect eMTBR-Tau, an emerging biomarker whose plasma level has been reported to reflect severity of Tau pathology in symptomatic AD patients^(*20,21*)^, from 1 µL of 4× diluted plasma samples. The eMTBR-Tau biomarker is characterized by endogenous cleavage immediately downstream of V256, and a fraction of eMTBR-Tau also contains naturally deamidated N255, converting it to D255. Using a peptide having the residues 220 to 256 of Tau with deamidated N255 (peptide ‘256-dN’) as the calibrator, the SPEAR eMTBR-Tau assay showed LOD of 0.05 fM (**Supplementary Fig. S3g**), and was able to detect plasma eMTBR-Tau from 98.6% (69 out of 70) of Tau PET-negative samples. The assay proved to be highly selective to the cleavage downstream of V256, as it generates 11,000-fold lower signal if residue 257 is included in the peptide (peptide ‘257-dN’), or 7,300-fold if residues 257–260 (peptide ‘260-dN’) or 4,400-fold if residues 257–269 (peptide ‘269-dN’) are included (**Fig. 2d**, left). It also displayed strong preference for the peptide having deamidated N255, as peptide ‘256’ (without deamidation at N255) generated ~550-fold lower signal than 256-dN. In a cohort of 127 subjects with different radiologically confirmed amyloid and Tau PET status, this assay detected substantially higher (mean value 4.54-fold higher) levels of eMTBR-Tau in the amyloid PET-positive and Tau PET-positive (A+/T+) subjects, than in A+/T− and A−/T− subjects, consistent with the notion that eMTBR-Tau is an effective biomarker for Tau pathology.

### Immune surveillance from finger-prick dried blood spots

Because SPEAR reports its output as DNA amplicons quantified by qPCR, it can in principle run on widely available qPCR instrumentation. Toward this end, we first assessed whether a SPEAR assay can generate the same absolute quantification results across multiple qPCR platforms. We therefore ran the NFL assay using a series of control samples on five different qPCR instrument models in both 96- and 384-well formats, and observed consistent results (mean CV 4.3%, **Fig. 2e**). With this established, we set out to apply SPEAR to immune surveillance for infectious diseases.

Although active infection of most infectious diseases can be readily detected with NAAT, long-term monitoring of immune response, which informs risk management and vaccine recommendation, remains difficult to perform outside centralized laboratories. Taking COVID-19 as an example, immune surveillance typically requires large volumes of venous samples and, in the case of the gold-standard neutralization assays, live cell-based reagents, both of which limit decentralized sampling or testing. We thus asked whether immune surveillance could instead be performed from finger-prick DBS. We focused on three aspects of immune surveillance: whether antigen protein generated by vaccination can be detected, whether antibody response against the antigen can be measured, and whether the neutralizing activity of the response can be quantified. We first developed SPEAR assays to detect SARS-CoV-2 Spike protein (the Spike assay) and to detect pan-immunoglobulins against the Spike protein (the Pan-Ig assay, **Fig. 2f**, top). For the Pan-Ig assay, the binders for both the P and S probes are the Spike protein, and the monoclonal anti-Spike antibody clone R2B17 was used as the model antibody (see **Supplementary Fig. S6a** for data of other clones). Both assays showed high analytical sensitivities, with LOD of 0.10 fM and 0.11 fM, respectively (**Fig. 1h**). In subjects receiving mRNA vaccines for COVID-19, when 1 µL of DBS eluate was used as samples, the Spike protein translated from the mRNA vaccine was readily detectable in the DBS eluate after the first dose (**Fig. 2f**, middle, blue), consistent with plasma measurements in mRNA-1273 recipients^(*25*)^. The Spike protein was not observed after the second dose, likely because the vaccine-induced serum anti-Spike antibody masked or cleared the serum Spike proteins. We observed robust and quantitative readouts of anti-Spike antibody (Pan-Ig, **Fig. 2f**, middle, Moderna; bottom, Pfizer; green) as early as 7 days after the first dose, and >10-fold boost 4 to 6 days after the second dose.

We next evaluated whether the more challenging neutralization assay using DBS eluate is feasible. We designed a signal-off molecular neutralization assay using SPEAR (**Supplementary Fig. S6b**) where the Spike protein S1 subunit and ACE2 are conjugated to the SPEAR S and P probes, respectively. In the absence of a neutralizing antibody, the interaction between Spike and ACE2 colocalizes the two probes and promotes the generation of the full record. Neutralizing antibodies block this process, resulting in a signal-off assay. When tested against six commercially available monoclonal anti-Spike antibodies, this assay clearly distinguished neutralizing from non-neutralizing clones (**Supplementary Fig. S6c**). Moreover, using serum samples from SARS-CoV-2 vaccine recipients and convalescent patients recovered from non-SARS-CoV-2 viral infections, the assay only generated neutralization signal with SARS-CoV-2 vaccine recipients, demonstrating high specificity (**Supplementary Fig. S6d**). In molecular neutralization assays using similar signal-off principles, the Spike- and ACE2-mimicking probes must be used at sufficiently high concentration to produce strong signal so that its inhibition can be accurately quantified. When using DBS eluates, the amounts of serum antibodies in the sample may not be high enough to inhibit a sufficient fraction of such probes simply due to low stoichiometry, leading to ambiguous results. Because of the high sensitivity of SPEAR, as low as 1 pM of these probes resulted in signals several orders of magnitude higher than background. Consequently, this SPEAR assay should be able to generate quantitative results with much lower antibody input. Indeed, in head-to-head comparisons it showed 2-orders-of-magnitude higher sensitivity compared to CPass, an ELISA-based surrogate virus neutralization assay^(*26*)^ (**Supplementary Fig. S6e**). From the DBS eluates of subjects receiving the Moderna and Pfizer mRNA vaccine, the SPEAR assay detected neutralizing antibodies as early as 10 to 12 days after receiving the first dose of either vaccine (**Fig. 2g**).

Using DBS eluates from another cohort of subjects, conventional CPass and PsVNA (a pseudotyped virus neutralization assay^(*27*)^) assays showed substantial overlap among results in the COVID-naïve, infected, and vaccinated groups, while SPEAR showed clear separation between the COVID-naïve group and the other two groups (**Supplementary Fig. S6f**). In serially diluted samples, the NT50 values (i.e., dilution fold that yielded 50% of the maximum inhibition activity compared to the undiluted sample) between the SPEAR assay using DBS samples, and the gold-standard PRNT (plaque reduction neutralization test) using large volumes of serum sample were highly concordant (**Supplementary Fig. S6g**). Longitudinal tracking of the NT50 values from two subjects who experienced several events of vaccination and infection showed rising and waning of neutralization activity against both the wildtype and the Omicron BA.1 variant of SARS-CoV-2 (**Supplementary Fig. S6h**). Data from a limited number of subjects suggest that neutralizing activity against both wildtype and Omicron BA.1 strains in subjects receiving the booster dose appeared to have lower titer (**Supplementary Fig. S6i**, middle compared to right) and wane faster (**Supplementary Fig. S6j**) compared to those with hybrid immunity from two-dose vaccination followed by Omicron infection.

### Dissecting Tau phosphorylation pattern toward scalable blood-based AD assessment

It is well established that pTau 217 is an effective blood marker of AD, with several tests now FDA cleared for use in patients with cognitive impairment^(*28*)^. Considering the high prevalence and long progression of AD, decentralized and self-operated blood collection methods would prove highly valuable for drug development, early detection and disease management. Capillary microsampling devices, with their low-pain and low-stress experience and ability to meter the blood volume (which DBS lacks), have the potential to achieve both wide access and high sample quality, and have gained interest for AD screening in recent years^(*29*)^. However, it has been reported that capillary blood samples often show elevated readings of pTau 217 compared to matched venous samples^(*30,31*)^.

To confirm this finding, we initially developed a pTau 217 assay using an antibody binding the epitope containing phosphorylated T217 (conjugated to the P probe) and an antibody binding to the N-terminal region of Tau (around aa. 5–25, conjugated to the S probe), which we call the N-pTau 217 assay. Using a cohort of 17 subjects with matched venous and capillary blood samples (collected by the Tasso device), the SPEAR N-pTau 217 assay also showed substantially higher (geometric mean, or GM ratio of 8.3×) and discordant (ρ = 0.14) readings in capillary samples compared to the matched venous samples (**Fig. 3a**, left); so did the SPEAR BD-pTau 217 assay (GM ratio of 6.68×, ρ = 0.06, **Fig. 3a**, middle), even though targeting of the exon 4-5 junction was intended to exclude peripherally generated Tau proteins^(*32*)^. This mirrors what was reported for the NULISA BD-pTau 217 assay^(*30*)^.

**Figure 3.**
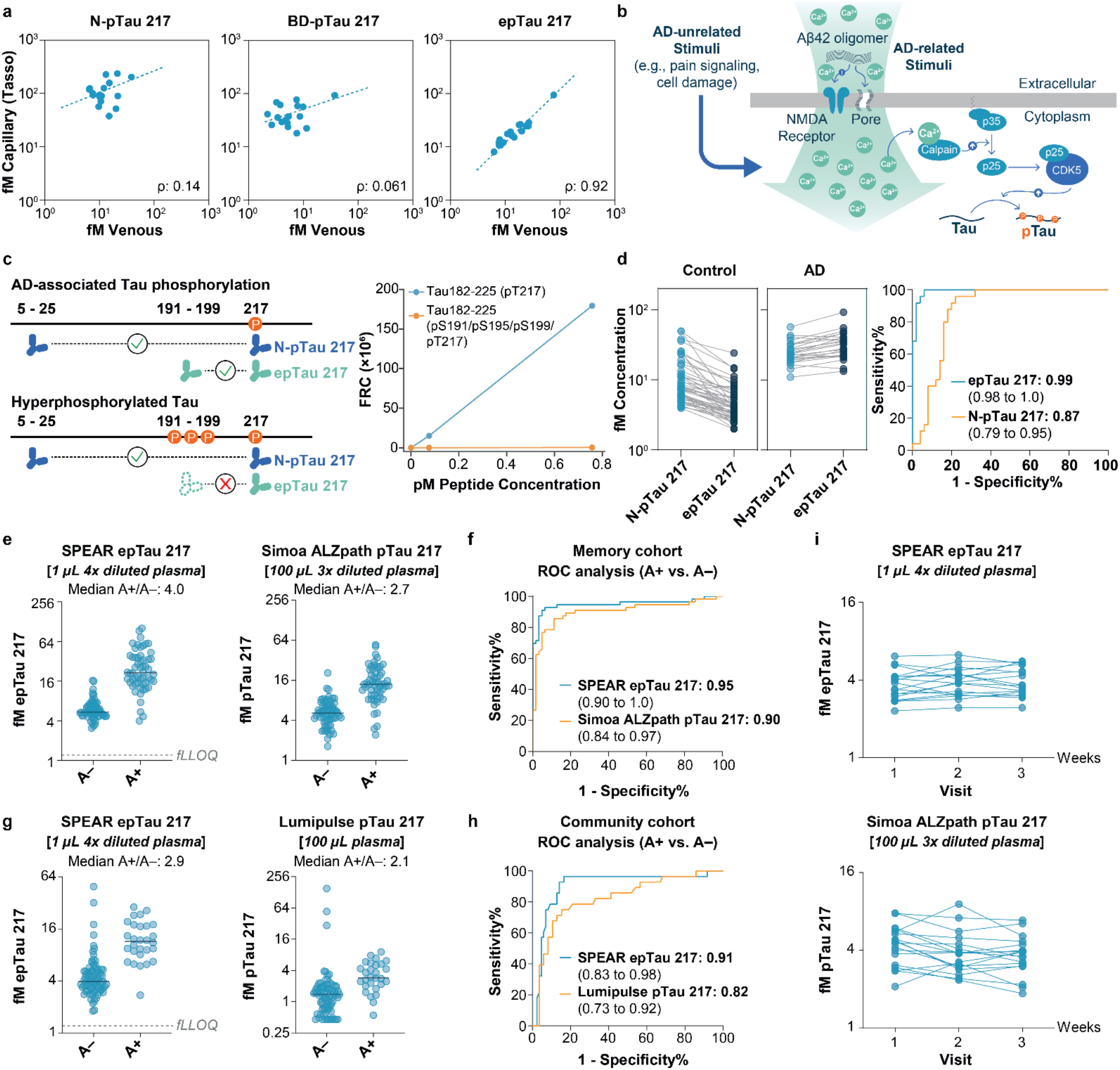
SPEAR epTau 217 assay resolves capillary-venous discordance and improves diagnostic power over pTau 217 assays. **a**, Concordance analysis of capillary and venous samples using N-pTau 217, BD-pTau 217 and epTau 217 assays. **b**, Putative signaling pathway leading to phosphorylation of Tau by CDK5. **c**, Left: Scheme of SPEAR N-pTau 217 and epTau 217 assays. Right: response of the SPEAR epTau 217 assay to Tau peptide 182–225 having T217 phosphorylation, with and without phosphorylation on S191, S195 and S199. **d**, Clinical performance of epTau 217 in a cohort containing 25 healthy controls, 25 ALS patients and 25 AD patients. **e**, Comparison of Simoa ALZpath and SPEAR epTau 217 assays in the UPMC ADRC cohort. **f**, ROC analysis of (**e**). **g**, Comparison of SPEAR epTau 217 and Lumipulse pTau 217 assays for amyloid PET-confirmed plasma (A−: *n* = 85; A+: *n* = 28) in a low-prevalence community cohort (Human Connectome Project). Fold change (A+/A−) indicated; bars show medians. **h**, ROC analysis for (**g**). **i**, Measurement variability across weekly plasma collections (*n* = 3 visits) from control donors without neurological conditions (*n* = 18) using Simoa ALZpath and SPEAR epTau 217 assays. Lines connect visits from the same donor.

We hypothesize that the abnormal increase in pTau 217 signal could be due to local phosphorylation and release of the Tau protein at the blood collection site. Since the dermis is densely innervated by nociceptive C-fibers and Aδ-fibers, which express Tau, its kinase CDK5, and the activator of CDK5, p35^(*33*)^, mechanical disruption and nociceptive signaling during capillary blood collection could trigger calcium influx in these sensory nerve endings, activating calpain, which cleaves p35 into p25 and thereby hyperactivates CDK5^(*33*)^ (**Fig. 3b**). To distinguish the brain-originated, AD-associated phosphorylated Tau from its dermis-originated counterpart, we investigated the potential utility of targeting phosphorylation patterns. It has been reported that pTau 217 and pTau 231 are among the earliest phosphorylation sites to become detectable in AD progression^(34)^. This observation is consistent with the *in vitro* sequence preference of CDK5^(*35*)^, which predicts T231 and T217 to be among the more favorable CDK5 substrate sites in Tau (**Supplementary Tables S4, S5**). In AD brains, such preference could contribute to a phosphorylation pattern in which preferred CDK5 sites (e.g., T217, T231) are phosphorylated and unpreferred CDK5 sites remain unphosphorylated. However, in the acute process of dermis damage, CDK5 activity may rise sharply and cause phosphorylation on both preferred and unpreferred sites. Therefore, a SPEAR assay that targets phosphorylated T217 but an unphosphorylated unpreferred CDK5 site may still detect brain-originated, AD-associated pTau 217, but reject the injury-induced, hyperphosphorylated Tau that also includes phosphorylated T217.

We specifically investigated S199 not only because of its much lower preference by CDK5^(*35*)^ (**Supplementary Table S5**), but also because of the presence of S195 and S191, two putative GSK3β substrates whose phosphorylation depends on S199 phosphorylation^(36)^. The proximity of these three sites offers an opportunity to use antibodies that are highly specific to the unphosphorylated state. We thus developed a SPEAR assay using an antibody targeting this region (around aa. 185–199) in its unphosphorylated state and paired it with the phospho-T217 antibody, which we call the epTau 217 assay for its intended use to detect Tau in its early phosphorylation state (**Fig. 3c**, left). Using peptides having residues 182 to 225 with phosphorylated T217 as model substrates, we confirmed that simultaneous phosphorylation of S191, S195 and S199 abolished the detectability of the peptide by this SPEAR assay (**Fig. 3c**, right). Since Tau protein in this phosphorylation state is a subset of pTau 217, its detection requires higher sensitivity than what is needed to detect pTau 217, which is already a low-abundance biomarker. Nevertheless, this SPEAR assay (LOD = 0.03 fM, LLOQ = 0.30 fM, **Supplementary Fig. S3i**) successfully detected and quantified epTau 217 in all capillary and venous samples, and more importantly, yielded highly consistent readings between these two sample types (GM ratio of 1.28×, ρ of 0.92, **Fig. 3a**, right).

Because calcium influx and the subsequent CDK5 hyperactivation may occur in many biological processes aside from AD pathology (including diabetes^(*37–40*)^ and other forms of brain pathology), we asked whether the epTau 217 assay may improve the specificity of AD diagnosis compared to the N-pTau 217 assay in venous blood. We used plasma samples for the studies below because of the availability of well-characterized sample cohorts. In a cohort of 75 individuals including 50 non-AD controls (25 healthy controls and 25 ALS) and 25 clinically confirmed AD patients, the epTau 217 assay (AUC = 0.99) substantially outperformed the N-pTau 217 assay (AUC = 0.87) in distinguishing AD patients from non-AD controls (**Fig. 3d**, right). This improvement stems from lower readings in the control group (**Fig. 3d**, left), possibly due to rejection of hyperphosphorylated Tau originating in non-AD conditions, and higher readings in the AD group (**Fig. 3d**, middle), possibly due to detection of degraded Tau protein that lost the N-terminal epitope.

We next carried out a head-to-head comparison with the widely adopted Simoa ALZpath pTau 217 assay (a dELISA assay) in a memory clinic cohort (the UPMC ADRC cohort) with subjects whose amyloid pathology was confirmed by amyloid PET scan and labeled as A+ and A− (*n* = 56 and *n* = 63, respectively). The SPEAR epTau 217 assay showed a substantially higher fold change of the median reading between the A+ and A− groups (“median A+/A− fold change”, 4.0-fold) than the dELISA assay (2.7-fold, **Fig. 3e**), as well as higher AUC (0.95 vs. 0.90, **Fig. 3f**). The epTau 217 level also correlated with PET-based amyloid PiB standardized uptake value ratio (SUVR) (Spearman ρ = 0.80, **Supplementary Fig. S7a**). In another memory clinic cohort where amyloid status was assigned by CSF-based testing evaluating Aβ42/40 ratio, pTau 181 level and total Tau level (the MGH CSF cohort, *n* = 88; A+, *n* = 40; A−, *n* = 48), the A+ group also showed substantially higher (6.3-fold) median epTau 217 readings than the A− group (AUC = 0.99, **Supplementary Fig. S7b**).

These data prompted us to explore whether the epTau 217 assay could contribute to broad screening of AD. The patient cohorts in the studies above were from well-controlled clinical studies operated in memory clinics, while broad screening will face more variability such as in participant ethnicity and lower positivity rate. We therefore tested the SPEAR epTau 217 assay in a community cohort (the Human Connectome Project cohort) and compared its performance to Lumipulse pTau 217, a well-established assay based on chemiluminescent enzyme immunoassay (CLEIA)^(*41*)^. Both median A+/A− ratio and AUC of the SPEAR epTau217 assay exceeded those of the Lumipulse pTau 217 assay (2.9-vs 2.1-fold and 0.91 vs 0.82, respectively, **Fig. 3g, h**).

Finally, the ability to reject Tau phosphorylated at both T217 and S191/195/199, which may arise from non-AD-associated hyperphosphorylation, may also prove helpful in reliable tracking of the change of AD-specific T217 phosphorylation during disease progression or drug response. To this end, we tested samples collected during three consecutive weekly visits by non-AD subjects (the ‘MGH longitudinal’ cohort) using both dELISA (Simoa ALZpath pTau 217) and the SPEAR epTau 217 assay. During this span the AD-related pTau 217 level was not expected to change. As shown in **Fig. 3i**, while the dELISA results displayed considerable visit-to-visit variability for some subjects, the SPEAR epTau 217 results remained consistent, suggesting that it could reduce variability in longitudinal monitoring.

## DISCUSSION

SPEAR achieves near single-molecule detection sensitivity, which we define as an LOD below 100 molecules, a level where Poisson sampling error alone limits precision to ~10% CV. The key to the improved sensitivity of SPEAR lies in its ability to compound discrimination between analyte-dependent and background reactions through engineered kinetic proofreading. The high sensitivity of SPEAR allowed us to carry out challenging assays using remotely collected samples, such as immune surveillance of infectious diseases and resolving subspecies of the Tau protein with different posttranslational modification patterns. In preliminary studies with relatively small cohorts, the epTau 217 assay showed improved clinical performance over conventional pTau 217 assays. As capillary microsampling devices continue to gain popularity, evaluating the utility of the epTau 217 assay in AD diagnostics using remotely collected samples in large-scale studies would be of particular interest. Looking forward, SPEAR is in principle compatible with multiplexed PCR and with sequencing-based readouts for protein profiling, which would further facilitate discovery of novel low-abundance biomarkers. The small sample volume capability also means a single archived aliquot of clinical sample can potentially support hundreds of measurements rather than a handful, allowing samples from a well-characterized cohort to be repeatedly utilized for biomarker discovery and assay development. Finally, the separation-free nature and qPCR readout of SPEAR also enable its operational simplicity and accessibility, and make low-abundance biomarkers previously only detectable with specialized instruments now detectable on common lab instruments, and with better performance in our comparisons.

