## Supplementary Information for "Achieving near single-molecule sensitivity in separation-free protein detection using engineered kinetic proofreading"

*2.1.3 LOD for allostery-based assays* ... 17
*2.1.4 LOD for proximity-based signal generation* ... 19
*2.1.5 Enhancement of rate differentiation by two-step kinetic proofreading* ... 23
*2.1.6 LOD for SPEAR* ... 25
*2.1.7 LODs under realistic parameters* ... 27
3 Supplementary Figures ... 30
Fig. S1. Simulation of signal loss due to spontaneous dissociation between analyte and antibody in wash-based immunoassays and in separation-free SPEAR assay. ... 30
Fig. S2. SPEAR proof-of-concept validation using streptavidin assay (SA). ... 32
Fig. S3. Calibration curves and analytical sensitivity of SPEAR immunoassays. ... 33
Fig. S4. Calibration curves for LOD estimation for SPEAR, dELISA (Simoa) and conventional ELISA for assays using identical antibodies. ... 34
Fig. S5. Theoretically achievable LODs under realistic parameters. ... 35
Fig. S6. Performance of SPEAR SARS-CoV-2 Neutralization Assay. ... 36
Fig. S7. Clinical performance of SPEAR epTau 217 Assay for amyloid PET confirmation ... 38
Fig. S8. General concepts and schemes. ... 39
Fig. S9. Theoretically achievable LOD in competition-based protein-detection assays as a function of the concentrations of the signal-generating molecule (CE) and binder (CB). ... 40
Fig. S10. LOD of allostery-based binders under realistic parameters. ... 41
Fig. S11. Abstract models for proximity-based immunoassays. ... 42
Fig. S12. Kinetics of full record formation in SPEAR assays to detect IFN-$\gamma$ (left) and streptavidin (right). ... 43
4 Supplementary Tables ... 44
Table S6. Participant demographics for capillary and Tasso concordance study from Memory/Cognitive Disorders clinic at Banner Sun Health Research Institute ... 44
Table S7. Participant demographics for the Bio-Hermes-002 cohort. ... 45
Table S8. Participant demographics for the Pitt-ADRC cohort. ... 46
Table S9. Participant demographics of the CSF AD biomarker confirmed cohort from MGH. ... 47
Table S10. Participant demographics for the Human Connectome Project (HCP) cohort. ... 48

#### 1 Materials and Methods

##### *1.1 SPEAR Oligonucleotides Design and Probe Preparation*

Oligonucleotides were obtained from IDT in lab-ready format (1× TE, pH 7.5), except those intended for protein conjugation, which were reconstituted in nuclease-free water. Oligonucleotides at concentrations below 1 µM were diluted in 1× TE (pH 7.5) supplemented with 20 µg/mL tRNA. Probe S was prepared by incubating Strand A and Strand B in 1× Isothermal Amplification Buffer containing 8 mM $MgSO_4$. Annealing was performed using the following thermal program: 85 °C to 65 °C with 10 min/°C decrements, then 65 °C to 55 °C with 5 min/°C decrements, followed by a final hold at 4 °C.

##### *1.2 Antibody Conjugation and Purification*

SPEAR probes were conjugated to antibodies using lysine chemistry adapted from Agasti et al.[1]. The Primer P strand and a Linker strand, which hybridizes to the duplex Probe S, were thiol-modified at the 5′ and 3′ ends, respectively, and reduced by incubation with 100 mM DTT in 1× PBS (pH 7.9) containing 1 mM EDTA at 4 °C for 2 hours. The mixture was purified using a NAP™-5 column (Cytiva), and fractions with maximal absorbance at 260 nm were identified using a NanoDrop spectrophotometer and pooled. Antibodies were concentrated with a 50 kDa Amicon Ultra-0.5 centrifugal filter unit and reacted with PEGylated SMCC crosslinker (Thermo Fisher Scientific™) at 4 °C for 2.5 hours. Excess crosslinkers were removed using a Zeba™ Spin Desalting Column. SMCC-modified antibodies were then incubated with thiol-reduced DNA at 4 °C overnight.

The conjugate mixture was subsequently exchanged into 1× Tris buffer (pH 7.0) using a 50 kDa Amicon Ultra-0.5 centrifugal filter unit and further purified through a Pierce™ Strong Anion Exchange Spin Column by centrifugation at 2000 × g for 5 minutes at 4 °C with a gradient of NaCl concentrations. Elution fractions were selected based on band intensity and purity confirmed by SDS denaturing PAGE using Krypton (protein) and Sybr Gold (DNA) stains.

##### *1.3 Streptavidin Gel Electrophoresis Test for Validating SPEAR Assay*

Biotinylated Probe S (50 nM) and Cy5-labeled Primer P (50 nM) were incubated with or without streptavidin (25 nM) in 1× Isothermal Amplification Buffer at room temperature for 20 minutes. The reaction mixture was then supplemented with 125 µM each of dATP, dTTP, and dCTP, along with 10 U/µL Bst 2.0 polymerase in 1× Isothermal Amplification Buffer and incubated at 37 °C. Aliquots of 10 µL were collected at defined timepoints (1, 5, 15, 30, and 60 minutes). Collected samples were immediately quenched with EDTA and formamide, followed by heat inactivation at 90 °C for 5 minutes and rapid chilling on a cooling block. Denatured samples were run on 15% TBE-Urea PAGE (Novex®) gels at 180 V for 50 minutes at 65 °C. Gels were stained with Sybr Gold for 15 minutes and scanned using a Typhoon™ gel scanner with Cy5 (Ex 647/Em 670) and Sybr Gold (Ex 495/Em 537) fluorescence channels. Gel band intensities were quantified using ImageJ software[2].

##### *1.4 SPEAR Streptavidin Assays*

Biotinylated SPEAR probes (50 pM, 1 µL) were incubated with streptavidin (1 µL) in 0.1% Bovine Serum Albumin and 1× Isothermal Amplification Buffer at 4 °C overnight. Samples were equilibrated at 37 °C for 20 minutes and reacted with Bst 2.0 polymerase (8 U/10 µL) and 100 µM dATP, dTTP, and dCTP in 1× Isothermal Amplification Buffer at 37 °C for 5 minutes, followed by inactivation at 85 °C for 30 minutes. Reaction products were then mixed with qPCR reagents consisting of 250 nM PCR probe, 500 nM forward and reverse primers, and 1× PrimeTime® Gene Expression Master Mix. qPCR was performed on a Bio-Rad instrument with the following thermal program: 95 °C for 3 minutes, 40 cycles of 95 °C for 15 seconds and 55.9 °C for 1 minute, and a final hold at 10 °C. Fluorescence was measured in the FAM channel using CFX Maestro™ Analysis Software.

For comparative studies of single and double proximity extensions, Probe S constructs were designed with either single or double toeholds (see **Fig. S2b** for schematics).

The Full Record Count (FRC) represents the absolute copy number of full-length SPEAR amplicon molecules ("full records") and serves as the primary readout for all SPEAR assays. After the two-step extension reaction is complete and the polymerase is heat-inactivated, the reaction products are quantified by quantitative PCR (qPCR) using primers and a fluorogenic probe targeting the full record sequence. Ct values are converted to copy numbers (FRC) by normalization with a synthetic reference DNA preparation whose absolute concentration is determined using limiting dilution PCR. Next, analyte concentrations are interpolated from the standard curve (FRC versus calibrator concentration) fitted by 4PL logistic fitting.

##### *1.5 Cytokine Sensitivity Test by SPEAR*

Calibration curves were established by serial dilution of antigens in SPEAR Assay Diluent (1% BSA, 1× PBS, pH 7.9, and 0.1% Tween-20, heterophilic blocking reagents) using 10-fold dilution series. SPEAR probes conjugated to the corresponding affinity reagents, or Probe S and Primer P sequences embedded within aptamers for PDGF-BB detection, were diluted in Probe Diluent (1% BSA, 1× Isothermal Amplification Buffer, 0.1% Tween-20, 60 µg/mL sperm DNA, and 2 mM $MgSO_4$) to final concentrations ranging from 50 pM to 500 pM. One microliter of probe solution was incubated with 1 µL of diluted calibrators or biological samples at the minimal required dilutions (4-fold for plasma or serum) at 37 °C for 1 hour. Antibodies for cytokine assays were purchased from BD Biosciences, and calibrators were obtained from R&D Systems.

Following incubation, samples were reacted with 6 µL of reaction mixture containing 8 U/10 µL Bst2.0 DNA polymerase and 100 µM each of dATP, dTTP, and dCTP in 1× Isothermal Amplification Buffer. Extension reactions were performed at 37 °C for 8 minutes, followed by enzyme inactivation at 85 °C for 30 minutes and cooling to 4 °C. Quantitative PCR was subsequently conducted on a Bio-Rad instrument using the qPCR mixture and thermal cycling program described in **Section 1.4**. Data were fitted using four-parameter logistic regression, and limits of detection (LOD) were defined as the blank signal ± 2.5 standard deviations.

##### *1.6 Cytokine sensitivity test by Simoa*

Simoa assays were performed on an HD-X Analyzer (Quanterix). For each assay, the antibody pairs and calibrators were identical to those used in the corresponding SPEAR assays (IL-2:

5344.111 and B33-2; IL-4: 8D4-8 and MP4-25D2; IFN-γ: NIB42 and 4S.B3; all from BD Biosciences). Preparation of capture antibodies and experimental procedures followed the protocol described by Rivnak et al.[3], while detection antibodies were supplied pre-conjugated to biotin. All assays were conducted using a two-step immunoassay format. Calibration curves were generated by serial dilution of recombinant proteins in Homebrew Detector/Sample Diluent (Quanterix). Signal detection and data analysis were performed using the manufacturer's software, with sensitivity and dynamic range determined from the calibration curves. Data were fitted using four-parameter logistic regression, and LOD was defined as the blank signal ± 2.5 standard deviations.

##### *1.7 Cytokine Sensitivity Test by ELISA*

Cytokine sensitivity was assessed using BD OptEIA™ ELISA kits. Antibody pairs and calibrators were identical to those employed in the corresponding SPEAR assays described in **Section 1.6**. Capture antibodies were coated onto MaxiSorp™ 96-well ELISA plates at concentrations of 4 µg/mL for IL-2 and 8 µg/mL for the remaining cytokines, followed by incubation at 4 °C overnight. Plates were washed six times with washing buffer after antibody coating and subsequently incubated with human recombinant cytokine proteins at concentrations ranging from 0.1 pg/mL to 1640 pg/mL at room temperature for 2 hours. Plates were then washed six times with washing buffer.

Biotin-labeled detection antibodies (1 µg/mL) for IL-2, IL-4, and IFN-γ were combined with streptavidin–HRP (1:1000 dilution) and incubated with the plates at room temperature for 1 hour, followed by six washes. Substrate reactions were performed in the dark for 30 minutes, and absorbance was measured at 450 nm using a microplate reader. Data were analyzed by fitting four-parameter logistic regression models, and LOD was defined as the blank signal ± 2.5 standard deviations.

##### *1.8 Dilution Linearity of SPEAR and Simoa Cytokine Assays in Human Serum*

Two human serum samples were serially diluted from 4-fold to 128-fold in either SPEAR Assay Diluent or Simoa Homebrew Detector/Sample Diluent, corresponding to the SPEAR and Simoa assays, respectively. Intrinsic concentrations of human IL-2, IL-4, and IFN-γ were quantified using both platforms, employing identical antibody clone pairs as described in **Section 1.6**. In the SPEAR assay, each dilution was tested with two independent biological replicates, and each replicate was analyzed with two qPCR technical repeats to ensure reproducibility.

##### *1.9 SPEAR SARS-CoV-2 Immunoassays*

###### *1.9.1 DBS specimen collection for longitudinal studies*

Healthy human subjects aged 18 to 65 years old without a history of immunological diseases or treatment with immune-modulating agents were recruited at the Wyss Institute of Harvard University, and informed consent was obtained. The study was approved by the Institutional Review Board of Harvard University and performed in accordance with relevant guidelines and regulations for the protection of human subjects.

Eligible participants completed a survey regarding prior COVID-19 exposure and vaccination status. Prior to blood collection, participants were instructed to drink adequate water and wash their hands with warm water to enhance local blood flow. Capillary blood was collected via sterilized finger prick from the tip of the middle or ring finger and dropped onto Whatman® 903 protein saver cards, ensuring coverage sufficient for at least one 6 mm punch. Following collection, DBS samples were air-dried for at least four hours at ambient temperature, and were placed into a foil ziplock biohazard with desiccants and stored at -20 ˚C.

##### *1.9.2 DBS, plasma and serum specimen for comparative studies of SPEAR vs CPass vs PsVNA*

Negative DBS samples from individuals with no self-reported history of COVID-19, COVID-19 convalescent patients, and SARS-CoV-2 vaccine recipients (Pfizer-BioNTech or Moderna) were obtained from BioIVT, Inc. (Westbury, NY, USA). Paired venous serum/plasma and DBS samples from SARS-CoV-2 vaccine recipients for platform concordance studies were sourced from BioIVT, Inc. (Westbury, NY, USA) and Precision for Medicine, Inc. (Bethesda, MD, USA).

##### *1.9.3 DBS punching and elution*

6 mm DBS punches were obtained using either a manual or semi-automated DBS puncher and placed into 1.5 mL low protein binding Eppendorf tubes. To prevent cross-contamination, the puncher was cleaned by punching blank cards five times between samples. Each punch was eluted in 500 µL DBS elution buffer (1% BSA, 0.2% NP40, 0.2 × Protease inhibitor in 1 × PBS) for 1 hour at 37 °C with shaking at 500 rpm. Following elution, samples were centrifuged at 10,000 × g for 10 minutes, and 400 µL of supernatant was collected per sample. Eluates were stored at 4 °C for up to 3 days or snap-frozen in liquid nitrogen and stored at -80 °C for long-term preservation.

##### *1.9.4 SPEAR SARS-CoV-2 Spike and Pan-Ig measurements*

SPEAR measurements of SARS-CoV-2 Spike and Pan-Ig were conducted using 1 µL of neat DBS eluate and 1 µL of DBS eluate diluted 50-fold in elution buffer. Respective probes comprising anti-SARS-CoV-2 RBD antibodies (GenScript) and SARS-CoV-2 S1 proteins (BioLegend) were incubated at 37 °C for 1 hour, followed by the SPEAR enzymatic reaction and quantitative PCR (qPCR).

##### *1.9.5 SARS-CoV-2 neutralizing antibody detection assay*

SPEAR SARS-CoV-2 neutralization assays were performed using a competitive binding format with two sequential incubation steps. Briefly, 1 µL of sample was incubated with 1 nM SPEAR Probe S conjugated S1 protein (Fc tagged S1, BioLegend) at 25 °C for 2 hours, followed by incubation with 5 pM Primer P constructed with angiotensin-converting enzyme 2 (Fc-tagged ACE2, BioLegend) at 25 °C for 30 minutes. The SPEAR reaction was then carried out using 3.75 U Bst 2.0 polymerase and 100 µM dATP, dTTP, and dCTP at 25 °C, followed by qPCR. Neutralizing antibodies reduced ACE2 binding to S1 proteins, resulting in lower SPEAR signals. Inhibition was calculated relative to the negative control using the following equation:

$$\text{Inhibition} = \left(1 - \frac{1}{2^{\Delta \mathrm{Ct}}}\right) \times 100\% \qquad \mathbf{(1.1)}$$

For quantitative neutralization titer measurement, DBS samples were serially diluted 1:2 from 2- to 256-fold, and plasma/serum samples were diluted 1:4 from 10- to 163,840-fold. Inhibition curves were generated using four-parameter logistic (4PL) sigmoidal fitting, and the dilution fold corresponding to 50% inhibition (NT50) was interpolated using Prism analysis software. The final NT50 in whole blood was corrected by a dilution factor of 50 for DBS, based on the estimation that a 6 mm DBS punch contained approximate 10 µL of blood eluted into 500 µL of DBS elution buffer.

##### *1.9.6 CPass™ SARS-CoV-2 neutralization antibody detection assay*

Neutralizing antibodies were measured using the CPass™ SARS-CoV-2 Neutralization Antibody Detection Kit (GenScript) following the manufacturer's protocol. ACE2-coated plates were blocked with assay diluent for 1 hour at room temperature. Samples and controls were diluted with sample dilution buffer at 1:9, incubated with HRP-conjugated RBD at 37 °C for 30 minutes, then transferred to ACE2-coated plates and incubated at 37 °C for 15 minutes. After washing, TMB substrate was added for 15 minutes, and absorbance was measured at 450 nm. Neutralizing activity was expressed as the percentage inhibition:

$$\text{Inhibition} = \left(1 - \frac{\text{OD value of sample}}{\text{OD value of negative control sample}}\right) \times 100\% \qquad \textbf{(1.2)}$$

##### *1.9.7 Plaque reduction neutralization test (PRNT)*

The PRNT assay was conducted by Corgenix, Inc. (Broomfield, CO, USA). Vero E6 cells were seeded in six-well plates, and SARS-CoV-2 viruses (WA01/2020 isolate) were diluted to approximately 2000 pfu/mL. Following 1:1 mixing with serum/plasma samples, the final virus concentration was ~1000 pfu/mL. Serum/plasma samples from vaccine recipients were serially diluted two-fold in BA-1 medium from 1:5 to 1:10,240 (equivalent to 1:10 to 1:20,480 after mixing with virus). Diluted samples were incubated with virus at 37 °C for 1 hour, then added to Vero E6 cells and incubated at 37 °C for 45 minutes with rocking every 15 min. Cell monolayers were overlaid with 1% agarose in culture medium and incubated at 37 °C for 24 hours. A second overlay containing 1% agarose and 0.005% neutral red was applied, followed by incubation at 37 °C for another 24 hours. Plaques were counted in each well. Each assay included three negative controls and one positive control. Inhibition values for each dilution were calculated as:

$$\text{Inhibition} = \left(1 - \frac{\text{Plaque number of sample}}{\text{Plaque number of negative control sample}}\right) \times 100\% \qquad \textbf{(1.3)}$$

NT50 was determined by fitting inhibition versus dilution curves using 4PL sigmoidal regression.

##### *1.9.8 Pseudovirus-based neutralization assay (PsVNA)*

SARS-CoV-2 spike–pseudotyped HIV virions were generated in-house, and neutralization assays were performed as described by Si et al.[4]. Serum/plasma samples were serially diluted two-fold in 1% BSA and 1× PBS (pH 7.4) from 1:2 to 1:256, corresponding to final dilutions of 1:20 to 1:2560 after mixing with virus and cells. For each reaction, 20 µL of diluted sample was combined with 80 µL pseudovirus (Multiplicities of infection, MOI = 0.1) and incubated at 37 °C for 30 minutes. The mixture was then added to 100 µL medium containing 20,000 ACE2-293T cells and incubated at 37 °C for 72 hours. Luciferase activity, reflecting pseudovirus entry into host cells, was quantified using the Bright-Glo™ Luciferase Assay System (Promega).

##### *1.9.9 Binding affinity evaluation for anti-SARS-CoV-2 antibodies using ELISA*

Monoclonal antibodies targeting the SARS-CoV-2 RBD (BioLegend) were coated onto Max-iSorp™ 96-well ELISA plates at 1 µg/mL and incubated overnight at 4 °C. Plates were washed and blocked with assay diluent for 1 hour at room temperature. RBD-targeting monoclonal antibodies (GenScript) were serially diluted 1:4 in SARS-CoV-2 antibody–negative DBS eluates, starting at 8 µg/mL, and incubated with RBD-coated plates for 1 hour at room temperature. After washing, HRP-conjugated secondary antibodies (anti-human 1:10,000; anti-mouse 1:25,000; anti-rabbit 1:10,000) were added for 1 hour at room temperature. Substrate reaction was performed with TMB for 15 min in the dark, and absorbance was measured at 450 nm using a microplate reader.

#### *1.10 Neuroassays Measurements*

##### *1.10.1 SPEAR NFL and pTau 231 measurement*

NFL and pTau 231 concentrations were measured using the SPEAR UltraDetect™ NFL Assay Kit (Catalog#: SPR90004) and SPEAR UltraDetect™ pTau 231 Assay Kit (Catalog#: SPR90009), respectively, in combination with the SPEAR UltraDetect™ Reaction Kit (Catalog#: SPR90024). SPEAR probes and calibrators were reconstituted with the Kit Probe Diluent and Assay Diluent, respectively. For each reaction well, 1 µL of calibrator or diluted sample was used. Serum and plasma samples were diluted 4-fold, whereas CSF samples were diluted 100-fold. Samples were processed using a semi-automated FAST workflow according to the assay manuals, and qPCR readouts were acquired using QuantStudio instruments. Matrix comparability for NFL was assessed using commercially obtained healthy control samples from Precision for Medicine, including matched K2EDTA plasma ($n$ = 10) and serum ($n$ = 10), as well as unmatched lithium-heparin plasma ($n$ = 10) and CSF ($n$ = 12).

Cross-platform comparison of pTau 231 measurements was performed using K2EDTA plasma samples from healthy controls ($n$ = 25) and individuals with clinically diagnosed AD ($n$ = 15) obtained from Precision for Medicine and BioIVT. SPEAR pTau 231 measurements were performed using 1 µL of 4-fold diluted plasma per reaction well. Comparative Simoa measurements were performed at the Clinical and Translational Research Unit (CTRU) Biomarker Core at Massachusetts General Hospital (MGH) using 100 µL of 2-fold diluted plasma with the Simoa® pTau231 Advantage PLUS Kit on the HD-X platform (Quanterix, Catalog#: 104512).

Concordance of NFL concentration measurements between SPEAR and Simoa was evaluated using plasma samples from healthy controls ($n$ = 8) and individuals with clinically confirmed Alzheimer's disease (AD; n = 6), amyotrophic lateral sclerosis (ALS; $n$ = 1), progressive supranuclear palsy (PSP; $n$ = 5), and frontotemporal dementia (FTD; $n$ = 6), commercially obtained from Precision for Medicine and BioIVT. Simoa NFL measurements were performed using the Simoa® NF-Light Advantage PLUS Kit on the HD-X platform (Quanterix; Catalog No. 104364) according to the manufacturer's instructions. These samples were used to assess assay quantifiability, precision and cross-platform concordance.

##### *1.10.2 SPEAR and Nulisa BD-pTau 217 comparison study*

K2EDTA plasma samples from healthy controls ($n$ = 44) and individuals with a clinical diagnosis of Alzheimer's disease (AD, $n$ = 30) were purchased commercially from Precision for

Medicine. SPEAR BD-pTau 217 was measured at Spear Bio using the SPEAR BD-pTau 217 Development Assay Kit in combination with the SPEAR UltraDetect™ Reaction Kit (Catalog#: SPR90024). SPEAR probes and calibrators were reconstituted with the kit Probe Diluent and Assay Diluent, respectively. For each reaction, 1 µL of calibrator or four-fold diluted plasma sample was analyzed using a semi-automated FAST workflow. qPCR readouts were acquired using QuantStudio instruments.

NULISA BD-pTau217 was measured at the CTRU Biomarker Core at MGH according to the manufacturer's instructions using the NULISAqpcr™ BD-pTau217 Assay on the ARGO™ HT System (Catalog#: 800470), with 25 µL of undiluted plasma per sample.

##### 1.10.3 *SPEAR eMTBR-Tau peptide specificity and Bio-Hermes-002 clinical measurement*

To evaluate the C-terminal specificity of the SPEAR eMTBR-Tau Assay for the V256 endpoint, the SPEAR eMTBR-Tau Development Assay Kit was used in combination with the SPEAR UltraDetect™ Reaction Kit (Catalog#: SPR90024). Synthetic tau peptides (LifeProtein) were tested at 10 pM, including peptides spanning T220–N255, T220–V256, and T220–K257. An additional peptide panel containing deamidated N255 (dN255) included T220–N255, T220–V256, T220–K257, T220–I260, and T220–Q269 peptides. Each peptide was tested using 1 µL per reaction well.

Clinical sensitivity was evaluated using K2EDTA plasma samples from participants enrolled in the Bio-Hermes-002 study (NCT06584357), an observational biomarker study conducted by the Global Alzheimer's Platform Foundation and comprising participants across the Alzheimer's disease clinical continuum. Participants underwent clinical and cognitive assessments, blood collection, brain MRI, amyloid PET, and tau PET imaging. Tau PET imaging in the analyzed cohort was performed using either [^18F]MK-6240 or [^18F]flortaucipir. Detailed demographics of the cohort are provided in **Table S7**. Plasma samples were categorized according to amyloid and tau PET status and clinical diagnosis into three groups: A−T−, comprising cognitively normal participants negative for both amyloid and tau PET ($n$ = 33); A+T−, comprising amyloid PET-positive and tau PET-negative participants with cognitively normal $n$ = 11), mild cognitive impairment (MCI, $n$ = 16), or Alzheimer's disease (AD, $n$ =10) diagnoses; and A+T+, comprising participants positive for both amyloid and tau PET with MCI ($n$ = 25) or AD ($n$ = 32) diagnoses. Plasma eMTBR-Tau concentrations were measured at Spear Bio with 1 µL of 4-fold diluted plasma per reaction well, and associations with imaging-defined amyloid and tau pathology were evaluated.

##### 1.10.4 *Cross-site precision studies of NFL measurements across qPCR machines*

Cross-site precision studies were conducted to evaluate reproducibility of NFL measurements across multiple qPCR instruments. The SPEAR UltraDetect™ NFL Assay Kit (Catalog#: SPR90004) was employed to quantify NFL concentrations from a predefined set of plasma and serum samples, using 1 µL of diluted material. Assays were paired with the SPEAR UltraDetect™ Reaction Kit (Catalog#: SPR90024) and performed according to manufacturer protocols in both 96- and 384-well formats. Final readouts were acquired on multiple qPCR platforms, including BioRad, QuantStudio 5, QuantStudio 7 Pro, QuantStudio 12K, and Roche instruments.

##### 1.10.5 *SPEAR epTau 217, BD-pTau 217 and N-pTau 217 capillary (Tasso) and Venous plasma concordance comparison*

###### 1.10.5.1 Consent Statement

A total of 17 participants were recruited from longitudinal cohort studies of aging and neurodegenerative disorders and the Memory/Cognitive Disorders clinic at Banner Sun Health Research Institute (Sun City, Arizona).

All participants' cognitive, functional and clinical status was characterized through integrated assessment of validated instruments and cognitive testing. Participants underwent informed consent process and assessment of decisional capacity and they, or their legally authorized representatives, as appropriate, provided signed consent. The study was approved by the WCG Western Institutional Review Board.

1.10.5.2 Sample collection

Each participant's capillary blood was collected using the Tasso+ and venous blood was collected by standard venipuncture. Both sample types were collected into ethylenediaminetetraacetic acid (EDTA) anticoagulant tubes. For Tasso+ collection, the upper arm was warmed with a heat pack then cleansed with 70% isopropyl alcohol. The device was applied to the upper arm and activated according to the manufacturer's instructions by pressing the release button to deploy the lancet. Blood was allowed to collect for up to 10 minutes or until the collection tube was filled. Following collection, the tube was capped and gently inverted 10–12 times to ensure adequate mixing with the anticoagulant. A phlebotomist collected venous blood into 10 mL EDTA tubes, which were similarly inverted 10–12 times immediately after collection. All samples were transferred directly to laboratory personnel for processing immediately after acquisition.

1.10.5.3 Sample Processing

Samples were immediately placed on wet ice. Venipuncture tubes were centrifuged at 3500×g for 10 minutes at 4ºC. Venous plasma was then aliquoted into 1.5 mL protein LoBind tubes (Eppendorf). Tasso+ EDTA tubes were also centrifuged at 3500×g for 10 minutes at 4ºC. Plasma was transferred to a 1.5 mL protein LoBind tube and re-spun with the same settings. Tasso+ plasma was then aliquoted into a final 1.5 mL protein LoBind tube. Samples were stored at -80ºC until analysis. Detailed demographics of the cohort are provided in **Table S6**.

1.10.5.4 SPEAR epTau 217, BD-pTau 217, and N-pTau 217 Measurements

After thawing, samples were diluted 4-fold and measured at Spear Bio using the SPEAR epTau 217 Development Assay Kit, SPEAR BD-pTau 217 Development Assay Kit, and SPEAR N-terminal pTau 217 Development Assay Kit. The SPEAR N-terminal pTau 217 assay was constructed using the same anti-phospho-threonine 217 antibody as the SPEAR epTau 217 Development Assay Kit, paired with Tau13 (BioLegend) and Probe S. qPCR readouts were acquired using QuantStudio instruments.

##### *1.10.6 SPEAR epTau 217 and N-Tau 217 clinical comparison*

Non-AD control samples (*n* = 50; 25 healthy controls and 25 patients with ALS) and samples from individuals with clinically confirmed Alzheimer's disease (AD, *n* = 25) were commercially obtained from Precision for Medicine and evaluated at Spear Bio. epTau 217 and N-Tau 217 were measured using the SPEAR epTau 217 Development Assay Kit and SPEAR N-terminal pTau 217 assay as described above. Both assays were performed using the same calibrator, Probe Diluent, and Assay Diluent supplied with the SPEAR epTau 217 Development Assay Kit. Samples were diluted 4-fold and tested using 1 µL per reaction well. qPCR readouts were acquired using QuantStudio instruments.

##### *1.10.7 Specimen collection and epTau 217 measurement for ADRC-Pitt Cohort*

This study involved blinded testing of EDTA plasma samples from a subgroup of participants ($n = 119$) recruited through the University of Pittsburgh Alzheimer's Disease Research Center (ADRC), Pittsburgh, Pennsylvania, USA. All participants had amyloid pathology confirmed by positron emission tomography (PET) using Pittsburgh compound B (PiB) tracer. The study protocol was approved by the University of Pittsburgh Institutional Review Board (MOD19110245-023). Participants underwent annual clinical evaluations to monitor brain health and assess cognitive decline. Written informed consent was obtained from all participants in accordance with the ethical standards of the University's Human Research Protection Office.

Blood samples were collected via venipuncture by trained nursing staff between 9:00 am and 2:00 pm, with documentation of the time of the last meal. Each sample was drawn into a 10 mL lavender-top EDTA plasma tube (BD Biosciences, Catalog#: 366643), inverted 8–10 times, centrifuged at 2000×g for 10 minutes at 4 °C, aliquoted into cryovials, and stored at −80 °C until analysis. Amyloid (A) status was determined from global [11C] PiB standardized uptake value ratios (SUVR), calculated by volume-weighted averaging across nine composite regions (anterior cingulate, posterior cingulate, insula, superior frontal cortex, orbitofrontal cortex, lateral temporal cortex, parietal cortex, precuneus, and ventral striatum). Participants were classified as amyloid-positive (A+) if SUVR > 1.346. Details of the cohort demographics are provided in **Table S8**.

epTau 217 or pTau 217 concentrations were measured using both the SPEAR and Simoa ALZpath pTau-217 assays. SPEAR measurements were conducted using 1 µL of 4-fold diluted clarified plasma, prepared by centrifugation of thawed EDTA plasma at 10,000×g for 5 minutes. Assays were performed with the SPEAR epTau 217 Development Assay Kit and the SPEAR UltraDetect™ Reaction Kit (Catalog#: SPR90024) according to assay manuals, in a 384-well plate format with a semi-automated workflow on the FAST machine, and qPCR readouts were acquired using the QuantStudio instrument. Simoa pTau 217 measurements were performed using the Simoa® ALZpath p-Tau 217 Advantage PLUS Reagent Kit (Quanterix, Catalog#. 104570) on the HD-X platform at the Biofluid Biomarker Laboratory, Department of Psychiatry, University of Pittsburgh School of Medicine, with 100 µL of 3-fold diluted clarified plasma prepared by centrifugation at 4000×g for 10 minutes. For both assays, all samples were analyzed in duplicate, and researchers remained blinded to participant information until completion of data acquisition.

##### *1.10.8 Specimen collection and pTau 217 bio-variability with MGH longitudinal cohort*

Banked plasma samples were selected from the Massachusetts General Hospital Institute for Neurodegenerative Diseases (MIND) Biobank and consisted of samples collected between June 2019 and November 2020 to evaluate the short-term stability of blood biomarkers of neurodegenerative disease. The study population was actively recruited to represent the general healthy adult population, aged 40 or older, without a history of neurodegenerative disease. Participants reported underlying conditions and concomitant medication in a formalized medical history questionnaire conducted by a licensed clinician and participants with an unstable chronic condition or an acute infection were excluded. No acute illness or medication changes occurred during the sample collection window. The study was approved by the Institutional Review Board of Mass General Brigham (#2019P000878) and all study participants provided written informed consent for use of their samples in biomarker research.

Plasma samples were obtained from 20 healthy study participants (10 female/10 male) aged 40.3 to 72.7 years (mean ± SD: 56 ± 9.4 years). Participants were predominantly Caucasian ($n$ = 17; 85%), non-Latinx ($n$ = 19; 95%), and highly educated (>16 years of education: $n$ = 15; 75%). None of the participants had a history of acute infection within the last 90 days or an unstable medical condition (per self-report). EDTA plasma was collected at three visits, each one week apart. Two participants with missing samples from one or more visits were excluded, resulting in 18 participants included in the analysis.

epTau217 and pTau 217 concentrations were measured using the SPEAR and Simoa ALZpath assays respectively to evaluate assay precision across visits. SPEAR epTau 217 measurements were performed using the SPEAR epTau 217 Development Assay Kit and the SPEAR Ultra-Detect™ Reaction Kit (Catalog#: SPR90024) according to the assay manual. Clarified plasma was prepared by centrifugation of thawed EDTA plasma at 10,000×g for 5 minutes, and 1 μL of 4-fold diluted plasma was analyzed in duplicate in a 384-well plate format with a semi-automated work-flow on the FAST machine. qPCR readouts were acquired using QuantStudio instruments.

Simoa pTau 217 measurements were performed using Simoa® ALZpath p-Tau 217 Advantage PLUS Reagent Kit(Quanterix, Catalog#: 104570) on the HD-X platform at the CTRU Biomarker Core at MGH. For this assay, clarified plasma was prepared by centrifugation at 10,000×g for 5 minutes, and 100 μL of 3-fold diluted plasma was analyzed in duplicate.

##### *1.10.9 Specimen collection and epTau 217 measurement with amyloid status confirmed by CSF AD biomarkers at MGH*

Plasma samples ($n$ = 88; participants aged 52–89 years), were selected from the Mass General Institute for Neurodegenerative Disease (MIND) and LifeSPAN biorepositories at MGH. Samples were collected at the time of a diagnostic or research lumbar puncture performed in the Department of Neurology at MGH. Clinical diagnoses were established by chart review by a senior neurologist (SEA), following the 2011 National Institute on Aging–Alzheimer's Association diagnostic criteria for Alzheimer's disease (AD) and mild cognitive impairment (MCI) due to AD[6,7], and AD diagnoses were confirmed using CSF AD biomarkers at the time of plasma sampling.

Amyloid status was verified using CSF biomarkers measured either by the MIND Biomarker Core (Aβ42/40 ratio, pTau181, and total Tau immunoassays; Euroimmun, Lübeck, Germany) or by Athena Diagnostics (ADmark Aβ42, pTau181, and total Tau; Marlboro, MA). Participants with Euroimmun biomarkers were classified as AD if they exhibited a low CSF Aβ42/40 ratio (<0.082), increased pTau181 (> 41.8 pg/mL), and total Tau (> 280 pg/mL). Participants with Athena ADmark biomarkers were classified as AD if the Aβ42/total Tau Index (ATI)[8], calculated as:

$$ATI = \frac{A\beta 42}{240 + (1.18 \times total\ Tau)} \quad \mathbf{(1.10)}$$

below 0.814. Detailed demographic information is provided in **Table S9**.

epTau 217 and pTau 217 concentrations were measured using both the SPEAR epTau 217 Development Kit and the Simoa ALZpath pTau-217 assay. SPEAR measurements were performed using 1 μL of 4-fold diluted clarified plasma, prepared by centrifugation of thawed EDTA plasma at 10,000×g for 5 minutes. Assays were conducted on a 384-well plate format with semi-automated FAST workflow. Simoa pTau-217 measurements were performed at the CTRU Biomarker Core at MGH using Simoa® ALZpath p-Tau 217 Advantage PLUS Reagent Kit (Quanterix, Catalog#:

104570) on the HD-X platform. 100 µL of 3-fold diluted clarified plasma was analyzed in duplicate, prepared by centrifugation of thawed EDTA plasma at 10,000×g for 5 minutes. For both assays, all plasma samples remained deidentified until completion of data acquisition.

##### *1.10.10 Specimen collection and biomaker measurement for HCP-CoBRA Cohort*

A subgroup ($n$ = 113) of the community-based cohort with amyloid PET confirmation was recruited through the Human Connectome Project – Connectomics in Brain Aging (HCP-CoBRA), comprising both cognitively normal and cognitively impaired individuals from Pittsburgh, Pennsylvania, USA, aged 50–89 years. Recruitment sources included the University of Pittsburgh Alzheimer's Disease Research Center (Pitt-ADRC), the Pitt + Me web portal (primarily used to recruit non-college educated Black/African American and non-Hispanic White individuals), as well as active links and word of mouth.

Participants underwent comprehensive clinical and cognitive assessments over three days. Neuroimaging protocols included functional magnetic resonance imaging (fMRI) tasks performed under fasting and non-fasting conditions, magnetoencephalography (MEG), and [11C] Pittsburgh Compound B (PiB) positron emission tomography (PET) imaging for brain Aβ plaques, conducted one week after the final MRI session. Neuropsychological testing followed the Pittsburgh ADRC protocol and included the Montreal Cognitive Assessment (MoCA), verbal fluency, a 30-item visual naming test, Trail Making, verbal free recall, and the Rey–Osterrieth Complex test. Demographic information, including self-identified race, sex, and education, was also collected.

Whole blood was collected in 10 mL K2-EDTA tubes (BD Biosciences, Catalog#: 366643) and centrifuged within 2 hours at 2000×g for 10 minutes at 4 °C to isolate plasma. Plasma aliquots were stored at −80 °C until analysis.

Amyloid (A) status was determined from global [11C] PiB standardized uptake value ratios (SUVR), calculated by volume-weighted averaging across nine composite regions (anterior cingulate, posterior cingulate, insula, superior frontal cortex, orbitofrontal cortex, lateral temporal cortex, parietal cortex, precuneus, and ventral striatum). Participants were classified as amyloid-positive (A+) if SUVR > 1.346. Neurodegeneration (N) status was defined using cortical thickness (CT) measures from MRI. An Alzheimer's disease (AD)-signature composite CT index was calculated from a surface-area weighted average of mean CT across four FreeSurfer[9] regions of interest (entorhinal cortex, inferior and middle temporal gyrus, and fusiform). The study protocol was approved by the University of Pittsburgh Institutional Review Board (STUDY19100015C). Written informed consent was obtained from all participants in accordance with the ethical standards of the University's Human Research Protection Office. Details of the cohort demographics are provided in **Table S10**.

epTau 217 or pTau 217 concentrations were measured using both the SPEAR and Lumipulse assays at the Biofluid Biomarker Laboratory, Department of Psychiatry, University of Pittsburgh School of Medicine. Plasma samples were thawed at room temperature and centrifuged at 4000×g for 10 minutes to remove particulates. SPEAR measurements were conducted using 1 µL of 4-fold diluted clarified plasma. Assays were performed with the SPEAR epTau 217 Development Assay Kit and the SPEAR UltraDetect™ Reaction Kit (Catalog#: SPR90024) according to assay manual, in a 384-well plate format with a semi-automated workflow on the FAST machine. All samples were assayed in duplicate, and qPCR readouts were acquired using QuantStudio instruments. Lumipulse pTau-217 measurements were performed using commercially available Lumipulse® G pTau 217 Plasma assay on the Lumipulse G1200 platform (Fujirebio; Catalog No. 81471) in singlicate from 100 µL of neat, clarified plasma.

For all assays, researchers remained blinded to participant information until completion of data acquisition.

##### *1.11 Statistical Analysis*

The limit of detection (LOD) was defined as 2.5 times the standard deviation (SD) of the blank plus the mean of the blank. The lower limit of quantification (LLOQ) was defined as the lowest calibrator concentration achieving 80–120% standard recovery and ≤ 20% coefficient of variation (CV) of interpolated concentrations derived from the logistic fit. The functional LLOQ (fLLOQ) was calculated by multiplying the LLOQ by the minimum required dilution.

Concordance in comparative studies was assessed using Spearman's rank correlation coefficient (r) and simple linear regression, performed in GraphPad Prism. Intraplate precision was defined as the %CV within a run, calculated from two technical replicates.

The discriminatory performance of plasma and serum biomarkers for amyloid status was evaluated by calculating the area under the receiver operating characteristic curve (AUC) using GraphPad Prism, with assay cutoffs determined at maximal accuracy. Fold changes between amyloid-positive and amyloid-negative groups were calculated as the ratio of their median concentrations (A+/A−).

### 2 Supplementary Text

#### *2.1 Determinants of analytical sensitivity in separation-free immunoassays*

##### *2.1.1 Overview and Definitions*

In this section we aim to provide a theoretical framework to understand the theoretically highest achievable sensitivity of separation-free, homogeneous protein-detection assays in different formats. We specifically focus on competition-, allostery- and proximity-based signal generation mechanisms and highlight how their sensitivities are limited by the $K_D$ of the antibodies (or other binders) used. Against this backdrop, we analyze how a two-step kinetic proofreading mechanism can substantially improve sensitivity even when the affinities of the antibodies are suboptimal.

We define analytical sensitivity in terms of the limit of detection (LOD), which is the lowest analyte concentration that produces a signal statistically distinguishable from the background. In this document, the term 'signal' ($S$) refers to the readout of the analytical instrument used in the assay or another quantity that is proportional to such readout (e.g., fluorescence value measured by a fluorometer, concentration of a fluorescence-generating enzyme, absolute amplicon copy number measured by a qPCR instrument). The readout is often non-zero even in the absence of the analyte. We refer to such readout as 'background signal' ($S_{BG}$). We refer to the readout above background signal in the presence of analyte (denoted as A) as 'true signal' ($S_{True}$). Both $S$ and $S_{True}$ are functions of analyte concentration $C_A$ and by definition:

$$S(C_A) = S_{True}(C_A) + S_{BG}$$

In reality, both background and true signals are always measured with noise or variation. While the noise has many contributing factors, each contributing with a different magnitude, in this document, for simplicity, we assume the noise is proportional to the signal and refer to this signal-to-noise ratio as $1/\sigma$. Since the vast majority of sensitive protein-detection assays are gain-of-signal assays, the LOD can be understood as the lowest $C_A$ that generates a signal that rises above the combined noises of background signal and true signal. In other words, the LOD is defined as the lowest $C_A$ that satisfies this equation (**Fig. S8a**):

$$(1-\sigma)\cdot S(C_A) > (1+\sigma)\cdot S_{BG}$$

In practice, depending on the required statistical significance in the use case, $\sigma$ is usually 1 to 3 times the experimentally determined coefficient of variation (CV) of the readout. Since in gain-of-signal assays $S(C_A)$ monotonically increases (at least when $C_A$ is near LOD), we have:

$$(1-\sigma)\cdot S(\text{LOD}) = (1+\sigma)\cdot S_{BG}$$

or:

$$S(\text{LOD}) = \frac{1+\sigma}{1-\sigma}\cdot S_{BG} \quad \textbf{(2.1)}$$

Considering

$$S(\text{LOD}) = S_{True}(\text{LOD}) + S_{BG}$$

An equivalent expression of LOD is:

$$S_{\mathrm{True}}(\mathrm{LOD}) = S(\mathrm{LOD}) - S_{\mathrm{BG}} = \frac{2\sigma}{1-\sigma} \cdot S_{\mathrm{BG}} \tag{2.2}$$

In other words, LOD can be mathematically defined as $S^{-1}\left(\frac{1+\sigma}{1-\sigma} \cdot S_{\mathrm{BG}}\right)$ or $S_{\mathrm{True}}{}^{-1}\left(\frac{2\sigma}{1-\sigma} \cdot S_{\mathrm{BG}}\right)$. In the following analyses we will use **Eq. 2.1** or **2.2** to calculate LOD, whichever is more convenient.

##### *2.1.2 LOD for competition-based assays*

A popular form of homogeneous protein detection assays is achieved through competition between the free-floating analyte in the sample and a 'decoy', which is a moiety structurally similar to the analyte, and linked to a signal-generating molecule (e.g., an enzyme such as HRP, shown as 'E' in **Fig. S8b**). Both the analyte and the decoy compete for the binder to the analyte (e.g., an antibody, shown as 'B' in **Fig. S8b**). In the absence of the analyte, the binder binds to the decoy, blocking the active site of E or otherwise blunting its signal. The analyte ('A' in **Fig. S8b**) can bind to the binder, depleting the pool of the binder that can bind to decoy, thus liberating the signal-generating molecule, resulting in higher signal. This concept is used in assay format such as Cloned Enzyme Donor Immunoassay (CEDIA)[10], Enzyme Multiplied Immunoassay Technique (EMIT)[11] and Fluorescence Polarization Immunoassay (FPIA)[12].

If the system is allowed to reach equilibrium, the equilibrium concentration of free A, B, E, B-A complex, and B-E complex (written as $[\mathrm{A}]_{\mathrm{eq}}$, $[\mathrm{B}]_{\mathrm{eq}}$, $[\mathrm{E}]_{\mathrm{eq}}$, $[\text{B-A}]_{\mathrm{eq}}$ and $[\text{B-E}]_{\mathrm{eq}}$, respectively) can be obtained by solving the following the equation set:

$$\frac{[\mathrm{B}]_{\mathrm{eq}}[\mathrm{A}]_{\mathrm{eq}}}{[\text{B-A}]_{\mathrm{eq}}} = K_{\mathrm{D}}$$

$$\frac{[\mathrm{B}]_{\mathrm{eq}}[\mathrm{E}]_{\mathrm{eq}}}{[\text{B-E}]_{\mathrm{eq}}} = K_{\mathrm{DE}}$$

$$[\mathrm{A}]_{\mathrm{eq}} + [\text{B-A}]_{\mathrm{eq}} = C_{\mathrm{A}}$$

$$[\mathrm{B}]_{\mathrm{eq}} + [\text{B-E}]_{\mathrm{eq}} + [\text{B-A}]_{\mathrm{eq}} = C_{\mathrm{B}}$$

$$[\mathrm{E}]_{\mathrm{eq}} + [\text{B-E}]_{\mathrm{eq}} = C_{\mathrm{E}}$$

where $K_{\mathrm{D}}$ is the dissociation constant of the binding between A and B; $K_{\mathrm{DE}}$ is the dissociation constant of the binding between A and E; $C_{\mathrm{A}}$, $C_{\mathrm{B}}$ and $C_{\mathrm{E}}$ are the total concentration of A, B and E, respectively. Since in this system the signal is proportional to the concentration of free E, we can replace $S$ in **Eq. 2.1** with $[\mathrm{E}]_{\mathrm{eq}}$, replace $S_{\mathrm{BG}}$ in **Eq. 2.1** with $[\mathrm{E}]_{\mathrm{eq}}|_{C_{\mathrm{A}}=0}$ and solve LOD when the total concentrations and dissociation constants are known.

For example, if $K_{\mathrm{D}}$ and $K_{\mathrm{DE}}$ are both 200 pM, the total concentrations of B and E are 200 pM and 50 pM, respectively, and the signal-to-noise ratio is 10 (i.e., $\sigma = 0.1$), it can be seen from **Fig. S9a** that only when $C_{\mathrm{A}}$ exceeds 212 pM the signal exceeds background signal beyond noise, i.e., LOD = 212 pM. The LOD can be marginally improved to 187.9 pM when the centration of E is lowered to 2 pM (**Fig. S9b-c**), primarily due to lowering of background. The LOD can also be marginally improved to 187.8 pM by using higher concentration of B (300 pM), but its further increase causes the LOD to increase (**Fig. S9d-f**).

In any case, one cannot achieve LOD substantially below the $K_\mathrm{D}$ using competition-based assays. Intuitively, this is because the signal only changes by a meaningful magnitude when the available pool of binder is meaningfully depleted by the analyte, which can only happen when the concentration of the analyte is at least comparable to $K_\mathrm{D}$.

##### *2.1.3 LOD for allostery-based assays*

An alternative strategy to achieve homogeneous protein detection is through allosteric binding. In fact, naturally existing detectors of protein often use allostery to transduce signals, where binding of the input protein (equivalent to an analyte) induces conformational (i.e., allosteric) change of the detector and alters the rate of a downstream process (e.g., downstream protein phosphorylation if the detector is a kinase, or transcription of downstream genes if the biosensor is a transcription factor).

A general form of allosteric protein detector can be depicted in **Fig. S8c**, where the detector B exists in two conformations: the active form $\mathrm{B^{act}}$ and the inactive form $\mathrm{B^{in}}$. We refer to the equilibrium constant between these two states as $K_1$. A key principle of such allosteric protein detector is that the active form generates much higher signal than the inactive form. In the most ideal case, the inactive form does not generate signal at all. Another key principle is that the active form binds the analyte A much more strongly than the inactive form. Ideally, the inactive form does not bind the analyte at all. We refer to the dissociation constant between the active form $\mathrm{B^{act}}$ and A as $K_\mathrm{D}$, and analyze these ideal cases to understand the limitation of allostery-based assays.

Under these conditions, the equilibrium concentration of each species can be solved with the following equation set:

$$\frac{[\mathrm{B^{act}}]_\mathrm{eq}}{[\mathrm{B^{in}}]_\mathrm{eq}} = K_1$$

$$\frac{[\mathrm{A}]_\mathrm{eq}[\mathrm{B^{act}}]_\mathrm{eq}}{[\mathrm{A\text{-}B^{act}}]_\mathrm{eq}} = K_\mathrm{D} \qquad \mathbf{(2.3)}$$

$$\left[\mathrm{B^{in}}\right]_\mathrm{eq} + \left[\mathrm{B^{act}}\right]_\mathrm{eq} + \left[\mathrm{A\text{-}B^{act}}\right]_\mathrm{eq} = C_\mathrm{B}$$

$$[\mathrm{A}]_\mathrm{eq} + \left[\mathrm{A\text{-}B^{act}}\right]_\mathrm{eq} = C_\mathrm{A}$$

The relative simplicity of this equation set allows us to obtain analytical solution of LOD in a simple form. Since only the $\mathrm{B^{act}}$ is signal-generating, regardless of whether it is bound to A, we can regard $\left[\mathrm{B^{act}}\right]_\mathrm{eq} + \left[\mathrm{A\text{-}B^{act}}\right]_\mathrm{eq}$ as signal $S$. The background signal is simply the signal when $C_\mathrm{A} = 0$, which can be easily calculated:

$$S_\mathrm{BG} = [\mathrm{B^{act}}]_\mathrm{eq}|_{C_\mathrm{A}=0} = \frac{K_1}{1+K_1} \cdot C_\mathrm{B}$$

Using the definition of LOD expressed in **Eq. 2.1**, we can add the following equations to the **Eq. set 2.3**:

$$C_\mathrm{A} = \mathrm{LOD}$$

$$[\mathrm{B^{act}}]_{\mathrm{eq}} + [\mathrm{A\text{-}B^{act}}]_{\mathrm{eq}} = \frac{1+\sigma}{1-\sigma}\cdot\frac{K_1}{1+K_1}\cdot C_\mathrm{B}$$

We can obtain:

$$\mathrm{LOD} = \frac{2\sigma\left[K_\mathrm{D} + C_\mathrm{B}K_1 + K_1K_\mathrm{D} - \left(K_\mathrm{D} + C_\mathrm{B}K_1 + K_1K_\mathrm{D} + 2C_\mathrm{B}{K_1}^2\right)\sigma\right]}{1 - 2(K_1+1)\sigma + (2K_1+1)\sigma^2} \qquad \textbf{(2.4)}$$

From **Eq. 2.4**, it can be proven that LOD as a function of $K_1$ monotonically increases (within the physically meaningful regime, where the LOD condition admits a non-negative solution) and the lower bound of LOD is achieved when $K_1$ approaches zero, and:

$$\min(\mathrm{LOD}) = \lim_{K_1\to 0}\mathrm{LOD} = \frac{2\sigma}{1-\sigma}\cdot K_\mathrm{D}$$

In other words, the allosteric detector should be engineered to favor the inactive state in order to optimize for LOD. When the signal-to-noise ratio approaches infinite (or $\sigma$ approaches 0), LOD approaches the following limit:

$$\lim_{K_1,\sigma\to 0}\mathrm{LOD} = 2\sigma\cdot K_\mathrm{D} \qquad \textbf{(2.5)}$$

Therefore, the lower bound of LOD in an allosteric protein detector is $K_\mathrm{D}$ scaled down by the factor $2\sigma$ (which is usually less than 1). In practice, if CV of the signal is 10%, and $\sigma$ is 1 to 3 times the CV (depending on necessary threshold for statistical confidence in the use case), we can see that even in the most ideal cases, the lower bound of LOD is no lower than 1 order of magnitude below $K_\mathrm{D}$.

There are practical factors that make this modest theoretical lower bound of LOD hard to reach. What was not considered in the analysis above is the number of signal-generating molecules produced per molecule of analyte (i.e., $\frac{[\mathrm{B^{act}}]_{\mathrm{eq}}+[\mathrm{A\text{-}B^{act}}]_{\mathrm{eq}}}{C_\mathrm{A}}$), which can be understood as 'yield' of the analyte. In the analysis above, concentrations are treated as continuous variables. While this treatment is proper when the concentrations are high, when we consider concentrations in the near single molecule-level, they become discrete and Poisson error begins to appear. If one aims to limit Poisson error at 10% or lower, the number of signal-generating molecules $\left([\mathrm{B^{act}}]_{\mathrm{eq}} + [\mathrm{A\text{-}B^{act}}]_{\mathrm{eq}}\right)\cdot V\cdot N_\mathrm{A}$ (where $V$ is the volume of the binding reaction and $N_\mathrm{A}$ is the Avogadro number) needs to be at least 100. In this case, if one wishes to detect 1,000 molecules of analyte, the yield has to be greater than 10%. When $K_1$ approaches zero, the yield decreases as the fraction of the active binder that can bind the analyte decreases.

There are other reasons that preclude arbitrarily lowering $K_1$ and sacrificing yield. For example, as discussed above, the $\mathrm{B^{in}}$ state usually generates background signal to some extent. Therefore, if the yield is too low, the true signal cannot rise above the noise floor of the background. Along similar logic, according to **Eq. 2.4** lowering the concentration of the detector ($C_\mathrm{B}$) also lowers the LOD. However, this also comes at the expense of decreasing yield. In practice, the concentration of the binder must be at least 50 to 100 pM to ensure acceptable yield and to ensure that, in terms of kinetics, binding equilibrium can be achieved in a typical experimental time frame (e.g, within a few hours).

To quantitatively understand the relationships of these factors in realistic ranges, we can first refine the definition of signal and background to the following equations,

$$S = \left([\mathrm{B}^{\mathrm{act}}]_{\mathrm{eq}} + [\mathrm{A}\text{-}\mathrm{B}^{\mathrm{act}}]_{\mathrm{eq}}\right) \cdot E_{\mathrm{act}} + [\mathrm{B}^{\mathrm{in}}]_{\mathrm{eq}} \cdot E_{\mathrm{in}}$$

$$S_{\mathrm{BG}} = \left([\mathrm{B}^{\mathrm{act}}]_{\mathrm{eq}} \cdot E_{\mathrm{act}} + [\mathrm{B}^{\mathrm{in}}]_{\mathrm{eq}} \cdot E_{\mathrm{in}}\right)|_{C_{\mathrm{A}}=0}$$

where $E_{\mathrm{act}}$ and $E_{\mathrm{in}}$ reflect the signal-generating ability of the active and inactive form of the binder, respectively. We can apply a realistic set of parameters (e.g., $K_{\mathrm{D}}$ = 200 pM, $C_{\mathrm{B}}$ = 200 pM, $\sigma$ = 0.1, $E_{\mathrm{in}}/E_{\mathrm{act}}$ = 0.01) to numerically compute the optimal $K_1$ to minimize LOD. As shown in **Fig. S10a – c**, the background can be lowered and the LOD can be improved when one lowers $K_1$ (e.g., from 1 to 0.1, **Fig. S10a,b**). However, when $K_1$ becomes too low (e.g., 0.01, **Fig. S10c**) the sensitivity worsens. There is an optimal $K_1$, with which the optimal LOD is 59.5 pM (**Fig. S10d**).

If the $K_{\mathrm{D}}$ is higher (e.g., raised from 200 pM to 1 nM) the lowest achievable LOD is also proportionally higher (e.g., 282.4 pM) under optimal $K_1$ (**Fig. S10e**). In fact, for any $K_{\mathrm{D}}$ value and any $E_{\mathrm{in}}/E_{\mathrm{act}}$ ratio, one can numerically compute the optimal $K_1$ and its corresponding LOD. As shown in **Fig. S10f**, lowering the $E_{\mathrm{in}}/E_{\mathrm{act}}$ ratio indeed pushes the lowest achievable LOD closer to the limit set out in **Eq. 2.5** (i.e., $2\sigma \cdot K_{\mathrm{D}}$).

From these analyses, it can be seen that since binders of proteins typically have $K_{\mathrm{D}}$ values in the range of 100 pM to 10 nM, the lowest achievable LOD using the mechanism of allosteric signaling is at best 20 pM to 2 nM (assuming $\sigma = 0.1$), several orders of magnitude away from single-molecule level sensitivity (note that 100 molecules in 10 $\mu$L is 17 aM).

The functional form of the lower bound in **Eq. 2.5** admits a clear physical interpretation. The factor $2\sigma$ represents the statistical floor imposed by readout variation: an analyte-induced signal increment must rise above this noise band to register as a true detection. The factor $K_{\mathrm{D}}$ represents the thermodynamic floor imposed by the binder: at analyte concentrations far below $K_{\mathrm{D}}$, the target fails to occupy a meaningful fraction of the binder, regardless of how quiet the readout is. Homogeneous single-binder architectures cannot lower either floor, and the product $2\sigma \cdot K_{\mathrm{D}}$ is the lowest LOD achievable when both are simultaneously saturated. This dual constraint motivates the strategies developed in **Sections 2.1.4** and **2.1.5**: introducing a second, proximity-dependent binder effectively replaces the single-$K_{\mathrm{D}}$ barrier with a product of two $K_{\mathrm{D}}$'s scaled by a local concentration, while a two-step kinetic-proofreading process further suppresses background through rate differentiation between proximity-driven and bimolecular reactions. Together, these strategies lower the achievable LOD by orders of magnitude relative to the bound derived here.

###### *2.1.4 LOD for proximity-based signal generation*

Protein biomarkers often have large surfaces to accommodate multiple antibody-like binders to bind simultaneously. This fact can be exploited to develop proximity-based homogeneous protein detection assays, which operates on a simple principle: when two binders bind the same analyte molecule, they are at much shorter distance from each other than when they are freely diffusing in the solution. So long as the proximity can be converted to an observable readout, an assay can be built. Examples of proximity-based homogeneous assays include Fluorescence Resonance Energy Transfer (FRET)[13], Luminescent Oxygen Channeling Immunoassay (LOCI)[14], Proximity Ligation Assay (PLA)[15] and Proximity Extension Assay (PEA)[16].

Model for an ideal proximity sensor

A relatively simple model to understand the power and limitation of proximity-based separation-free assays is an ideal FRET-like signal generation system shown in **Fig. S11a**, where the binders B and B2 (which can bind to different epitopes of the same analyte, forming a tripartite immunocomplex) carry an 'ideal FRET donor' ($F_D$) and an 'ideal FRET acceptor' ($F_A$), respectively. We refer to the average distance between $F_D$ and $F_A$ in the tripartite immunocomplex as $d$. For illustration, we can assume that when $F_D$ is excited, it transfers energy to $F_A$ only if the distance between the two moieties is shorter than $d$, which is (1) a reasonable binary approximation of the Förster equation considering its steep curve originating from the dependence on the sixth power of $r/R_0$ ($r$ being the distance between $F_D$ and $F_A$, and $R_0$ being the Förster distance) combined with (2) an idealistic assumption that the $R_0$ of the FRET pair perfectly matches $d$. We refer to the dissociation constants of B and B2 as $K_D$ and $K_{D2}$, respectively. Therefore, the signal can be regarded as the concentration of the tripartite immunocomplex:

$$S_{\mathrm{True}}(C_A) = C_A \cdot \frac{C_B}{K_D + C_B} \cdot \frac{C_{B2}}{K_{D2} + C_{B2}}$$

In this model system, the background signal can be generated if a B2 molecule diffuses into the proximal space of a B molecule, defined as a sphere centered at the B molecule and has a radius of $d$ (**Fig. S11b**). Therefore the background signal can be regarded as the concentration of B molecule that happen to be 'accompanied' by a B2 molecule:

$$S_{BG} = C_B \cdot \frac{4}{3}\pi d^3 \cdot C_{B2} \cdot N_A$$

Applying **Eq. 2.2** we have:

$$\mathrm{LOD} \cdot \frac{C_B}{K_D + C_B} \cdot \frac{C_{B2}}{K_{D2} + C_{B2}} = \frac{2\sigma}{1-\sigma} C_B \cdot \frac{4}{3}\pi d^3 \cdot C_{B2} \cdot N_A$$

or

$$\mathrm{LOD} = \frac{2\sigma}{1-\sigma} \cdot (K_D + C_B) \cdot (K_{D2} + C_{B2}) \frac{4}{3}\pi d^3 N_A \qquad \textbf{(2.6)}$$

We can see from **Eq. 2.6** that lower LOD is achieved when lower concentrations of the binders ($C_B$ and $C_{B2}$) are used (lower yield notwithstanding), and we can observe the following lower limit of LOD:

$$\min(\mathrm{LOD}) = \lim_{C_B, C_{B2}, \sigma \to 0} \mathrm{LOD} = 2\sigma \cdot K_D \cdot K_{D2} \cdot \frac{4}{3}\pi d^3 N_A$$

Note that $\frac{1}{\frac{4}{3}\pi d^3 N_A}$ (i.e., the concentration of 1 molecule in the sphere with radius of $d$) can be understood as the local concentration of B2 with respect to B when B2 is brought to proximity of B through both binding to the analyte. We can replace this term with $C_{\mathrm{Local}}$ and rewrite the **Eq. 2.6** as:

$$\min(\mathrm{LOD}) = \lim_{C_B, C_{B2}, \sigma \to 0} \mathrm{LOD} = 2\sigma \cdot K_D \cdot \frac{K_{D2}}{C_{\mathrm{Local}}} \qquad \textbf{(2.7)}$$

By comparing **Eqs. 2.5** and **2.7**, we can see that by using an additional binder, the LOD can be scaled by a factor of $\frac{K_{\mathrm{D2}}}{C_{\mathrm{Local}}}$. This scaling factor depends on the distance $d$. If $d$ = 30 nm (which is reasonable considering the typical sizes of antibodies and protein analytes) then $C_{\mathrm{Local}}$ can be as high as 15 μM. If $K_{\mathrm{D2}}$ is 100 pM, this scaling factor can be as low as $6.7\times10^{-6}$. Even if $d$ = 200 nm, then $C_{\mathrm{Local}}$ can still be as high as 50 nM. And the scaling factor can be $2\times10^{-3}$. This represents an opportunity to improve LOD by several orders of magnitude.

Model for proximity amplicon-generation assays

To implement the 'ideal FRET-like' assay described above and achieve single-molecule level sensitivity, one needs to be able to observe signals emitted by a few $F_A$ molecules and meet all the assumptions that were made above, which is an engineering challenge that has not been met yet. However, to recapitulate this proximity effect and the enhancement of sensitivity, (bio)chemical reaction can be engineered in a way that high local concentration can promote the generation of a physical product. If the product can be readily detected at single-molecule level (such as a DNA amplicon), the lower theoretical LOD afforded by the proximity effect can be combined with simple and sensitive amplicon detection such as qPCR and NGS. This is indeed the bases of proximity amplicon-generation (PAG) assays such as PLA and PEA.

To provide a general model for PAG assays, we consider that the antibodies B and B2 are modified with two reactants. When both antibodies bind to the analyte A, the two reactants within the immunocomplex can react with an apparent first-order rate constant $k_{\mathrm{uni}}$ (the subscript 'uni' signifies uni-molecular reaction, **Fig. S11c**-left). The background signal in such assays can arise because a pair of reactants may still undergo the bi-molecular reaction in the absence of the analyte, with an apparent second-order rate constant $k_{\mathrm{bi}}$ (the subscript 'bi' signifies bimolecular reaction, **Fig. S11c**-right).

PAG assays are usually carried out by first incubating the modified antibodies and the analyte without the enzyme (or a key co-factor that completes the amplicon-generation) to reach equilibrium, after which the enzyme (or the key co-factor) is added. We first consider the systems where the spontaneous dissociation of the antibodies from the analyte happens very slowly and can be ignored in the time scale of the amplicon generation reaction. We will later consider cases where this assumption does not hold. At any time point, true signal $S_{\mathrm{True}}$ is represented by the concentration of the amplifiable reaction product (i.e., the amplicon), which we refer to as A-B-B2$^*$. Namely,

$$S_{\mathrm{True}}(t) = [\text{A-B-B2}^*]_t \qquad \textbf{(2.8)}$$

At any time point the rate of true signal generation is the concentration of 'unreacted' tripartite immunocomplex times $k_{\mathrm{uni}}$:

$$\frac{d[\text{A-B-B2}^*]_t}{dt} = [\text{A-B-B2}]_t \cdot k_{\mathrm{uni}} \qquad \textbf{(2.8.1)}$$

This reaction also consumes 'unreacted' tripartite immunocomplex:

$$\frac{d[\text{A-B-B2}]_t}{dt} = -[\text{A-B-B2}]_t \cdot k_{\mathrm{uni}} \qquad \textbf{(2.9)}$$

When the antibody-analyte binding achieves equilibrium, the concentration of the tripartite immunocomplex is $C_{\mathrm{A}} \cdot \frac{C_{\mathrm{B}}}{K_{\mathrm{D}}+C_{\mathrm{B}}} \cdot \frac{C_{\mathrm{B2}}}{K_{\mathrm{D2}}+C_{\mathrm{B2}}}$, which is also the initial concentration of unreacted tripartite immunocomplex:

$$[\text{A-B-B2}]|_{t=0} = C_{\mathrm{A}} \cdot \frac{C_{\mathrm{B}}}{K_{\mathrm{D}} + C_{\mathrm{B}}} \cdot \frac{C_{\mathrm{B2}}}{K_{\mathrm{D2}} + C_{\mathrm{B2}}} \tag{2.9.1}$$

The initial concentration of the reaction product (i.e., the amplicon) is zero:

$$[\text{A-B-B2}^*]|_{t=0} = 0 \tag{2.9.2}$$

Integration of the **Eqs. 2.8.1 – 2.9** with the initial conditions above results in the classic single exponential decay kinetics:

$$S_{\mathrm{True}}(t) = [\text{A-B-B2}^*]_t = C_{\mathrm{A}} \cdot \frac{C_{\mathrm{B}}}{K_{\mathrm{D}} + C_{\mathrm{B}}} \cdot \frac{C_{\mathrm{B2}}}{K_{\mathrm{D2}} + C_{\mathrm{B2}}} (1 - e^{-k_{\mathrm{uni}} \cdot t}) \tag{2.10}$$

Now we consider the background signal which is represented by the amplicon formed in the absence of the analyte. We refer to this product as B-B2*:

$$S_{\mathrm{BG}}(t) = [\text{B-B2}^*]_t \tag{2.10.1}$$

We can track the formation of B-B2* from the perspective of B. Since only a tiny fraction of B2 is converted to B-B2* (femtomolar-level B-B2* compared to nanomolar-level $C_{\mathrm{B2}}$), we can view the concentration of B2 as constant, therefore:

$$\frac{d[\mathrm{B}]_t}{dt} = -[\mathrm{B}]_t \cdot C_{\mathrm{B2}} \cdot k_{\mathrm{bi}} \tag{2.10.2}$$

$$\frac{d[\text{B-B2}^*]_t}{dt} = [\mathrm{B}]_t \cdot C_{\mathrm{B2}} \cdot k_{\mathrm{bi}} \tag{2.10.3}$$

with the following initial conditions:

$$[\mathrm{B}]|_{t=0} = C_{\mathrm{B}} \tag{2.10.4}$$

$$[\text{B-B2}^*]|_{t=0} = 0 \tag{2.10.5}$$

We can obtain:

$$S_{\mathrm{BG}}(t) = [\text{B-B2}^*]_t = C_{\mathrm{B}} \cdot (1 - e^{-C_{\mathrm{B2}} \cdot k_{\mathrm{bi}} \cdot t}) \tag{2.10.6}$$

By applying the definition of LOD in **Eq. 2.2**, we have

$$\mathrm{LOD}(t) \cdot \frac{C_{\mathrm{B}}}{K_{\mathrm{D}} + C_{\mathrm{B}}} \cdot \frac{C_{\mathrm{B2}}}{K_{\mathrm{D2}} + C_{\mathrm{B2}}} (1 - e^{-k_{\mathrm{uni}} \cdot t}) = \frac{2\sigma}{1-\sigma} C_{\mathrm{B}} \cdot (1 - e^{-C_{\mathrm{B2}} \cdot k_{\mathrm{bi}} \cdot t})$$

Therefore:

$$\mathrm{LOD}(t) = \frac{2\sigma}{1-\sigma} \cdot (K_{\mathrm{D}} + C_{\mathrm{B}}) \cdot \frac{K_{\mathrm{D2}} + C_{\mathrm{B2}}}{C_{\mathrm{B2}}} \cdot \frac{1 - e^{-C_{\mathrm{B2}} \cdot k_{\mathrm{bi}} \cdot t}}{1 - e^{-k_{\mathrm{uni}} \cdot t}} \tag{2.11}$$

It can be proved that LOD monotonically increases with respect to $t$, $C_{\mathrm{B}}$ and $C_{\mathrm{B2}}$ and reaches its minimum value when these parameters all approach zero (again, the sacrifice of yield notwithstanding). When $\sigma$ is small, we can obtain the lower bound of LOD:

$$\min(\text{LOD}) = \lim_{C_{\mathbf{B}},C_{\mathbf{B2}},t,\sigma\to 0} \text{LOD}(t) = 2\sigma \cdot K_{\text{D}} \cdot \frac{K_{\text{D2}}}{\frac{k_{\text{uni}}}{k_{\text{bi}}}} \quad \textbf{(2.12)}$$

The term $\frac{k_{\text{uni}}}{k_{\text{bi}}}$ has a unit of molar and can be understood as how fast the reaction is accelerated due to the proximity, so we can understand it as 'functional local concentration' and refer to it as $C'_{\text{Local}}$.

There are several key differences between $C_{\text{Local}}$ and $C'_{\text{Local}}$. $C_{\text{Local}}$ is fundamentally defined by geometry while $C'_{\text{Local}}$ is defined by rate constants. More importantly, while $C_{\text{Local}}$ can only be estimated based on structural models, $C'_{\text{Local}}$ can be experimentally measured by measuring $k_{\text{uni}}$ and $k_{\text{bi}}$. Our experimental data show that $C'_{\text{Local}}$ in antibody-based proximity assays is on the order of 2 $\mu$M (see **Section 2.1.7**, **Eq. 2.20**, and **Fig. S12** for the experimental estimation).

Thus, we can rewrite **Eq. 2.12** as:

$$\min(\text{LOD}) = \lim_{C_{\text{B}},C_{\text{B2}},t,\sigma\to 0} \text{LOD}(t) = 2\sigma \cdot K_{\text{D}} \cdot \frac{K_{\text{D2}}}{C'_{\text{Local}}} \quad \textbf{(2.13)}$$

Noting the similarity between equations **2.7** and **2.13**, we can similarly state that the lower bound of LOD in a proximity-based amplicon generation assay is the $K_{\text{D}}$ of one binder, scaled down by $2\sigma$, and scaled down again by $\frac{K_{\text{D2}}}{C'_{\text{Local}}}$. If $K_{\text{D2}}$ is 1 nM and $C'_{\text{Local}}$ is 2 $\mu$M, the scaling factor can be as low as 1/2,000.

For example, applying **Eq. 2.13** with $\sigma = 0.1$ and the experimentally measured $C'_{\text{Local}} = 2\ \mu\text{M}$, the LOD reaches ~1 fM (~600 molecules in 1 $\mu$L) for high-affinity antibody pairs ($K_{\text{D}} = K_{\text{D2}} = 100$ pM) — a substantial improvement over the single-binder allostery bound of ~20 pM at the same affinity, and approaching the threshold of single-molecule sensitivity. However, the quadratic dependence on $K_{\text{D}}$ again means that with antibody pairs of more typical affinity ($K_{\text{D}} = K_{\text{D2}} = 1$ nM), the LOD rises to ~100 fM (~60,000 molecules in 1 $\mu$L), and with weaker pairs ($K_{\text{D}} = K_{\text{D2}} = 10$ nM) to ~10 pM (~6 × $10^6$ molecules in 1 $\mu$L) — well short of single-molecule sensitivity. Achieving the ~100-molecule threshold across the broader range of realistic antibody affinities therefore requires a mechanism that further suppresses background beyond what proximity alone provides, motivating the kinetic-proofreading approach discussed in **Section 2.1.5** and **2.1.6**.

##### *2.1.5 Enhancement of rate differentiation by two-step kinetic proofreading*

Before analyzing how two-step kinetic proofreading can improve sensitivity of immunoassays, we discuss a general property of this process. We consider a precursor **P** that can be converted to wanted product $\mathbf{P_W}$ and, independently, unwanted product $\mathbf{P_U}$ with first-order rate constants $\boldsymbol{k_W}$ and $\boldsymbol{k_U}$, respectively:

$$\mathbf{P} \xrightarrow{k_{\mathrm{W}}} \mathbf{P_W}$$

$$\mathbf{P} \xrightarrow{k_{\mathrm{U}}} \mathbf{P_U}$$

The kinetics of the formation of $\mathbf{P_W}$ can be described in the following differential equation set:

$$\begin{cases} \frac{d[\mathbf{P}]_t}{dt} = -[\mathbf{P}] \cdot \boldsymbol{k}_{\mathrm{W}} \\ \frac{d[\mathbf{P}_{\mathbf{W}}]}{dt} = [\mathbf{P}] \cdot \boldsymbol{k}_{\mathrm{W}} \end{cases}$$

with initial condition:

$$[\mathbf{P}]|_{t=0} = \boldsymbol{C}_{\mathbf{P}}$$

$$[\mathbf{P}_{\mathbf{W}}]|_{t=0} = 0$$

Similarly, the kinetics of the formation of $\mathbf{P}_{\mathrm{U}}$ can be described in the following differential equation set:

$$\begin{cases} \frac{d[\mathbf{P}]}{dt} = -[\mathbf{P}] \cdot \boldsymbol{k}_{\mathrm{U}} \\ \frac{d[\mathbf{P}_{\mathbf{U}}]}{dt} = [\mathbf{P}] \cdot \boldsymbol{k}_{\mathrm{U}} \end{cases}$$

with initial conditions:

$$[\mathbf{P}]|_{t=0} = \boldsymbol{C}_{\mathbf{P}}$$

$$[\mathbf{P}_{\mathbf{U}}]|_{t=0} = 0$$

Integration of these differential equation sets yields:

$$[\mathbf{P}_{\mathbf{W}}] = \boldsymbol{C}_{\mathbf{P}}(1 - e^{-\boldsymbol{k}_{\mathrm{W}} \cdot t})$$

$$[\mathbf{P}_{\mathbf{U}}] = \boldsymbol{C}_{\mathbf{P}}(1 - e^{-\boldsymbol{k}_{\mathrm{U}} \cdot t})$$

Thus, the ratio of $[\mathbf{P}_{\mathbf{W}}]$ to $[\mathbf{P}_{\mathbf{U}}]$, written as $R$, can be calculated:

$$R = \frac{[\mathbf{P}_{\mathbf{W}}]}{[\mathbf{P}_{\mathbf{U}}]} = \frac{1 - e^{-\boldsymbol{k}_{\mathrm{W}} \cdot t}}{1 - e^{-\boldsymbol{k}_{\mathrm{U}} \cdot t}}$$

$R$ as a function of $t$ monotonically decreases when $\boldsymbol{k}_{\mathrm{W}} > \boldsymbol{k}_{\mathrm{U}}$ and approaches its maximum value as $t$ approaches 0. By applying Taylor series expansion of $e^{-\boldsymbol{k}_{\mathrm{W}} \cdot t}$ and $e^{-\boldsymbol{k}_{\mathrm{U}} \cdot t}$ and ignoring higher-order terms, it can also be deduced that:

$$\max(R) = \lim_{t \to 0} R = \frac{\boldsymbol{k}_{\mathrm{W}}}{\boldsymbol{k}_{\mathrm{U}}}$$

We call this ratio $\frac{\boldsymbol{k}_{\mathrm{W}}}{\boldsymbol{k}_{\mathrm{U}}}$ the characteristic ratio of this system and write it as $R_{\mathrm{C}}$.

When intermediates $\mathbf{P}^{\mathbf{0}}_{\mathbf{W}}$ and $\mathbf{P}^{\mathbf{0}}_{\mathbf{U}}$ are introduced (with the "0" superscript denoting the half-extended intermediate and "*" the fully extended final product, as described in **Fig. 1a** of the main text), the reactions can be described as:

$$\mathbf{P} \xrightarrow{\boldsymbol{k}_{\mathrm{W}}} \mathbf{P}^{\mathbf{0}}_{\mathbf{W}} \xrightarrow{\boldsymbol{k}_{\mathrm{W}}} \mathbf{P}^{*}_{\mathbf{W}}$$

$$\mathbf{P} \xrightarrow{\boldsymbol{k}_{\mathrm{U}}} \mathbf{P}^{\mathbf{0}}_{\mathbf{U}} \xrightarrow{\boldsymbol{k}_{\mathrm{U}}} \mathbf{P}^{*}_{\mathbf{U}}$$

The kinetics of the formation and consumption (when applicable) of $\mathbf{P}^{\mathbf{0}}_{\mathbf{W}}$ and $\mathbf{P}^{*}_{\mathbf{W}}$ can be described by the following differential equation set:

$$\begin{cases} \dfrac{d[\mathbf{P}]}{dt} = -[\mathbf{P}] \cdot \boldsymbol{k}_{\mathrm{W}} \\ \dfrac{d[\mathbf{P}_{\mathrm{W}}^{\mathbf{0}}]}{dt} = [\mathbf{P}] \cdot \boldsymbol{k}_{\mathrm{W}} - [\mathbf{P}_{\mathrm{W}}^{\mathbf{0}}] \cdot \boldsymbol{k}_{\mathrm{W}} \\ \dfrac{d[\mathbf{P}_{\mathrm{W}}^{*}]}{dt} = [\mathbf{P}_{\mathrm{W}}^{\mathbf{0}}] \cdot \boldsymbol{k}_{\mathrm{W}} \end{cases}$$

with initial conditions:

$$[\mathbf{P}]|_{t=0} = \boldsymbol{C}_{\mathbf{P}}$$

$$[\mathbf{P}_{\mathrm{W}}^{\mathbf{0}}]|_{t=0} = 0$$

$$[\mathbf{P}_{\mathrm{W}}^{*}]|_{t=0} = 0$$

The kinetics of the formation and consumption (when applicable) of $\mathbf{P}_{\mathrm{U}}^{\mathbf{0}}$ and $\mathbf{P}_{\mathrm{U}}^{*}$ can be similarly described by replacing subscript 'W' with 'U'.

Integration of these differential equation sets yields:

$$[\mathbf{P}_{\mathrm{W}}^{*}]_t = \boldsymbol{C}_{\mathbf{P}}[1 - (\boldsymbol{k}_{\mathrm{W}} \cdot t + 1)e^{-\boldsymbol{k}_{\mathrm{W}} \cdot t}] \quad \textbf{(2.14)}$$

$$[\mathbf{P}_{\mathrm{U}}^{*}]_t = \boldsymbol{C}_{\mathbf{P}}[1 - (\boldsymbol{k}_{\mathrm{U}} \cdot t + 1)e^{-\boldsymbol{k}_{\mathrm{U}} \cdot t}] \quad \textbf{(2.15)}$$

The ratio $R$ can be calculated as:

$$R = \frac{[\mathbf{P}_{\mathrm{W}}^{*}]}{[\mathbf{P}_{\mathrm{U}}^{*}]} = \frac{1 - (\boldsymbol{k}_{\mathrm{W}} \cdot t + 1)e^{-\boldsymbol{k}_{\mathrm{W}} \cdot t}}{1 - (\boldsymbol{k}_{\mathrm{U}} \cdot t + 1)e^{-\boldsymbol{k}_{\mathrm{U}} \cdot t}}$$

It can also be proven that $R$ as a function of $t$ monotonically decreases when $\boldsymbol{k}_{\mathrm{W}} > \boldsymbol{k}_{\mathrm{U}}$ and approaches its maximum value as $t$ approaches 0.

By applying Taylor series expansion and ignoring higher-order terms, it can be obtained that:

$$\max(R) = \lim_{t \to 0} R = \left(\frac{\boldsymbol{k}_{\mathrm{W}}}{\boldsymbol{k}_{\mathrm{U}}}\right)^2 = {R_{\mathrm{C}}}^2 \quad \textbf{(2.16)}$$

In other words, the maximum $R$ is increased from $\frac{\boldsymbol{k}_{\mathrm{W}}}{\boldsymbol{k}_{\mathrm{U}}}$ to $\left(\frac{\boldsymbol{k}_{\mathrm{W}}}{\boldsymbol{k}_{\mathrm{U}}}\right)^2$.

##### *2.1.6 LOD for SPEAR*

In SPEAR, precursor of the wanted product is the immunocomplex formed by A, B and B2. If the experiment is carried out in a way that DNA polymerase is added after the bindings reach equilibria, the initial concentration of the precursor is $C_{\mathrm{A}} \cdot \frac{C_{\mathrm{B}}}{K_{\mathrm{D}}+C_{\mathrm{B}}} \cdot \frac{C_{\mathrm{B2}}}{K_{\mathrm{D2}}+C_{\mathrm{B2}}}$. The intended intermediate $\mathbf{P}_{\mathrm{W}}^{\mathbf{0}}$ is represented by the once-extended half record (written as A-B-B2$^0$), and the intended product $\mathbf{P}_{\mathrm{W}}^{*}$ is represented by the twice-extended full record (written as A-B-B2$^*$). Its concentration, also representing signal, as a function of time can be described by **Eq. 2.14** where:

$$\boldsymbol{k}_{\mathrm{W}} = k_{\mathrm{uni}}$$

$$\boldsymbol{C}_{\mathbf{P}} = C_{\mathrm{A}} \frac{C_{\mathrm{B}}}{K_{\mathrm{D}} + C_{\mathrm{B}}} \cdot \frac{C_{\mathrm{B2}}}{K_{\mathrm{D2}} + C_{\mathrm{B2}}}$$

(Note that the antibody dissociation is assumed here to be slow relative to the extension reaction, and the precursor pool is not depleted by unbinding; this assumption is relaxed in **Section 2.1.7.**)

In other words:

$$S_{\mathrm{True}}(t) = C_{\mathrm{A}} \frac{C_{\mathrm{B}}}{K_{\mathrm{D}} + C_{\mathrm{B}}} \cdot \frac{C_{\mathrm{B2}}}{K_{\mathrm{D2}} + C_{\mathrm{B2}}} [1 - (k_{\mathrm{uni}} \cdot t + 1) e^{-k_{\mathrm{uni}} \cdot t}] \quad \textbf{(2.17)}$$

For background signal, $\mathbf{P}_{\mathrm{U}}^{*}$ is represented by $\mathrm{B}^{*}$, the twice-extended product of B, which is generated through transient collisional encounters with free B2 (which serves only as a diffusing template, not as a stable binding partner), in the absence of the analyte. Its concentration as a function of time can be described by **Eq. 2.15**, where:

$$\boldsymbol{C}_{\mathbf{P}} = C_{\mathrm{B}}$$

$$\boldsymbol{k}_{\mathrm{U}} = C_{\mathrm{B2}} \cdot k_{\mathrm{bi}}$$

(Similarly to **Section 2.1.4**, as the fraction of B2 consumed by the background pathway remains negligible, we treat the bimolecular B+B2 reaction as an effective first-order reaction with rate constant $C_{\mathrm{B2}} \cdot k_{\mathrm{bi}}$. We further assume that the first collisional extension $\mathrm{B} \rightarrow \mathrm{B}^{0}$ and the second collisional extension $\mathrm{B}^{0} \rightarrow \mathrm{B}^{*}$ share the same effective rate constant, as both reactions involve a free B2 encountering a similar extension site on B.)

In other words:

$$S_{\mathrm{BG}}(t) = C_{\mathrm{B}} [1 - (C_{\mathrm{B2}} \cdot k_{\mathrm{bi}} \cdot t + 1) e^{-C_{\mathrm{B2}} \cdot k_{\mathrm{bi}} \cdot t}] \quad \textbf{(2.18)}$$

According to definition of LOD in **Eq. 2.2**,

$$\mathrm{LOD}(t) = \frac{2\sigma}{1-\sigma} \cdot (K_{\mathrm{D}} + C_{\mathrm{B}}) \cdot \frac{K_{\mathrm{D2}} + C_{\mathrm{B2}}}{C_{\mathrm{B2}}} \cdot \frac{1 - (C_{\mathrm{B2}} \cdot k_{\mathrm{bi}} \cdot t + 1) e^{-C_{\mathrm{B2}} \cdot k_{\mathrm{bi}} \cdot t}}{1 - (k_{\mathrm{uni}} \cdot t + 1) e^{-k_{\mathrm{uni}} \cdot t}}$$

It can be proven that $\mathrm{LOD}(t)$ monotonically increases with respect to $t$ and reaches its lowest value as $t$ approaches zero:

$$\min(\mathrm{LOD}) = \lim_{t \to 0} \mathrm{LOD}(t) = \frac{2\sigma}{1-\sigma} \cdot (K_{\mathrm{D}} + C_{\mathrm{B}}) \cdot \frac{K_{D2} + C_{\mathrm{B2}}}{C_{\mathrm{B2}}} \cdot \left(\frac{C_{\mathrm{B2}}}{C'_{\mathrm{Local}}}\right)^{2}$$

Similar to the analyses before, the lower bound of LOD is achieved when $C_{\mathrm{B}}$, $C_{\mathrm{B2}}$ also approaches zero:

$$\min(\mathrm{LOD}) = \lim_{C_{\mathrm{B}}, C_{\mathrm{B2}}, t, \sigma \to 0} \mathrm{LOD}_{t} = 2\sigma \cdot K_{\mathrm{D}} \cdot \frac{K_{D2}}{C'_{\mathrm{Local}}} \cdot \frac{C_{\mathrm{B2}}}{C'_{\mathrm{Local}}} \quad \textbf{(2.19)}$$

Compare this with **Eq. 2.13**, it can be seen that the lower bound of LOD is further scaled down by a factor of $\frac{C_{\mathrm{B2}}}{C'_{\mathrm{Local}}}$. Although $C_{\mathrm{B2}}$ can be arbitrarily small, in practice it should be in the range of at least 100 to 200 pM to ensure proper yield and reaction time. Even so, this scaling factor can still be as low as 1/10,000 to 1/20,000 when $C'_{\mathrm{Local}}$ is on the order of 2 $\mu$M, and this is precisely the source of the 4-orders-of-magnitude improvement in sensitivity by SPEAR over PAG.

##### 2.1.7 LODs under realistic parameters

We start by experimentally measuring $C'_{\text{Local}}$. We set up 50 $\mu$L SPEAR reactions and took 2 $\mu$L aliquots at different time points and quenched with EDTA, then used qPCR to quantify the full record count (FRC). The experimentally measured FRC as a function of time was fitted to equation:

$$C = C_0 \cdot \left[1 - (1 + k_{\text{app}} \cdot t)e^{-k_{\text{app}} \cdot t}\right] \qquad \textbf{(2.20)}$$

to obtain that $k_{\text{app}}$ = 0.008 $s^{-1}$ for IFN$\gamma$ and 0.014 $s^{-1}$ for streptavidin (**Fig. S12**). Comparing equations **2.17** and **2.20**, we can see that $k_{\text{uni}} = k_{\text{app}}$. Thus, we can conclude that $k_{\text{uni}}$ is on the order of 0.01 $s^{-1}$ in SPEAR reactions, and we adopt this value as a representative $k_{\text{uni}}$ in the calculations below

To estimate $k_{\text{bi}}$ we can examine the data shown in **Fig. 1g**, where ~15 molecules (3.1 aM in the 8 µL reaction) of full records (B-B2*) were generated when the initial concentration of B and B2 were 100 pM and the reaction was carried out for 8 min (480 sec), using **Eq. 2.20** and the conditions $\frac{C}{C_0} = \frac{3.1\text{ aM}}{100\text{ pM}} = 3.1 \times 10^{-8}$ and $t = 480$ s, we can calculate that $k_{\text{app}}$ is roughly $5 \times 10^{-7}$ $s^{-1}$. Comparing equations **2.18** and **2.20** we can see that $k_{\text{bi}}$ equals $\frac{k_{\text{app}}}{C_{\text{B2}}}$, which is roughly 5,000 $M^{-1}s^{-1}$. We can then obtain $C'_{\text{Local}}$, which equals $\frac{k_{\text{uni}}}{k_{\text{bi}}} = \frac{0.01\text{ s}^{-1}}{5000\text{ M}^{-1}\text{s}^{-1}}$ and is on the order of 2 $\mu$M.

We now consider the kinetics of antibodies dissociating from the analyte, a process that was ignored in the analyses of **Sections 2.1.4** and **2.1.6**. To maintain clarity, we still disregard the rebinding of antibody after it dissociates from the analyte. This is largely valid especially considering that the immunocomplex is usually diluted during the amplicon generation reaction step. For example, in the workflow described in this manuscript, the antibody binding reaction is diluted by 4-fold; with $k_{\text{on}} \approx 5\times10^5$ $M^{-1}s^{-1}$ and [B] ≈ 100 pM after dilution, the rebinding probability over a 480-s extension window is $k_{\text{on}}\cdot[\text{B}]\cdot t \approx 2.4\%$. Even higher folds of dilution are usually done for PEA and PLA.

The differential equation set that describes the behavior of PAG assays should be modified to:

$$\frac{d[\text{A-B-B2}]_t}{dt} = -[\text{A-B-B2}]_t(k_{\text{uni}} + k_{\text{off1}} + k_{\text{off2}})$$

$$\frac{d[\text{A-B-B2}^*]_t}{dt} = [\text{A-B-B2}]_t \cdot k_{\text{uni}}$$

Solving this differential equation set yields:

$$S_{\text{True}}(t) = [\text{A-B-B2}^*]_t = \frac{[\text{A-B-B2}]|_{t=0} \cdot k_{\text{uni}}}{k_{\text{uni}} + k_{\text{off1}} + k_{\text{off2}}}\left[1 - e^{-(k_{\text{uni}}+k_{\text{off1}}+k_{\text{off2}})\cdot t}\right] \qquad \textbf{(2.21)}$$

where:

$$[\text{A-B-B2}]|_{t=0} = C_{\text{A}} \cdot \frac{C_{\text{B}}}{K_{\text{D}} + C_{\text{B}}} \cdot \frac{C_{\text{B2}}}{K_{\text{D2}} + C_{\text{B2}}}$$

By comparing **Eqs. 2.10** and **2.21**, we can see that dissociation of antibodies causes the plateau of $S_{\text{True}}$ (as $t$ approaches infinite) to be scaled down by a factor of $\frac{k_{\text{uni}}}{k_{\text{uni}}+k_{\text{off1}}+k_{\text{off2}}}$.

Similarly, LOD is scaled up by the inverse of this factor:

$$\mathrm{LOD}(t) = \frac{2\sigma}{1-\sigma} \cdot (K_{\mathrm{D}} + C_{\mathrm{B}}) \cdot \frac{1}{r_0 \cdot r_1} \cdot \frac{1 - e^{-C_{\mathrm{B2}} \cdot k_{\mathrm{bi}} \cdot t}}{1 - e^{-(k_{\mathrm{uni}} + k_{\mathrm{off1}} + k_{\mathrm{off2}}) \cdot t}} \tag{2.22}$$

where

$$r_0 = \frac{C_{\mathrm{B2}}}{K_{\mathrm{D2}} + C_{\mathrm{B2}}}$$

$$r_1 = \frac{k_{\mathrm{uni}}}{k_{\mathrm{uni}} + k_{\mathrm{off1}} + k_{\mathrm{off2}}}$$

This analysis highlights the importance of increasing $k_{\mathrm{uni}}$ (e.g., by using optimal concentration of the ligase or DNA polymerase in PLA and PEA, respectively) and using antibodies with low dissociation rate constants.

Similar adjustment can be made to the differential equation set that describes SPEAR:

$$\frac{d[\text{A-B-B2}]_t}{dt} = -[\text{A-B-B2}]_t \cdot (k_{\mathrm{uni}} + k_{\mathrm{off1}} + k_{\mathrm{off2}})$$

$$\frac{d[\text{A-B-B2}^0]_t}{dt} = [\text{A-B-B2}]_t \cdot k_{\mathrm{uni}} - [\text{A-B-B2}^0]_t \cdot (k_{\mathrm{uni}} + k_{\mathrm{off1}} + k_{\mathrm{off2}})$$

$$\frac{d[\text{A-B-B2}^*]_t}{dt} = [\text{A-B-B2}^0]_t \cdot k_{\mathrm{uni}}$$

This differential equation set has the following solution:

$$S_{\mathrm{True}}(t) = [\text{A-B-B2}^*]_t = [\text{A-B-B2}]|_{t=0} \cdot {r_1}^2 \cdot \left[1 - (1 + \lambda t) e^{-\lambda t}\right] \tag{2.22.1}$$

where

$$\lambda = k_{\mathrm{uni}} + k_{\mathrm{off1}} + k_{\mathrm{off2}}$$

$$r_1 = \frac{k_{\mathrm{uni}}}{\lambda}$$

$$[\text{A-B-B2}]_{t=0} = C_{\mathrm{A}} \cdot \frac{C_{\mathrm{B}}}{K_{\mathrm{D}} + C_{\mathrm{B}}} \cdot \frac{C_{\mathrm{B2}}}{K_{\mathrm{D2}} + C_{\mathrm{B2}}}$$

According to definition of LOD described in **Eq. 2.2**, while rewriting $k_{\mathrm{off1}}$ and $k_{\mathrm{off2}}$ as $k_{\mathrm{on}} \cdot K_{\mathrm{D}}$ and $k_{\mathrm{on2}} \cdot K_{\mathrm{D2}}$, respectively (to facilitate comparison with prior analyses such as those in **Fig. S10**), we can obtain:

$$\mathrm{LOD}(t) = \frac{2\sigma}{1-\sigma} (K_{\mathrm{D}} + C_{\mathrm{B}}) \frac{1}{r_0} \cdot \frac{1 - (C_{\mathrm{B2}} \cdot k_{\mathrm{bi}} \cdot t + 1) e^{-C_{\mathrm{B2}} \cdot k_{\mathrm{bi}} \cdot t}}{r_1^2 [1 - (1 + \lambda t) e^{-\lambda t}]} \tag{2.23}$$

where:

$$r_0 = \frac{C_{\mathrm{B2}}}{K_{\mathrm{D2}} + C_{\mathrm{B2}}}$$

$$\lambda = k_{\mathrm{uni}} + K_{\mathrm{D}} k_{\mathrm{on}} + K_{\mathrm{D2}} k_{\mathrm{on2}}$$

$$r_1 = \frac{k_{\mathrm{uni}}}{\lambda}$$

Finally, **Eqs. 2.4** (along with numerical adjustment to account for non-zero $E_{\text{in}}/E_{\text{act}}$), **2.22** and **2.23** define the LODs of allostery-based assays, conventional PAG assays and SPEAR, respectively, allowing us to conduct head-to-head comparison using realistic parameter sets. The signal-to-noise parameter $\sigma = 0.1$ corresponds to roughly 2× the typical assay CV (~5%) observed in our experiments. The on-rates $k_{\text{on}} = k_{\text{on2}} = 5\times10^5$ $M^{-1}s^{-1}$ are taken from typical antibody binding kinetics; the symmetric choice corresponds to a symmetric two-antibody system. In **Eqs. 2.22** and **2.23** the equilibrium occupancy terms are evaluated at the antibody concentrations of the binding step (400 pM in 2 μL), and the extension and background terms at the concentrations after addition of the Reaction Mix (100 pM in 8 μL), following the workflow in **Fig. 1f**; the LOD is expressed as the analyte concentration at the 8 μL stage, which is 1/4 of the value obtained by direct substitution into **Eqs. 2.22** and **2.23**. Using the parameter set below:

| Parameter | Value |
|---|---|
| $k_{\text{uni}}$ | 0.01 $s^{-1}$ |
| $k_{\text{bi}}$ | 5,000 $M^{-1}s^{-1}$ |
| $k_{\text{on}}, k_{\text{on2}}$ | $5\times10^5$ $M^{-1}s^{-1}$ |
| $C_{\text{B}}, C_{\text{B2}}$ | 100 pM (for PAG and SPEAR, this is the concentration in the 8 μL extension step; 400 pM in the 2 μL binding step) |
| $\sigma$ | 0.1 |
| $K_{\text{D}}$, $K_{\text{D2}}$ | Variable, $K_{\text{D}} = K_{\text{D2}}$ |
| $t$ | 8 min |
| $E_{\text{in}}/E_{\text{act}}$ | 0.01 |

we can compute the LODs for these three detection modalities (**Fig. S5**). We can see that the LOD of SPEAR is 3 to 4 orders of magnitude lower than PAG and can achieve below 100 molecules in 8 $\mu$L using antibodies with $K_{\text{D}}$ values in the range of 100 pM to 1 nM.

### 3 Supplementary Figures

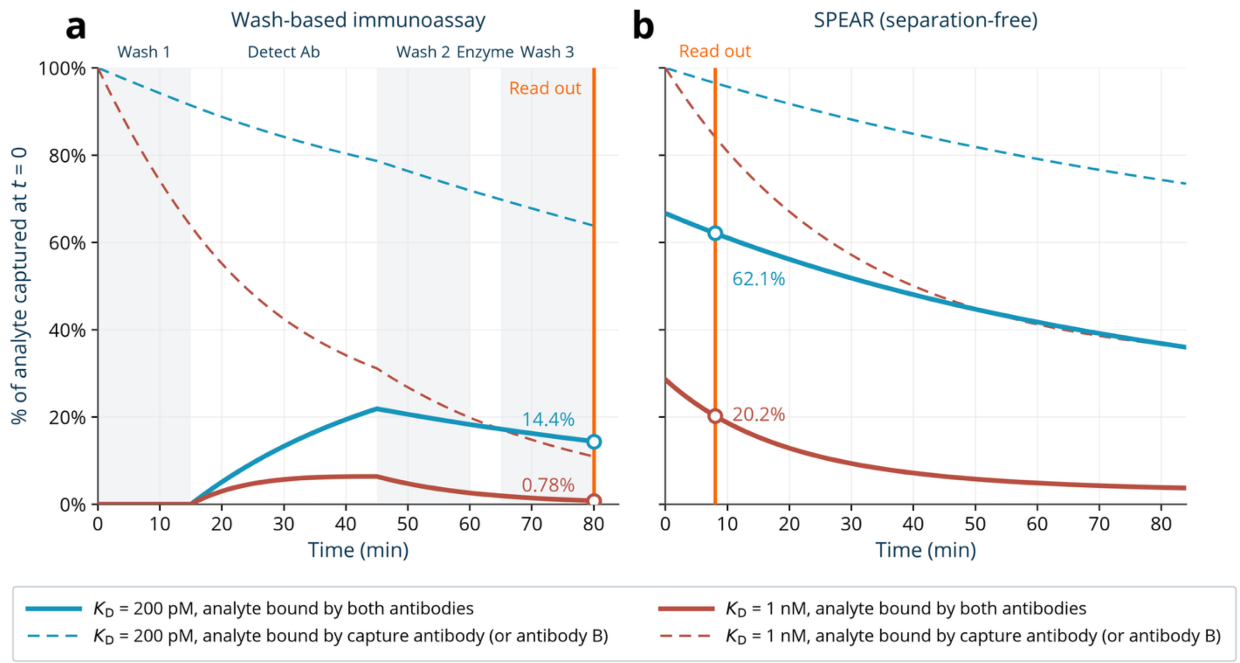

**Fig. S1. Simulation of signal loss due to spontaneous dissociation between analyte and antibody in wash-based immunoassays and in separation-free SPEAR assay. a**, In simulation of wash-based assays, we modeled the association and dissociation of surface-based binding the same way as solution-based binding and used 400 pM as the effective concentration of the capture antibody on the solid surface and 2 $\mu$L as the volume. The initial concentration of analyte was set to be 1 pM (although the result is independent of this concentration). For simplicity we assumed both antibodies have the same $k_{on}$ ($5\times10^5$ $M^{-1}s^{-1}$) and $k_{off}$. The simulation begins at the conclusion of the initial capture of the analyte by the surface-attached capture antibody, where the binding between analyte and capture antibody is assumed to reach equilibrium. The amount of analyte bound to the capture antibody at this equilibrium is defined as 100%. Next, during the 15-min wash (the 'Wash 1' step), the analyte is assumed to dissociate from the capture antibody with first-order rate constant $k_{off}$. The dissociated analyte is considered removed from the system (modeling washing with sufficiently large volume or continuous flow). Next, 400 pM detection antibody is added to the system for 30 min (the 'Detect Ab' step), during which the 'analyte to capture antibody' and 'analyte to detection antibody' bindings and dissociations are both allowed to happen in 2 $\mu$L volume and dissociated molecules are not removed from the system. In the next 15-min washing step (the 'Wash 2' step), dissociation between the analyte and the capture antibody and the dissociation between the analyte and the detection antibody are allowed to happen, and any species that does not contain the capture antibody is considered removed from the system. Next, during the 5-min addition of detection enzyme (such as HRP that binds to the biotinylated detection antibody, i.e., the 'Enzyme' step), binding and rebinding are both allowed to happen again in 2 $\mu$L volume. Lastly, another 15-min wash step (the 'Wash 3' step) is simulated similarly to previous wash steps. At the end of Wash 3, the fluorogenic substrate is added, and the amount of detection antibody still present in the system, which equals the amount of the analyte that is bound to both the capture and the detection antibodies, is quantified ('Read out'). Dashed lines show the amount of analyte that is bound to the capture antibody (with or without detection antibody), solid lines show the amount of analyte that is bound to both the capture and the detection antibodies, and open circle shows the latter at the moment of the 'Read out', all scaled to the 100% defined above. Simulations with two $K_D$ values (200 pM in teal, and 1 nM in terracotta) were performed, and $k_{off}$ values are derived from $K_D \cdot k_{on}$.

**b**, In the simulation of SPEAR assays, 400 pM of antibody B (conjugated with probe P), 400 pM of antibody B2 (conjugated with probe S), and 1 pM of analyte are assumed to reach binding equilibrium at 2 $\mu$L volume. The amount of analyte bound to the antibody B at this equilibrium is defined as 100%. Next, the reaction is diluted to 8 $\mu$L, and all species are allowed to re-establish equilibrium at the new volume over time according to $k_{on}$ and $k_{off}$ values. As SPEAR is a separation-free and wash-free assay, no species is considered removed from the system. $K_D$, $k_{on}$ and $k_{off}$ values were treated the same ways as in (**a**). Dashed lines show the amount of analyte that is bound to the antibody B (with or without antibody B2), solid lines show the amount of analyte that is bound to both antibody B and antibody B2, and open circles show the latter at the moment of the 'Read out', all scaled to the 100% defined above. Since SPEAR reaction reaches near completion within 8 min, we conservatively treat 8 min as the read-out moment although in reality, the SPEAR reaction happens continuously during the 8 min.

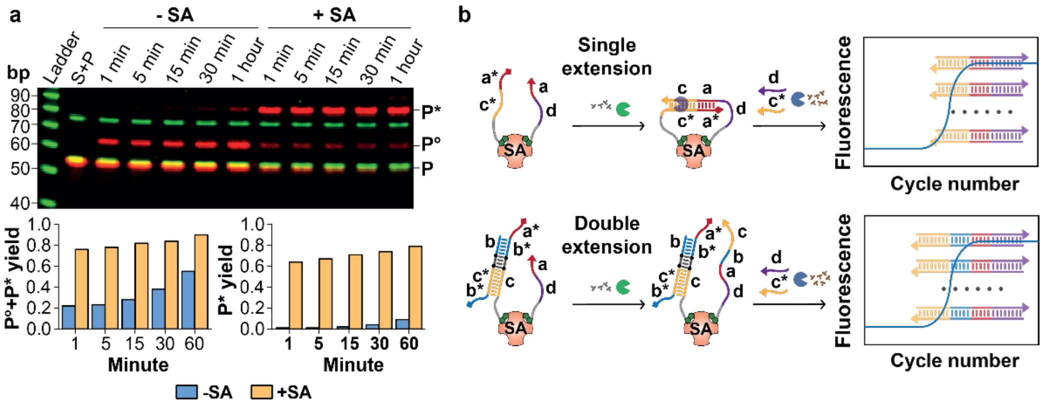

**Fig. S2. SPEAR proof-of-concept validation using streptavidin assay (SA). a,** Time-course analysis of SPEAR's two-step extension in the absence or presence of SA (25 nM) with biotinylated S and P probes (also Cy5 labelled, 50 nM). Reaction products were collected at 1, 5, 15, 30 and 60 minutes and visualized by SDS-PAGE. Red, Cy5 channel; green, SYBR Gold channel. Histograms show total yield of first and second extension products (bottom left) and yield of second extension product (bottom right) quantified from the Cy5 channel. Blue, without SA, and yellow, with SA. **b,** Schematic of sensitivity comparison between single and double extension. Biotinylated S probes with either a single toehold (a*) or double toeholds (a* and b*) were designed, requiring either a single extension (top) or double extensions (bottom) when in proximity with the P probe to generate a full record for qPCR amplification. a and a* indicate complementary sequences.

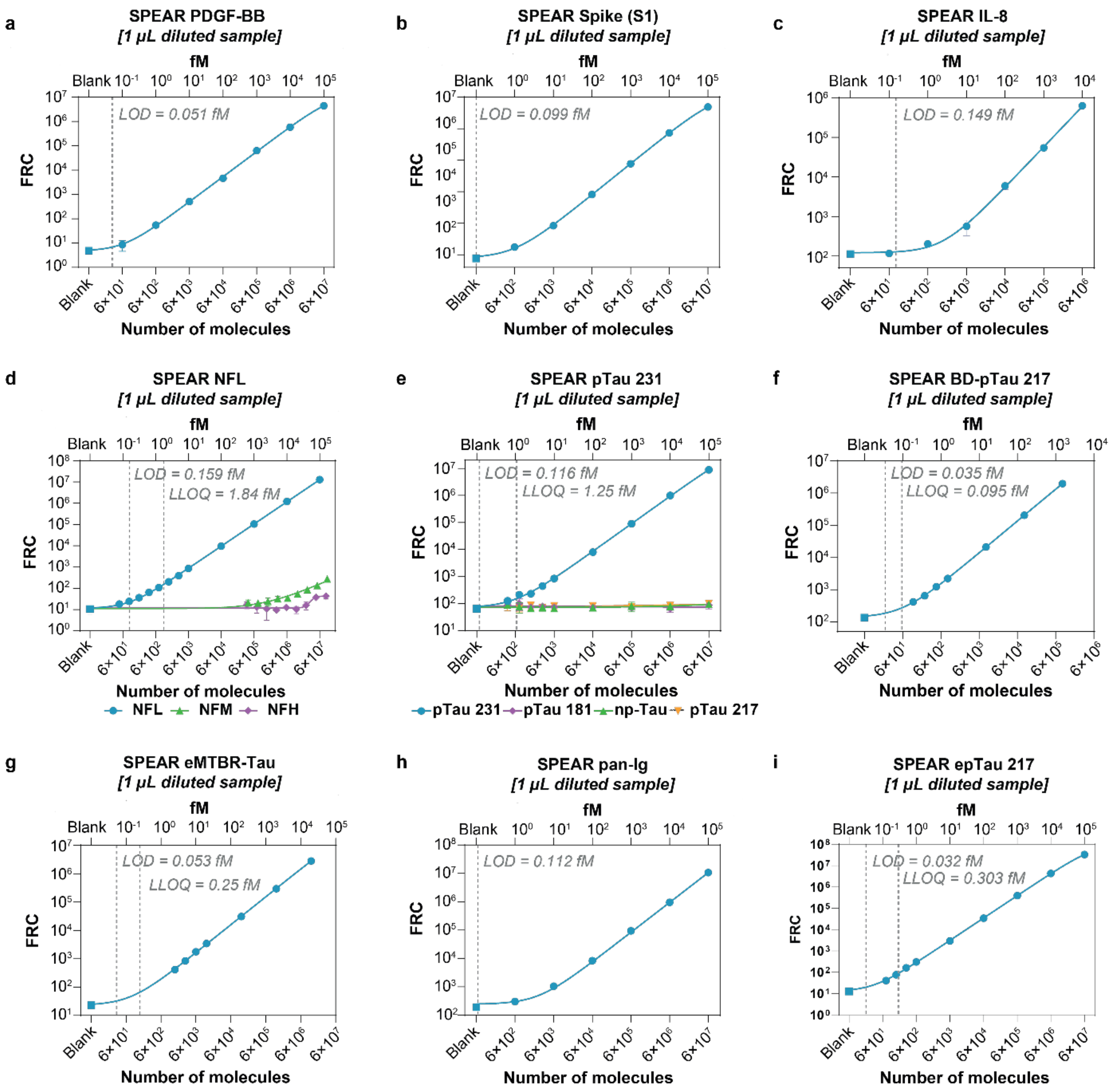

**Fig. S3. Calibration curves and analytical sensitivity of SPEAR immunoassays. a-i,** LODs for PDGF-BB, SARS-CoV-2 Spike (S1), IL-8, NFL, pTau 231, BD-pTau 217, eMTBR-Tau, pan-Ig, and epTau 217 were determined as $Mean_{blank}+2.5\times SD_{blank}$, with concentrations interpolated from a four-parameter logistic (4PL) fit. LLOQs were determined for the neurodegenerative disease assays (NFL, pTau 231, BD-pTau 217, and epTau 217) as the lowest interpolated concentrations meeting both ≤ 20% CV and 80 to 120% standard recovery criteria. **d, e,** The SPEAR NFL assay demonstrated minimal cross-reactivity with neurofilament medium (NFM) and heavy (NFH) chains, while the SPEAR pTau 231 assay showed specificity for pTau 231, with negligible cross-reactivity with pTau 181, pTau 217, and non-phosphorylated tau.

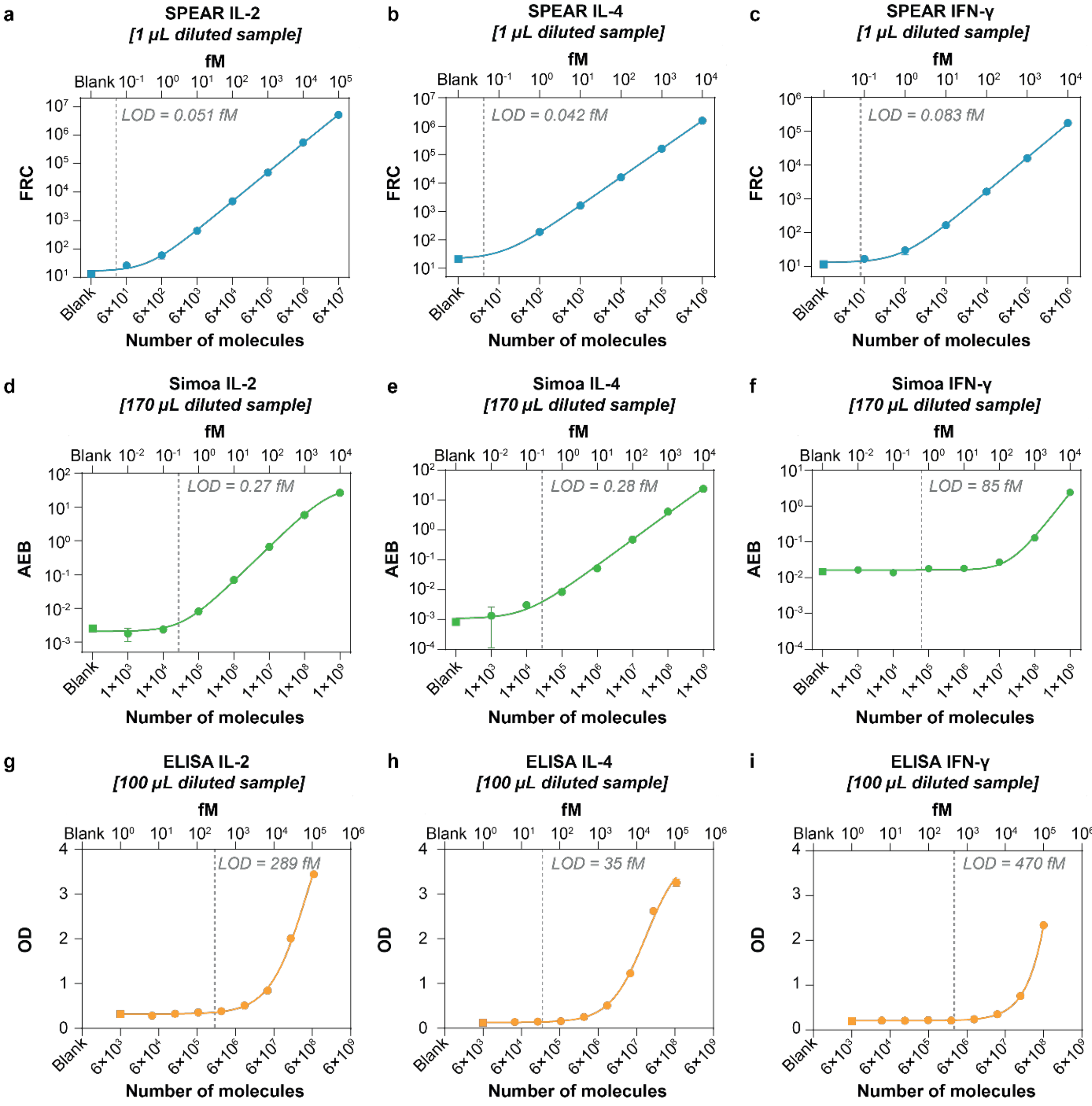

**Fig. S4. Calibration curves for LOD estimation for SPEAR, dELISA (Simoa) and conventional ELISA for assays using identical antibodies.** LOD was determined for SPEAR (**a-c**), Simoa (**d-f**), and ELISA (**g-i**) using identical antibody clones for human IL-2, IL-4 and IFN-γ detection. LOD values were calculated as $Mean_{blank}$+2.5×$SD_{blank}$ and interpolated from a four-parameter logistic fit of the calibration curves.

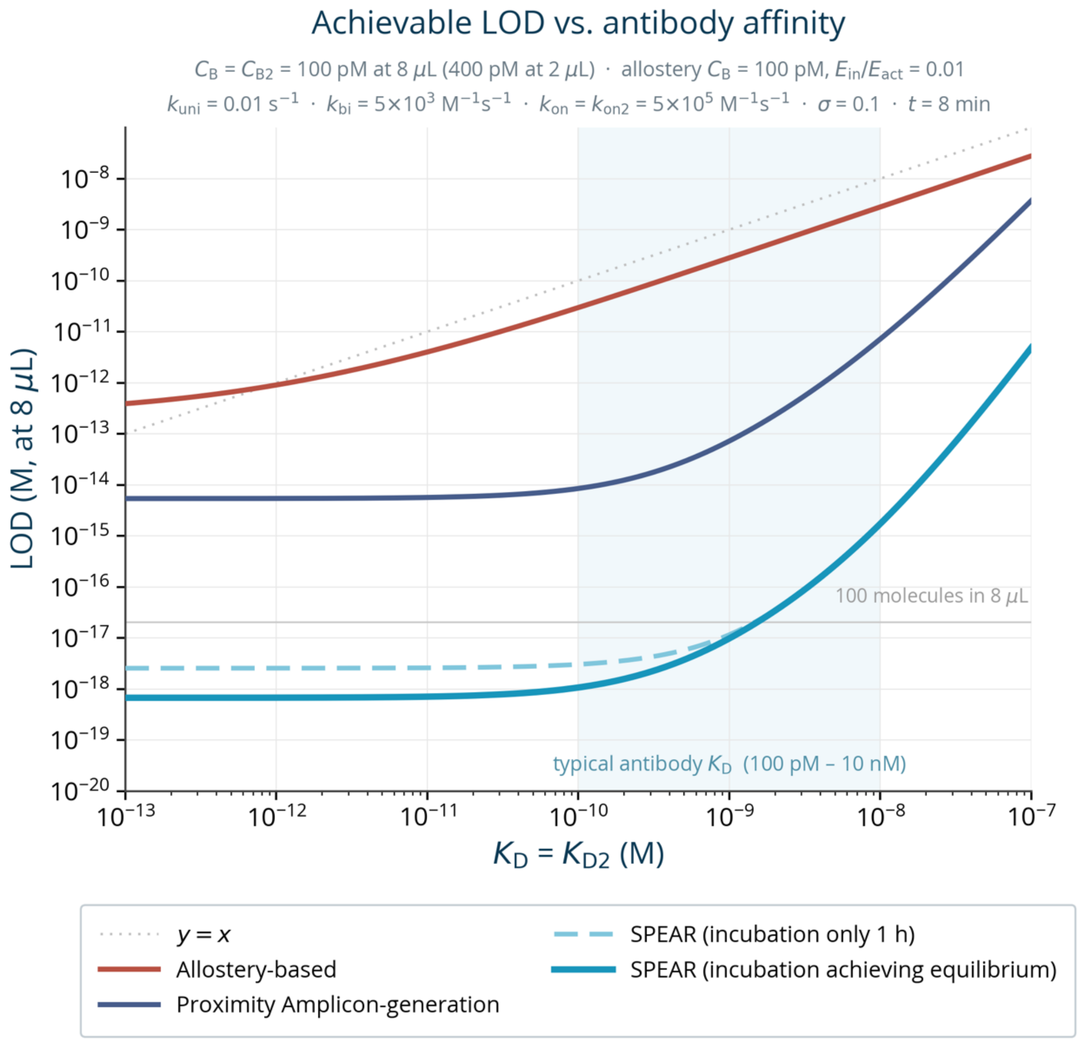

**Fig. S5. Theoretically achievable LODs under realistic parameters.** Theoretically achievable LODs for allosteric signaling binders (terracotta) as functions of $K_D$ of the binder in the active state are reproduced from **Fig. S10f** with binder concentration of 100 pM and $E_{in}/E_{act}$ ratio of 0.01 (see **Section 2.1.3** for details). Theoretically achievable LODs for conventional proximity amplicon-generation assays (dark blue) and SPEAR (teal, solid line) as functions of $K_D$ of the antibodies used are calculated using **Eqs. 2.22** and **2.23**, respectively. For SPEAR, LODs are also calculated through simulating the actual workflow shown in **Fig. 1f**. The LOD is reported as the concentration in the 8 $\mu$L stage (after addition of Reaction Mix, before the addition of qPCR Mix, light teal dashed line). This corresponds to 1/8 of the LOD reported for the developed SPEAR assays shown in **Fig. S3** and **Fig. 1h**. For these simulations, the initial incubation of the two antibodies and the sample is modeled to occur at 4× concentration (for all species, as the volume is 2 $\mu$L) for 1 hour and the primer extension reactions are modeled to initiate after the addition of the 6 $\mu$L Reaction Mix which dilutes the concentration of every species to 1×.

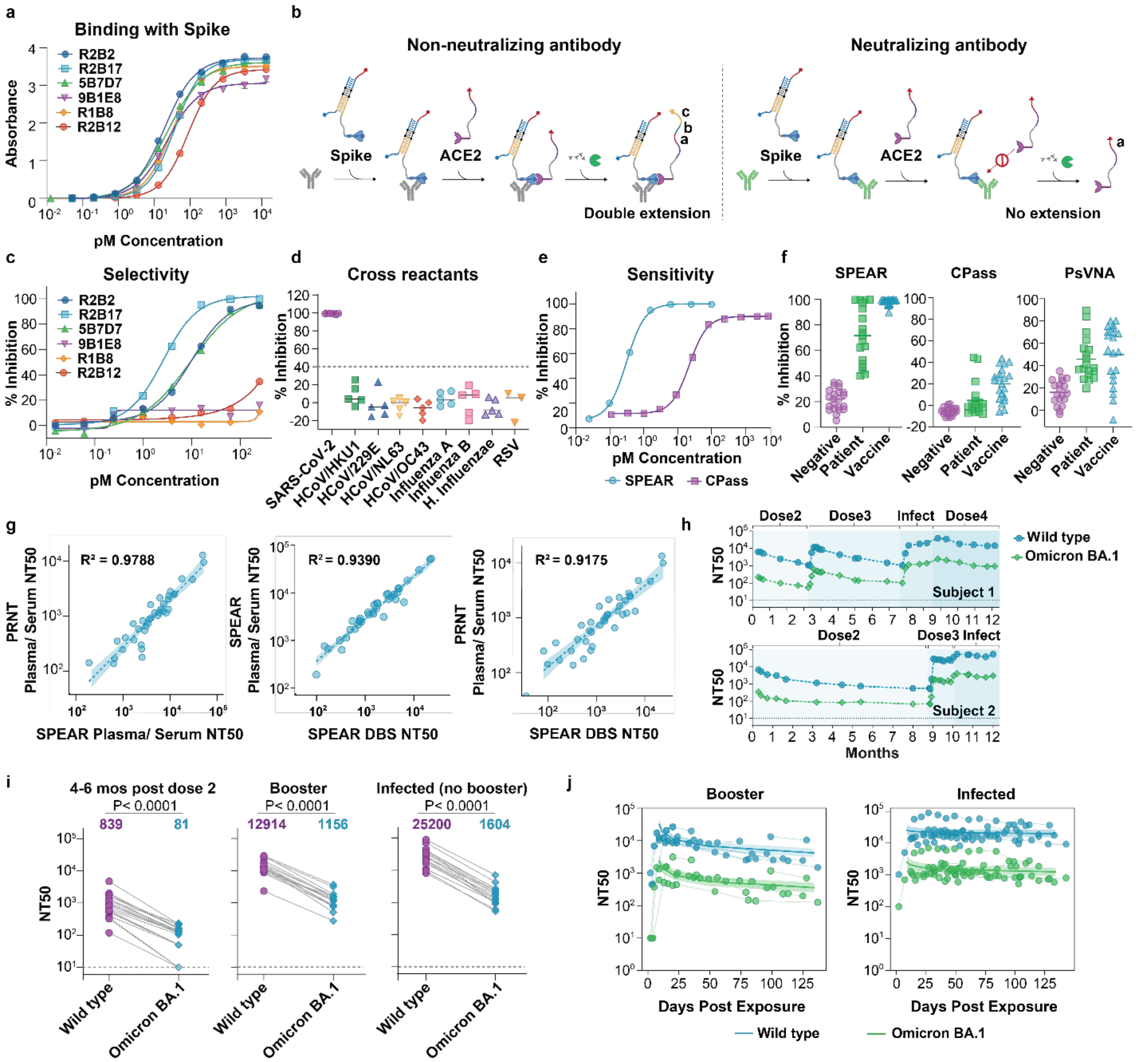

**Fig. S6. Performance of SPEAR SARS-CoV-2 Neutralization Assay. a,** Six anti-Spike monoclonal IgGs with varying reported neutralizing activities were evaluated by ELISA for binding to Spike protein. R2B17 was selected as the calibrator for the pan-Ig assay. **b,** Reaction schematic of the SPEAR signal-off SARS-CoV-2 Neutralization Assay. SPEAR probes are conjugated to Spike protein and human ACE2. Neutralizing antibodies compete with ACE2 for binding to Spike, preventing proximity-driven extension (right), whereas non-neutralizing antibodies allow ACE2- and Spike-conjugated probes to come into proximity and undergo extension (left). Thus, signal inhibition reflects neutralizing antibody levels. **c,** The six anti-Spike monoclonal IgGs shown in **a** were evaluated for neutralizing activity in the presence of ACE2. **d,** Specificity analysis using serum samples from SARS-CoV-2 vaccine recipients and convalescent individuals who had recovered from the indicated viral infections. HCoV/HKU1, human coronavirus HKU1; HCoV/229E, human coronavirus 229E; HCoV/NL63, human coronavirus NL63; HCoV/OC43, human coronavirus OC43; H. influenzae, Haemophilus influenzae; RSV, respiratory syncytial virus. **e,** Comparison of achievable sensitivity between SPEAR and CPass$^{TM}$ using the monoclonal antibody R2B17. **f,** Sensitivity comparison of SPEAR, CPass$^{TM}$, and the pseudovirus neutralization assay (PsVNA)

using DBS samples from negative individuals (purple, *n* = 22), COVID-19 patients (green, *n* = 19), and vaccine recipients (blue, *n* = 20). Sample volumes were 1 µL for SPEAR, 60 µL for CPass™, and 20 µL for PsVNA. Bars indicate mean values. **g,** Concordance of neutralization titers in vaccine donors (*n* = 35). Left: SPEAR versus cell-based PRNT measurements in plasma/serum; middle: SPEAR measurements in DBS versus plasma/serum; right: SPEAR measurements in DBS versus PRNT measurements in plasma/serum. Pearson $R^2$ values are shown. Dotted lines indicate simple linear regression, and shaded regions indicate 95% confidence intervals. **h,** Longitudinal monitoring of DBS neutralizing antibody titers in two individuals with different exposure histories against wild-type (blue) and Omicron BA.1 (green) variants. Exposure and vaccination events are indicated along the timelines. **i,** Neutralizing titers in vaccinated subjects four to six months after the second dose (left, *n* = 21), subjects who received a third-dose Moderna or Pfizer booster (middle, *n* = 18), and subjects who received a two-dose primary vaccination series followed by infection during the Omicron wave beginning in December 2021 without receiving a third-dose booster (right, *n* = 18). DBS samples were evaluated against wild-type (Wuhan-Hu-1) and Omicron BA.1. Geometric mean titers for wild-type (purple) and Omicron BA.1 (cyan) are shown. P values were calculated using unpaired t-tests; dotted lines indicate the assay detection limit. **j,** Longitudinal monitoring of neutralizing antibody waning against wild-type (blue) and Omicron BA.1 (green) in a larger cohort. Left: subjects who received two primary doses and one booster (*n* = 7). Right: individuals infected during the Omicron wave beginning in December 2021 (*n* = 16). Lines connect measurements from the same individual; solid lines indicate exponential decay fits, and shaded regions indicate 95% confidence intervals.

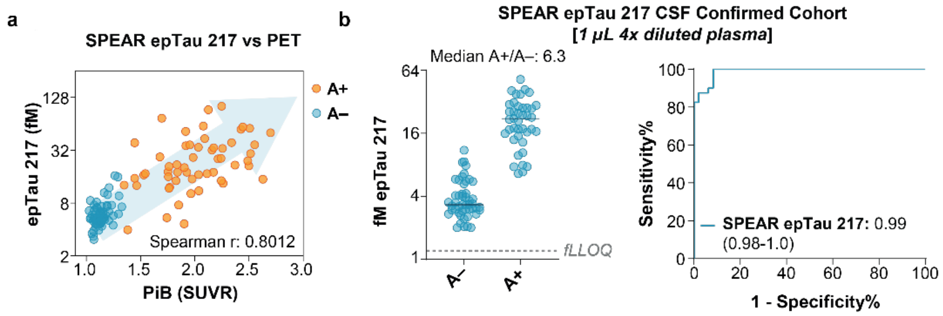

**Fig. S7. Clinical performance of SPEAR epTau 217 Assay for amyloid PET confirmation a,** Correlation of amyloid PiB standardized uptake value ratio (SUVR) and SPEAR epTau 217 measurement in the Pitt-ADRC cohort. Legends indicate Spearman correlation coefficient. Blue dots represent amyloid PET-negative individuals; orange dots represent amyloid PET-positive individuals. **b,** Clinical differentiation of amyloid status confirmed by CSF (A−: $n = 48$; A+: $n = 40$) using SPEAR epTau 217 assay with median A+/A− indicated (left) and corresponding Receiver Operating Characteristic (ROC) analysis of A+ vs. A− (right). Legends show Area under curve (AUC) of A+ vs. A− with 95% confidence interval. Bars show medians; grey dashed line: fLLOQ.

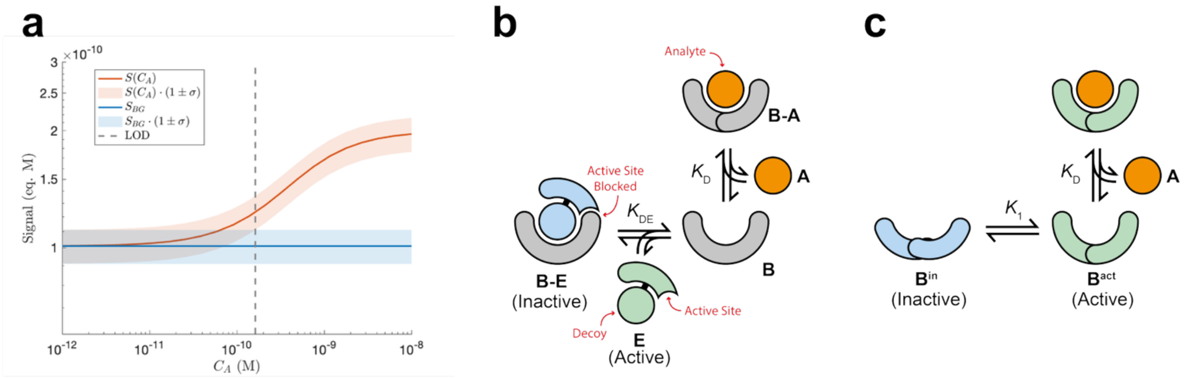

**Fig. S8. General concepts and schemes. a,** Definition of limit of detection (LOD). The mean signal and background as a function of analyte concentration ($C_{\mathrm{A}}$) are shown in red and blue lines, while their noises are shown in shades of the same color. The LOD is defined as the analyte concentration at which the two shades separate. **b,** Schematic of competition-based protein-detection assays. **c,** Schematic of allostery-based protein-detection assays.

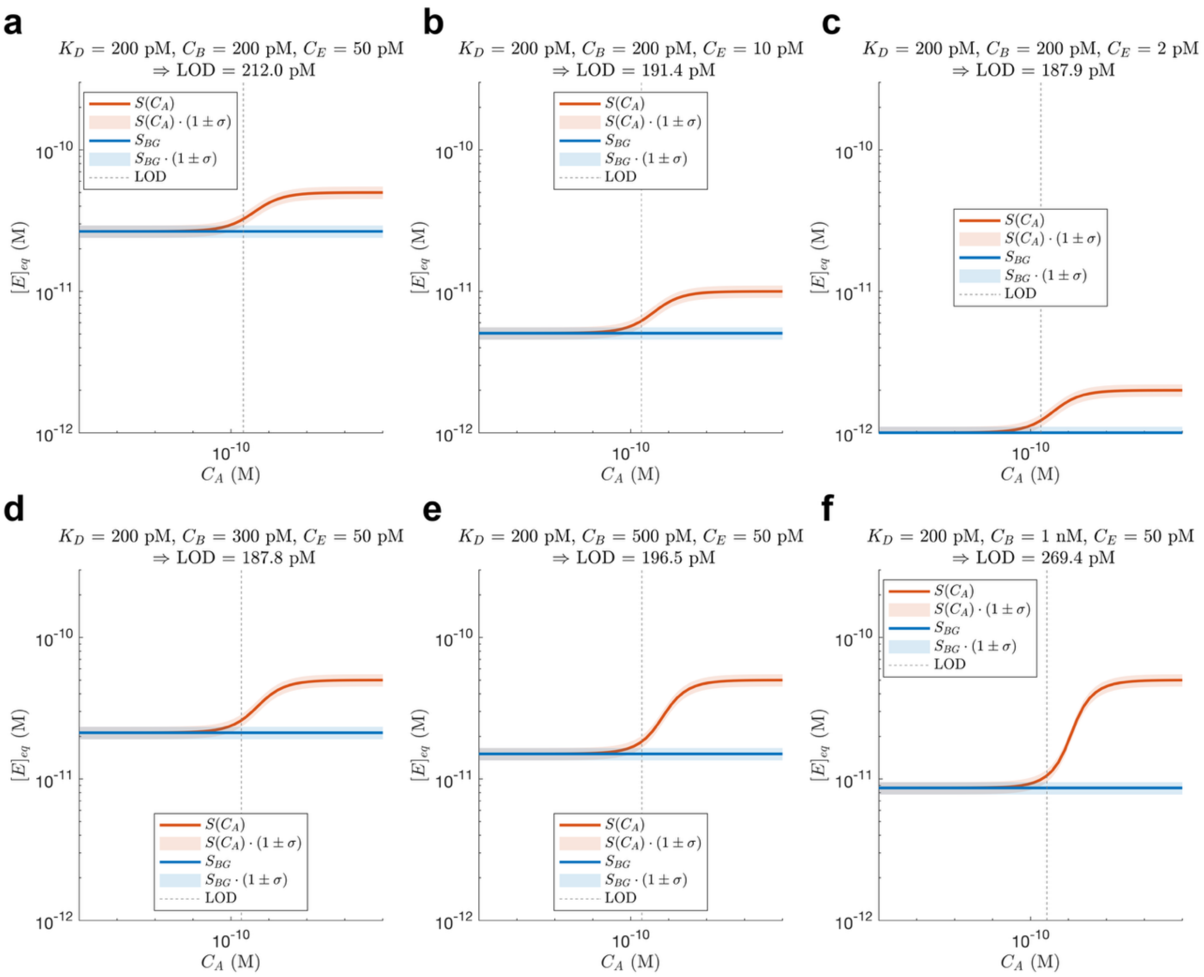

**Fig. S9. Theoretically achievable LOD in competition-based protein-detection assays as a function of the concentrations of the signal-generating molecule (CE) and binder (CB).** The parameters used are shown on top of each panel.

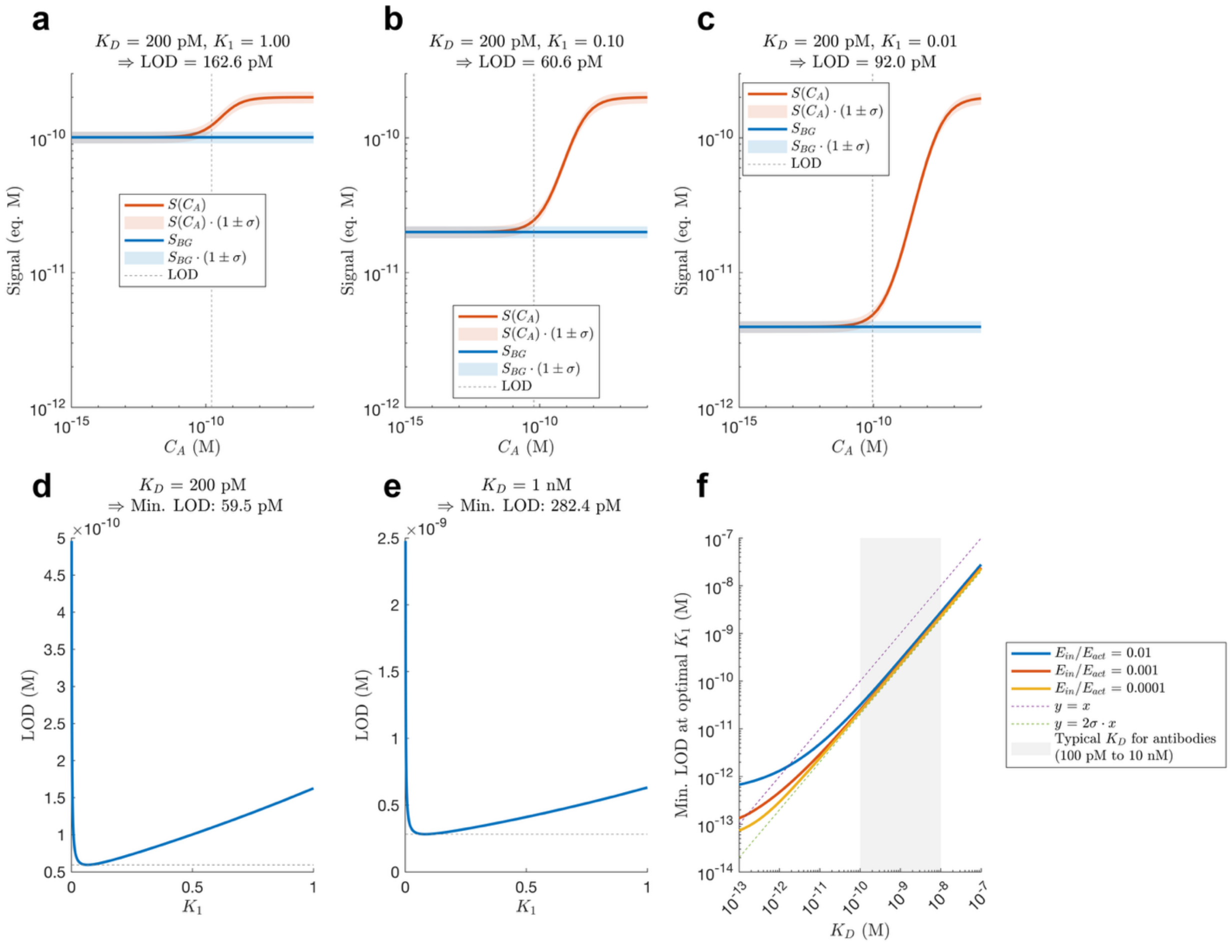

**Fig. S10. LOD of allostery-based binders under realistic parameters. a-c,** Signal, background and their variations $(1 \pm \sigma)$ as a function of analyte concentration $C_\mathrm{A}$, for different $K_1$ values. The lowest $C_\mathrm{A}$ at which the red shade separates from the blue shade is the definition of LOD. **d, e**, Optimum $K_1$ and their corresponding LOD for $K_\mathrm{D}$ of 200 pM and 1 nM. **f**, Lowest achievable LOD as a function of $K_\mathrm{D}$ for different $E_\mathrm{in}/E_\mathrm{act}$ values.

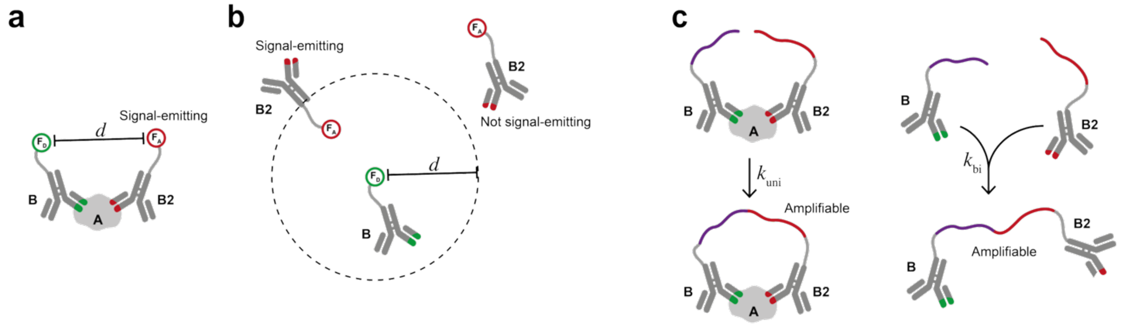

**Fig. S11. Abstract models for proximity-based immunoassays. a**, **b**, A general model for the generation of true signal (**a**) and background signal (**b**) in an ideal FRET-like system. **c**, A general model for the generation of true signal (left) and background signal (right) in conventional proximity amplicon-generation assays.

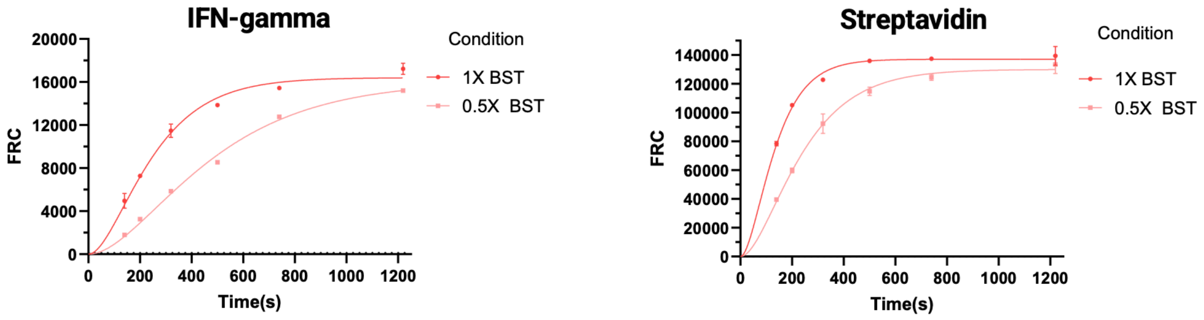

**Fig. S12. Kinetics of full record formation in SPEAR assays to detect IFN-$\gamma$ (left) and streptavidin (right).**

#### 4 Supplementary Tables

**Supplementary Tables S1** to **S5** are provided as a separate Excel file.

**Table S6. Participant demographics for capillary and Tasso concordance study from Memory/Cognitive Disorders clinic at Banner Sun Health Research Institute**

| Characteristic | Value |
| --- | --- |
| **Age (years)** | 80.0 ± 6.9 |
| **Female (n (%))** | 8 (47.1) |
| **CU (n (%))** | 8 (47.1) |
| **MCI (n (%))** | 4 (23.5) |
| **AD/Dementia (n (%))** | 5 (29.4) |

**Table S7. Participant demographics for the Bio-Hermes-002 cohort.**

| | Total | | A–/T– | | A+/T– | | A+/T+ | | p-value |
|---|---|---|---|---|---|---|---|---|---|
| **N (%)** | 127 | | 33 | (26.0%) | 37 | (29.1%) | 57 | (44.9%) | |
| **Age (SD), years** | 73.7 | (7.3) | 69.6 | (6.4) | 73.1 | (7.8) | 76.4 | (6.2) | <0.0001 |
| **Sex** | | | | | | | | | 0.0442 |
| Female (%) | 80 | (63.0%) | 26 | (78.8%) | 24 | (64.9%) | 30 | (52.6%) | |
| Male (%) | 47 | (37.0%) | 7 | (21.2%) | 13 | (35.1%) | 27 | (47.4%) | |
| **Race** | | | | | | | | | 0.200 |
| Non-hispanic White (%) | 117 | (92.1%) | 30 | (90.9%) | 32 | (86.5%) | 55 | (96.5%) | |
| Black/ African American (%) | 7 | (5.5%) | 3 | (9.1%) | 3 | (8.1%) | 1 | (1.8%) | |
| Others | 3 | (2.4%) | 0 | (0.0%) | 2 | (5.4%) | 1 | (1.8%) | |
| **MMSE (SD)** | 25.7 | (4.1) | 29.4 | (1.1) | 26.7 | (2.8) | 22.8 | (3.8) | <0.0001 |
| **Clinical diagnosis** | | | | | | | | | <0.0001 |
| Cognitive Normal (%) | 44 | (34.6%) | 33 | (100.0%) | 11 | (29.7%) | 0 | (0.0%) | |
| MCI (%) | 41 | (32.3%) | 0 | (0.0%) | 16 | (43.2%) | 25 | (43.9%) | |
| Probable AD (%) | 42 | (33.1%) | 0 | (0.0%) | 10 | (27.0%) | 32 | (56.1%) | |
| **Years of Education** | 16.0 | (2.7) | 16.8 | (1.8) | 15.9 | (2.9) | 15.5 | (2.9) | 0.1245 |

Note: Mean and standard deviation (SD) are reported for continuous variables; and frequences and percentages for categorical variables. p-values among amyloid PET negative/Tau PET negative (A–/T–), amyloid PET positive/Tau PET negative (A+/T–), and amyloid PET positive/Tau PET positive (A+/T+) groups were calculated using Kruskal–Wallis test and Fisher's exact test for categorical variables, respectively. MMSE: The Mini Mental State Examination; MCI: mild cognitive impairment

**Table S8. Participant demographics for the Pitt-ADRC cohort.**

| | Total | | A– | | A+ | | p-value |
|---|---|---|---|---|---|---|---|
| **N (%)** | 119 | | 63 | (52.9%) | 56 | (47.1%) | |
| **Age (SD), years** | 69.3 | (7.8) | 67.5 | (7.4) | 70.9 | (7.8) | 0.004 |
| **Sex** | | | | | | | 0.044 |
| Female (%) | 52 | (43.7%) | 22 | (34.9%) | 30 | (53.6%) | |
| Male (%) | 67 | (56.3%) | 41 | (65.1%) | 26 | (46.4%) | |
| **Education (SD), years** | 16.6 | (2.7) | 16.5 | (2.7) | 16.7 | (2.7) | 0.716 |
| **Race** | | | | | | | 0.017 |
| Non-hispanic White (%) | 111 | (93.3%) | 57 | (90.5%) | 54 | (96.4%) | |
| Black/ African American (%) | 4 | (3.4%) | 4 | (6.3%) | 0 | (0.0%) | |
| Others | 2 | (1.7%) | 0 | (0.0%) | 2 | (3.6%) | |
| **MMSE (SD)** | 23.9 | (7.8) | 25.9 | (7.2) | 21.7 | (8.0) | <0.001 |
| **Clinical diagnosis** | | | | | | | <0.001 |
| Normal (%) | 33 | (27.7%) | 25 | (39.7%) | 8 | (14.3%) | |
| MCI (%) | 41 | (34.5%) | 22 | (34.9%) | 19 | (33.9%) | |
| AD dementia (%) | 25 | (21.0%) | 4 | (6.3%) | 21 | (37.5%) | |
| Non-AD dementia (%) | 18 | (15.1%) | 10 | (15.9%) | 8 | (14.3%) | |
| **APOE ε4 carrier** | | | | | | | 0.079 |
| No (%) | 77 | (64.7%) | 45 | (71.4%) | 32 | (57.1%) | |
| Yes (%) | 40 | (33.6%) | 16 | (25.4%) | 24 | (42.9%) | |
| **PiB SUVR (SD)** | 1.54 | (0.50) | 1.13 | (0.07) | 2.01 | (0.34) | <0.001 |
| **AV1451 SUVR (SD)** | 1.31 | (0.33) | 1.14 | (0.07) | 1.50 | (0.39) | <0.001 |
| **Time of blood-draw before PET, years** | 2.40 | (3.7) | 2.99 | (4.4) | 1.74 | (2.6) | 0.050 |
| **N Status** | | | | | | | 0.003 |
| Negative (%) | 64 | (53.8%) | 42 | (66.7%) | 22 | (39.3%) | |
| Positive (%) | 51 | (42.9%) | 19 | (30.2%) | 32 | (57.1%) | |
| **Cortical thickness (SD)** | 2.70 | (0.15) | 2.75 | (0.14) | 2.64 | (0.15) | <0.001 |

Note: Mean and standard deviation (SD) are reported for continuous variables; and frequences and percentages for categorical variables. p-values between amyloid PET negative (A–) and positive (A+) groups were calculated using Wilcoxon rank-sum test and Fisher's exact test for categorical variables, respectively. MMSE: The Mini Mental State Examination; MCI: mild cognitive impairment; SUVR: Standardized uptake value ratio.

**Table S9. Participant demographics of the CSF AD biomarker confirmed cohort from MGH.**

| CSF Confirmed | Total | | A– | | A+ | | p-value |
|---|---|---|---|---|---|---|---|
| **N (%)** | 88 | | 48 | (54.5%) | 40 | (45.5%) | |
| **Age (SD), years** | 67.3 | (9.4) | 65.8 | (8.5) | 69.2 | (10.1) | 0.129 |
| **Sex** | | | | | | | 0.381 |
| Female (%) | 33 | (37.5%) | 20 | (41.7%) | 13 | (32.5%) | |
| Male (%) | 54 | (61.4%) | 27 | (56.3%) | 27 | (67.5%) | |
| **Clinical diagnosis** | | | | | | | <0.001 |
| CU (%) | 25 | (28.4%) | 25 | (52.1%) | 0 | (0.0%) | |
| MCI (%) | 38 | (43.2%) | 20 | (41.7%) | 18 | (45.0%) | |
| Dementia (%) | 24 | (27.3%) | 2 | (4.2%) | 22 | (55.0%) | |

Note: Mean and standard deviation (SD) are reported for continuous variables; and frequences and percentages for categorical variables. p-values between amyloid negative (A–) and positive (A+) groups were calculated using Wilcoxon rank-sum test and Fisher's exact test for categorical variables, respectively. CU: cognitive unimpaired; MCI: mild cognitive impairment.

**Table S10. Participant demographics for the Human Connectome Project (HCP) cohort.**

| | Total | | A– | | A+ | | p-value |
|---|---|---|---|---|---|---|---|
| **N (%)** | 113 | | 85 | (75.2%) | 28 | (24.8%) | |
| **Age (SD), years** | 66.4 | (9.0) | 64.1 | (8.7) | 73.3 | (5.9) | <0.001 |
| **Sex** | | | | | | | 0.016 |
| Female (%) | 67 | (59.3%) | 56 | (65.9%) | 11 | (39.3%) | |
| Male (%) | 46 | (40.7%) | 29 | (34.1%) | 17 | (60.7%) | |
| **Education (SD), years** | 15.4 | (3.1) | 14.9 | (2.8) | 16.9 | (3.6) | 0.005 |
| **Race** | | | | | | | 0.014 |
| Non-Hispanic White (%) | 64 | (56.6%) | 42 | (49.4%) | 22 | (78.6%) | |
| Black/ African American (%) | 48 | (42.5%) | 42 | (49.4%) | 6 | (21.4%) | |
| **MOCA (SD)** | 24.9 | (3.5) | 24.9 | (3.3) | 25.0 | (4.0) | 0.587 |
| **Clinical diagnosis** | | | | | | | 0.004 |
| Normal (%) | 80 | (70.8%) | 64 | (75.3%) | 16 | (57.1%) | |
| MCI (%) | 29 | (25.7%) | 20 | (23.5%) | 9 | (32.1%) | |
| AD (%) | 4 | (3.5%) | 1 | (1.2%) | 3 | (10.7%) | |
| **APOE ε4 carrier** | | | | | | | 0.356 |
| No (%) | 77 | (68.1%) | 60 | (70.6%) | 17 | (60.7%) | |
| Yes (%) | 36 | (31.9%) | 25 | (29.4%) | 11 | (39.3%) | |
| **PiB SUVR (SD)** | 1.26 | (0.33) | 1.10 | (0.06) | 1.75 | (0.31) | <0.001 |
| **Time of blood-draw before PET, days** | 19.5 | (120.1) | 23.6 | (109.9) | 7.1 | (148.5) | 0.608 |
| **N Status** | | | | | | | 0.266 |
| Negative (%) | 71 | (62.8%) | 56 | (65.9%) | 15 | (53.6%) | |
| Positive (%) | 42 | (37.2%) | 29 | (34.1%) | 13 | (46.4%) | |
| **Cortical thickness (SD)** | 2.71 | (0.11) | 2.73 | (0.10) | 2.66 | (0.15) | 0.042 |

Note: Mean and standard deviation (SD) are reported for continuous variables; and frequences and percentages for categorical variables. p-values between amyloid PET negative (A–) and positive (A+) groups were calculated using Wilcoxon rank-sum test and Fisher's exact test for categorical variables, respectively. MoCA: Montreal Cognitive Assessment; MCI: mild cognitive impairment; SUVR: Standardized uptake value ratio.
